# Neuronal lipid droplets play a conserved and sex-biased role in maintaining whole-body energy homeostasis

**DOI:** 10.1101/2024.09.19.613929

**Authors:** Romane Manceau, Danie Majeur, Celena M. Cherian, Colin J. Miller, Lianna W. Wat, Jasper D. Fisher, Audrey Labarre, Serena Hollman, Sanjana Prakash, Sébastien Audet, Charlotte F. Chao, Lewis Depaauw-Holt, Benjamin Rogers, Anthony Bosson, Joyce J.Y. Xi, Catrina A.S. Callow, Niyoosha Yoosefi, Niki Shahraki, Yi Han Xia, Alisa Hui, Jared VanderZwaag, Khalil Bouyakdan, Demetra Rodaros, Pavel Kotchetkov, Caroline Daneault, Ghazal Fallahpour, Martine Tetreault, Marie-Ève Tremblay, Matthieu Ruiz, Baptiste Lacoste, J.A. Parker, Ciaran Murphy-Royal, Tao Huan, Stephanie Fulton, Elizabeth J. Rideout, Thierry Alquier

## Abstract

Lipids are essential for neuron development and physiology. Yet, the central hubs that coordinate lipid supply and demand in neurons remain unclear. Here, we combine invertebrate and vertebrate models to establish the presence and functional significance of neuronal lipid droplets (LD) *in vivo*. We find that LD are normally present in neurons in a non-uniform distribution across the brain, and demonstrate triglyceride metabolism enzymes and lipid droplet-associated proteins control neuronal LD formation through both canonical and recently-discovered pathways. Appropriate LD regulation in neurons has conserved and male-biased effects on whole-body energy homeostasis across flies and mice, specifically neurons that couple environmental cues with energy homeostasis. Mechanistically, LD-derived lipids support neuron function by providing phospholipids to sustain mitochondrial and endoplasmic reticulum homeostasis. Together, our work identifies a conserved role for LD as the organelle that coordinates lipid management in neurons, with implications for our understanding of mechanisms that preserve neuronal lipid homeostasis and function in health and disease.

**HIGHLIGHTS:**

- Lipid droplets (LD) normally form in neurons across species Neuronal LD are regulated by a conserved gene network
- Neuronal LD regulation plays a conserved and sex-biased role in maintaining energy homeostasis
- LD regulation supports ER and mitochondrial function in hunger-activated neurons

**GRAPHICAL ABSTRACT:** 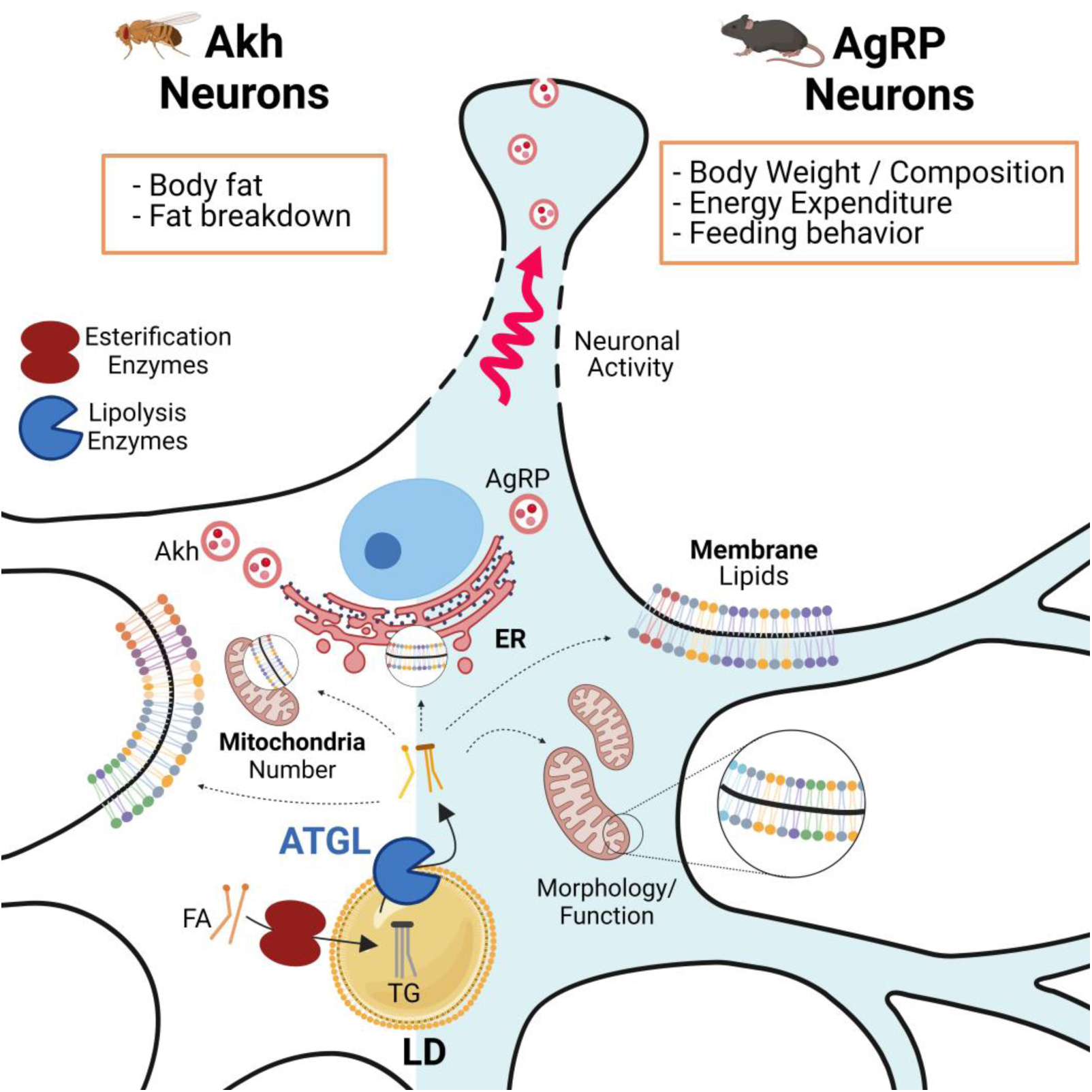

## MAIN

Lipids are essential for neuron development and function^1–3^. Indeed, a rich body of literature shows that phospholipids, glycerolipids, fatty acids (FA), and neutral lipids support diverse aspects of neuron physiology and function^3–6^. Yet, the hubs that coordinate neuronal lipid metabolism to maintain neuron function remain unclear. In non-neuronal cells, lipid droplets (LD) manage this tight coupling between lipid supply and demand^7–9^. LD are dynamic lipid-storing organelles with a phospholipid monolayer surrounding a neutral lipid core comprised of triacylglycerol (TG) and cholesterol esters^7–9^. LD are found across species in diverse non- neuronal cell types in both physiological conditions and during stress^8^. Despite evidence that LD form in cultured neurons^10–14^ and accumulate in neurons in pathological contexts and in disease models^15–23^, the prevailing view is that LD are not normally present in neurons *in vivo*.

Clues into LD regulation and function emerge from 20 years of studies on LD in non- neuronal cells across diverse species^7,8^. Many LD-associated proteins have been identified, including enzymes that control neutral lipid synthesis and breakdown, and factors that affect LD formation, maintenance, and transport^7,8^. These LD-associated proteins adjust LD dynamics to match lipid supply with demand, mechanisms that are well-conserved across eukaryotes^7,8^. LD were initially described as an energy storage organelle; however, LD are now known to play broad roles in supporting membrane homeostasis, cell signaling, and physiology^7–9^. For example, appropriate LD regulation under normal physiological conditions provides substrates for energy production via β oxidation, ligands to mediate cell signaling, and precursors to maintain plasma membrane and organelle homeostasis^7,8^. Supporting this, LD dysregulation is associated with mitochondrial dysfunction, endoplasmic reticulum (ER) stress, and defective lipid signaling^24^. While cultured neurons express LD-regulatory genes and esterify FA into TG^12,25^, it remains unclear whether LD and LD regulation are significant for neuron function *in vivo*.

Here, we combine invertebrate and vertebrate models to establish the presence and functional significance of neuronal LD *in vivo*. Across species, we provide robust evidence that LD are normally present in neurons *in vivo.* We identify multiple regulators of neuronal LD, and show this regulation is significant for maintaining whole-body energy homeostasis. For one gene, *adipose triglyceride lipase* (ATGL), we reveal a conserved and male-specific role in neurons that mediate metabolic and behavioral responses to food withdrawal (‘hunger-activated neurons’) in flies and mice. Mechanistically, ATGL loss in hunger-activated neurons caused profound lipid remodeling leading to defects consistent with mitochondrial and ER dysfunction, which impaired the ability of these neurons to sustain whole-body energy homeostasis. LD regulation therefore coordinates neuronal lipid distribution and utilization to support the function of neurons that maintain whole-body energy homeostasis.

## RESULTS

### Lipid droplets are present in neurons under normal physiological conditions across species

To determine whether neuronal LD form *in vivo* under normal physiological conditions^26^, we used multiple approaches across invertebrate (fly) and vertebrate (mouse) models. Neuronal LD were visualized in adult *Drosophila* males and females with pan-neuronal expression of a LD-targeted GFP (genotype *elav>GFP-LD*)^27,28^. Neuronal LD showed a distinct spatial distribution in the *Drosophila* brain that was consistent between individuals (Fig. 1a-e), an observation reproduced using an independent *UAS-GFP-LD* insertion line (Fig. S1a,b). Most LD were located in the Kenyon cell soma region, neurons whose processes comprise the mushroom body, and in the optic lobes. These regions are implicated in learning and homeostatic regulation of sleep and body fat^29–31^, and in vision^32^, respectively. We next developed and validated an automated method to count neuronal LD in the Kenyon cell soma region as they are found within a compact area (Fig. 1c,e; S1c,d). In 5-day-old adult males and females, we counted ∼400-600 LD per hemisphere in this region (Fig. S1c). Considering there are ∼2200 mushroom body Kenyon cells^33–36^, LD were not present in each neuron. In LD-positive neurons, LD were found mostly in cell bodies and occasionally within axonal projections (Fig. S1e). Neuronal LD are therefore enriched in distinct *Drosophila* brain regions under basal conditions.

**Figure 1.**
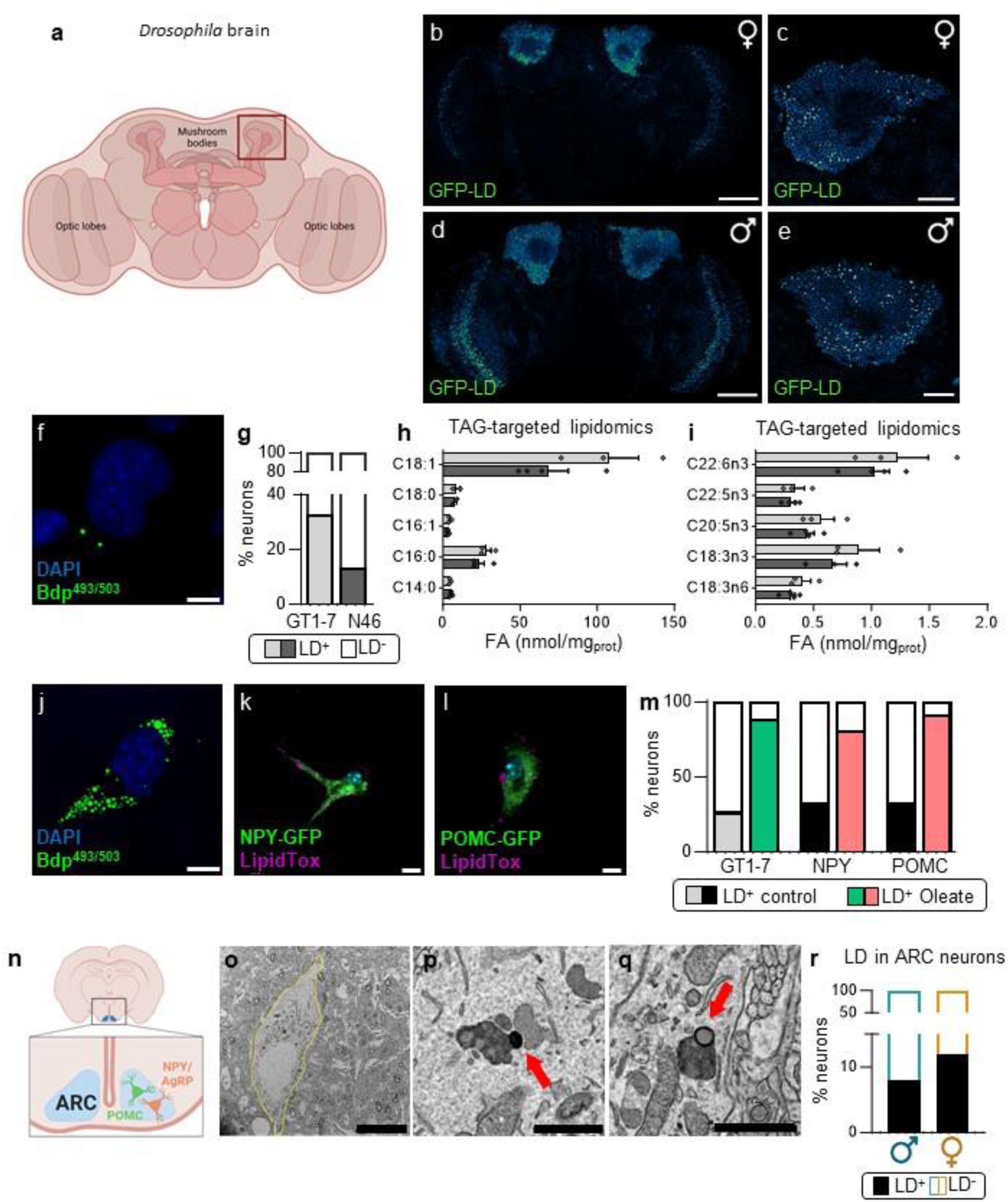
Lipid droplets are present in neurons across species under normal physiological conditions. **a**, Illustration of *Drosophila* brain; mushroom bodies and optic lobes indicated. **b,d**, Z-projection of confocal images of *Drosophila* whole brain and **c,e**, Kenyon cell soma region in 5-day-old female (**b,c**) and male (**d,e**) in *elav>GFP-LD(2.6)* animals. Green punctae represent neuronal lipid droplets (LD). (**b,d**) Scale=100 µm; (**c,e**) scale=20 µm. **f**, Bodipy^493/503^-stained LD in GT1-7 hypothalamic neurons. Scale=10 µm. **g**, Percentage of neurons with one or more LD in GT1-7 and N46 hypothalamic neurons in basal conditions. N=9033 GT1-7 and 28265 N46 cells from 6 independent experiments. **h,i**, Profile of fatty acids (FA) esterified in triglyceride (TG) in GT1-7 (N=3) and N46 neurons (N=4). C14:0= Myristic acid; C16:0= Palmitic acid; C16:1= Palmitoleic acid; C18:0= Stearic acid; C18:1= Oleic acid. Data are represented as mean ± SEM. **j**, LD stained with Bodipy^493/503^ in GT1-7 or with LipidTox in NPY (**k**) and POMC (**l**) primary neurons, supplemented with oleate for 5h. Scales=10, 25, 25 µm. **m**, Percentage of neurons with one or more LD in GT1-7, NPY and POMC neurons treated with vehicle (BSA) or oleate. GT1-7, N=7735 BSA and 6193 oleate; NPY, N=58 BSA and 59 oleate; POMC, N=77 BSA and 105 oleate. **n**, Illustration of the arcuate nucleus (ARC) of the hypothalamus containing hunger-activated Neuropeptide Y (NPY)/Agouti-related peptide (AgRP) and hunger-inhibited Pro- opiomelanorcortin (POMC) neurons which regulate energy homeostasis. **o-q**, Transmission electron microscopy (TEM) of LD (red arrows) in mouse ARC neurons (yellow outline). Scale=10 µm (**o**), 1 µm (**p-q**). **r**, Percentage of neurons containing at least one LD in males (turquoise) (N=53 cells from 2 mice) and females (orange) (N=41 cells from 2 mice). See related data in Supplemental Figure 1.

We next assessed neuronal LD in mammals using mouse hypothalamic neurons, as this brain region shares functional similarities with the *Drosophila* mushroom body. In hypothalamic N46 and GT1-7 neuronal cell lines, lines which co-express Neuropeptide Y (NPY) and Agouti-related peptide (AgRP)^25^, LD were detected with BODIPY 493/503 (Fig. 1f) and quantified using an automated counting system (Fig. 1g, S1f-g). Thirty-three percent of GT1-7 neurons had one or more LD whereas only thirteen percent of N46 neurons were LD-positive (Fig. 1g). Mass spectrometry-based FA profiling revealed oleate (C18:1) and palmitate (C16:0) were the most abundant FA esterified in neuronal LD (Fig. 1h), though additional species were detected including palmitoleate (C16:1), stearate (C18:0), and polyunsaturated FA (Fig. 1i). Supporting FA esterification in neurons, BODIPY-C12 was rapidly esterified in LD of GT1-7 neurons (Fig. S1j-k), and oleate treatment dramatically increased the number of LD-positive GT1-7 neurons (Fig. 1j,m; S1h-i). LipidTox-positive LD were also detected in ∼33% of hunger-activated NPY and hunger- inhibited Pro-opiomelanocortin (POMC) neurons in primary hypothalamic cultures from NPY-GFP and POMC-GFP male and female transgenic mice (Fig. 1k,l), a percentage strongly increased by oleate treatment (Fig. 1m). Together with GT1-7 and N46 data, this suggests mammalian hypothalamic neurons esterify FA in LD. *In vivo* under normal physiological conditions, transmission electron microscopy (TEM) on sections of the arcuate nucleus (ARC), which contains NPY/AgRP and POMC neurons (Fig. 1n), revealed LD in 8% and 12% of neurons from adult male and female mice, respectively (Fig. 1o-r). Thus, our *in vitro* and *in vivo* data across fly and rodent models provide strong evidence that LD form and are present in neurons under normal physiological conditions.

### A network of genes regulates neuronal lipid droplets

We reasoned that if LD normally support neuronal lipid distribution and utilization, genes encoding enzymes that regulate LD esterification, lipolysis, and dynamics (‘LD-regulatory genes’) should be expressed in neurons and participate in LD regulation (Fig. 2a). Our analysis of annotated single-cell RNAseq data confirmed expression of LD-regulatory genes in *Drosophila* neurons (Fig. S2a)^37^ and neurons of the ARC in both male and female mice (Fig. S2b-d)^38^. To test the significance of this expression, we used multiple approaches to inhibit LD-regulatory genes and examine neuronal LD abundance. Based on data from our group and others identifying phenotypes associated with retinal^39,40^ and neuronal inhibition or loss of adipose triglyceride lipase (ATGL)^10,27,41^, and brain lipidomic changes due to whole-body ATGL loss^42^, we cultured hypothalamic neurons with ATGL inhibitor ATGListatin^43^. In basal conditions, ATGListatin treatment led to LD accumulation in GT1-7 (Fig. 2b; Fig. S3a-b) and N46 (Fig. S3c) neurons, showing ATGL regulates basal lipolytic activity in hypothalamic cells.

**Figure 2.**
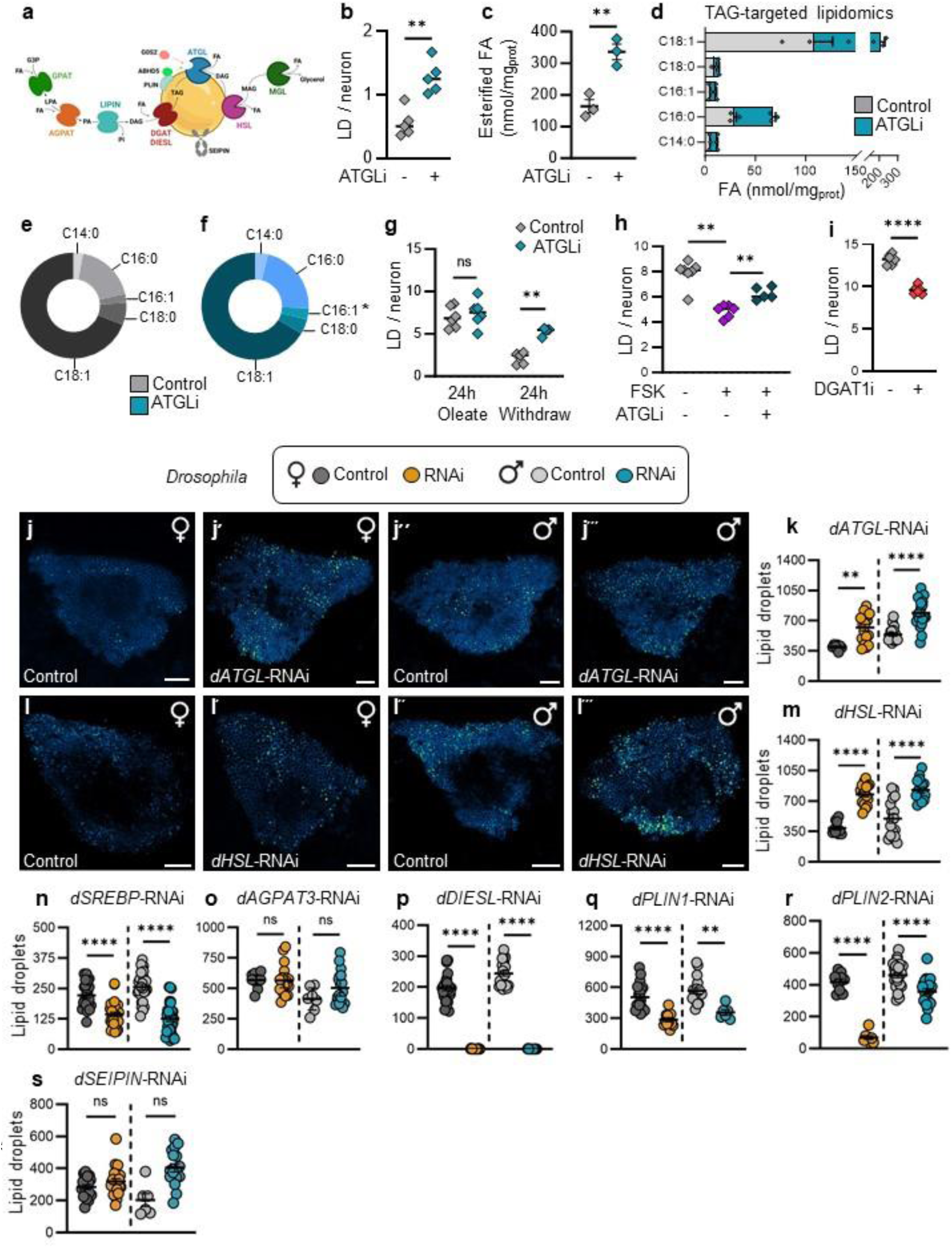
A network of genes regulates neuronal lipid droplets. **a**, LD formation relies on enzymes including GPAT and AGPAT; LIPIN and DGAT/DIESL while LD hydrolysis is mediated by lipases including ATGL, HSL and MGL (Suppl Table 1). The recruitment and activity of lipases is regulated by ABHD5, G0S2, PLIN at the surface of LD and SEIPIN, a docking protein. **b**, Number of LD in GT1-7 neurons treated with vehicle (DMSO) or ATGListatin (24h), N=5. **c**, Total amount of FA esterified into TG in GT1-7 neurons treated with DMSO or ATGListatin (24h). **d**, Profile of FA esterified into TG in GT1-7 cells treated with control (DMSO, control data regraphed from Fig. 1g) or ATGListatin. C14:0, Myristic acid; C16:0, Palmitic acid; C16:1, Palmitoleic acid; C18:0, Stearic acid; C18:1, Oleic acid. N=3 independent experiments. **e,f**, Relative proportion of FA esterified into TG in GT1-7 neurons treated with DMSO or ATGListatin (24h). **g**, Number of LD in GT1-7 neurons incubated with oleate for 24h, or 24h after oleate withdrawal with or without ATGListatin, N=4-6. **h**, Number of LD in GT1-7 neurons (preloaded with oleate) treated with Forskolin (FSK) or FSK + ATGListatin (2.5h), N=5-7. **i**, Number of LD in GT1-7 neurons incubated with oleate and vehicle or DGAT1 inhibitor A-922500 (24h), N=6. **j,l**, Maximum Z-projections and **k,m**, quantification of neuronal LD (green punctae) in the *Drosophila* Kenyon cell soma region of *elav>GFP-LD(3.4)* in adult female (orange) and male (turquoise) flies with neuronal loss of *dATGL* (**j,k**) and d*HSL* (**l,m**). **k,l**, Scale=20 µm. **n-s**, Quantification of neuronal LD in the Kenyon cell soma region of adult *elav>GFP-LD(3.4)* females and males with neuron-specific loss of *dSREBP, dAGPAT3, dDIESL dPLIN1, dPLIN2, dSEIPIN.* (**b,c,i**) Student’s t-test, (**h**) Kruskall-Wallis, (**e,f**) multiple t-test, (**g**) two-way ANOVA with Sidak post-hoc test, (**k,m-o,q-s**) Two-way ANOVA with Tukey post-hoc test. (**p**) Mann-Whitney Test. ns indicates not significant; \**p<0.05,* \*\**p*<0.01; \*\*\*\**p*<0.0001. Data are represented as mean ± SEM. See related data in Supplemental Figures 2 and 3.

Targeted lipidomics revealed total FA esterified in LD was increased by ATGListatin in GT1-7 (Fig. 2c) and N46 neurons (Fig. S3d). We observed greater effects on saturated and monounsaturated FA versus polyunsaturated FA (Fig. 2d, S3e-g), and a greater proportion of esterified palmitoleic acid in GT1-7 and N46 neurons (Fig. 2e-f; Fig. S3h-i), suggesting neuronal ATGL has a substrate preference for palmitoleic acid (C16:1). In oleate-preloaded GT1-7 neurons, ATGListatin blocked the reduction in LD abundance following oleate withdrawal (Fig. 2g), indicating neuronal ATGL eliminates excess neuronal LD. Indeed, pharmacological activation of lipolysis by forskolin in GT1-7 neurons reduced LD number in an ATGL-dependent manner (Fig. 2h). We next assessed how an LD-regulatory gene involved in LD esterification called diacylglycerol O-acyltransferase 1 (DGAT1) affected neuronal LD abundance. DGAT1 inhibition significantly decreased oleate-induced LD formation in GT1-7 neurons (Fig. 2i). Together, these findings reveal FA are esterified in neuronal LD, and identify roles for DGAT1 and ATGL as key regulators of LD in hypothalamic neurons.

To expand knowledge of neuronal LD regulation *in vivo*, we used RNAi to knock down LD- regulatory genes in *Drosophila*. Given results in GT1-7 and N46 neurons, we counted neuronal LD in flies with neuron-specific loss of *dATGL*. Pan-neuronal *dATGL* loss significantly increased LD abundance in neurons of *elav>GFP-LD* males and females (Fig. 2j-k), suggesting *dATGL* normally restricts neuronal LD *in vivo*. Pan-neuronal loss of the *Drosophila* homolog of *hormone-sensitive lipase* (*dHSL*) similarly increased the number of neuronal LD (Fig. 2l-m). Thus, *dHSL* and *dATGL* regulate LD lipolysis in neurons. We next tested if genes that promote FA and TG synthesis in non-neuronal cells affect neuronal LD abundance. Neuron-specific loss of the *Drosophila* homolog of *sterol response element binding protein* (*dSREBP*) reduced neuronal LD in both males and females (Fig. 2n), but neuron-specific loss of the *Drosophila* homolog of *1- acylglycerol-3-phosphate-O-acyltransferase 3* (*dAGPAT3*) had no effect on neuronal LD abundance (Fig. 2o). Pan-neuronal loss of the *Drosophila* homologs of *Lipin* (*dLIPIN*) and *DGAT1* (*dDGAT1*) caused near-total lethality at the late pupal stage (Fig. S3j-k), confirming neurons require these proteins for animal survival^44,45^.

Neuronal loss of the *Drosophila* homolog of newly-identified enzyme *DGAT1/2- independent enzyme synthesizing storage lipids* (*dDIESL*)^46^ that promotes cellular TG levels caused a near-complete loss of neuronal LD in males and females (Fig. 2p) with no effect on survival. Beyond enzymes that directly catalyze steps in TG synthesis and breakdown, neuron-specific loss of the *Drosophila* homologs of LD-regulatory genes *Perilipin 1* (*dPLIN1*) and *Perilipin 2* (*dPLIN2*) reduced neuronal LD abundance (Fig. 2q,r), whereas neuron-specific loss of the *Drosophila* homolog of *Seipin* (*dSEIPIN*) had no effect on LD abundance (Fig. 2s). For most genes, the magnitude of change in LD abundance was equivalent between males and females (Fig 2k, m-s); however, neuronal loss of *dSREBP* affected LD abundance more in males (sex:genotype interaction *p*=0.0179) and neuronal d*PLIN2* loss affected LD more in females (sex:genotype interaction *p*<0.0001). Together, our data reveal a network of genes that regulate neuronal LD and reveal ATGL as a conserved regulator of neuronal LD across invertebrates and vertebrates.

### Genes that regulate neuronal lipid droplets influence whole-body energy homeostasis in flies and worms

We next wanted to determine whether neuronal LD regulation was physiologically significant. FA metabolism in mammalian hypothalamic neurons affects energy homeostasis^47–51^, and we previously showed neuronal loss of *dATGL* impaired *Drosophila* fat breakdown post-fasting, a phenotype associated with whole-body energy homeostasis^27^. We therefore hypothesized that neuronal LD regulation impacts energy homeostasis-related phenotypes (*e.g*., fat storage and/or breakdown in flies, energy intake and/or expenditure in mammals). To test this, we analyzed additional *Drosophila* LD-regulatory genes. We examined fat storage under basal conditions and monitored fat breakdown post-fasting at early (0-12h) and late (12-24h) time points, as distinct mechanisms regulate each phenotype^52^. Neuron-specific loss of *dHSL, dPLIN1, dPLIN2,* and *dDIESL* led to defects in whole-body energy homeostasis (Fig. 3a-l). Pan-neuronal loss of these genes primarily affected fat breakdown; however, there were subtle phenotypic differences between animals with neuron-specific loss of individual LD-regulatory genes, as follows. Neuronal loss of *dHSL* reduced body fat and lowered late fat breakdown in males with no effect in females (Fig. 3a-c), whereas pan-neuronal loss of *dPLIN1* and *dPLIN2* had no effect on body fat in either sex but lowered early fat breakdown only in males (Fig. 3d-i). We reproduced these trends with independent RNAi lines (Fig. S3l,m), and further show pan-neuronal loss of *dDIESL* caused a male-specific increase in body fat and faster early fat breakdown (Fig. 3j-l). Thus, neuronal LD regulation plays a role in maintaining *Drosophila* energy homeostasis, revealing the biological significance of this regulation. Importantly, a role for neuronal ATGL in regulating energy homeostasis is conserved across invertebrates, as we show pan-neuronal RNAi-mediated knockdown of *ATGL* in *C. elegans* similarly increased fat storage (Fig. 3m-p) and impaired fasting-induced fat breakdown (Fig. 3q-u).

**Figure 3.**
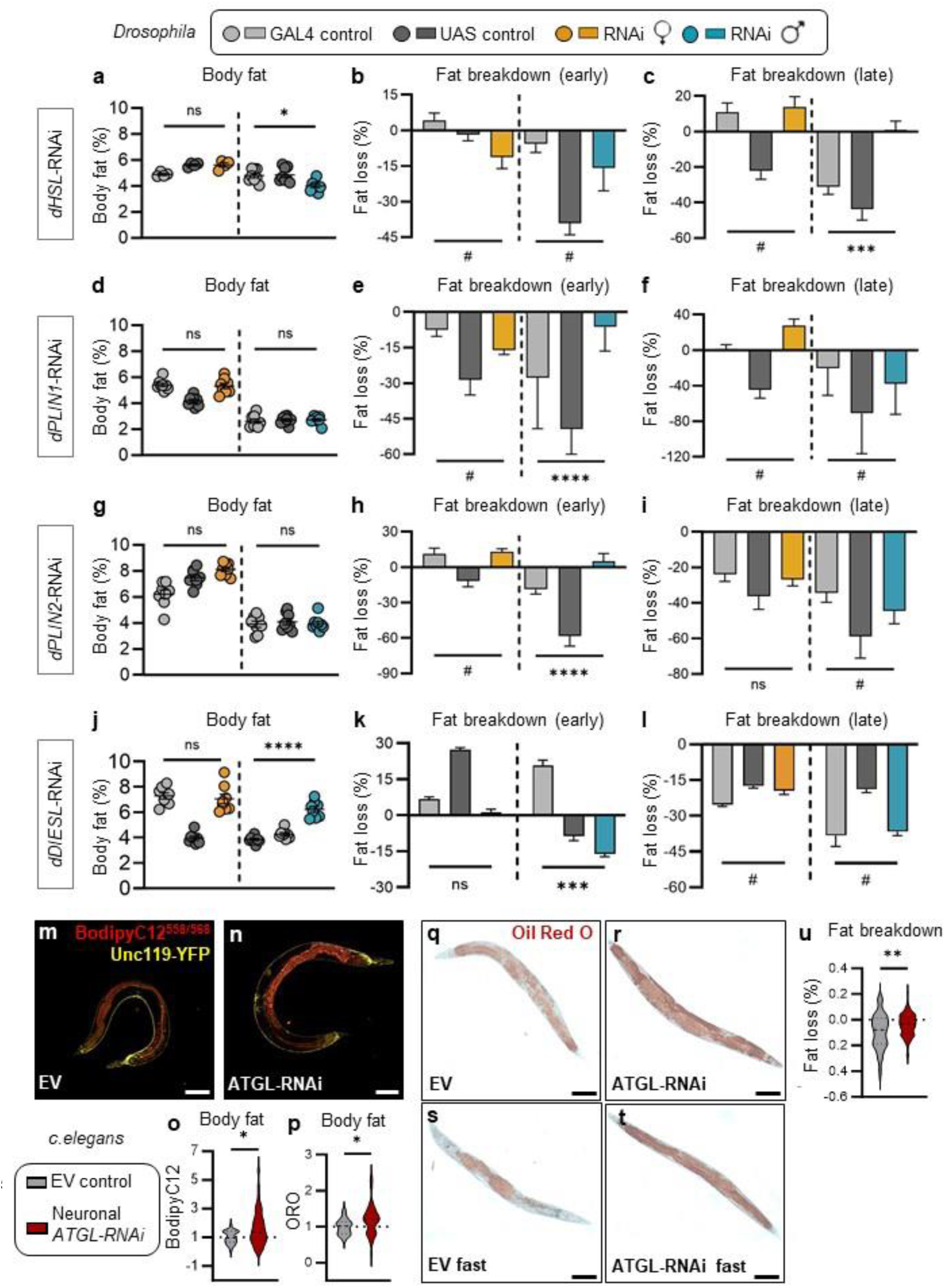
Neuronal lipid droplet regulation affects whole-body energy homeostasis in worms and flies. **a-l**, Whole-body energy homeostasis in adult *Drosophila* males (turquoise) and females (orange) with pan-neuronal loss of genes that encode LD-associated proteins *dHSL* (**a-c**), *dPLIN1* (**d-f**), *dPLIN2* (**g-i**) and *dDIESL* (**j-l**). (**a,d,g,j**) Percent body fat in fed conditions. Mean percent body fat +/- SEM. (**b,e,h,k**) Magnitude of body fat loss from 0-12h post-fasting (early). (**c,f,i,l**) Magnitude of body fat loss from 12-24h post-fasting (late). Fat breakdown data expressed as the mean body fat loss over a given period post-fasting +/- coefficient of error. Two-way ANOVA: ns indicates not significant, \**p<*0.05, *** *p<*0.001, \*\*\*\**p<*0.0001 from RNAi genotype interaction, # indicates control genotype interaction. **m,n**, Fluorescence microscopy of *unc-119p::YFP* neuronal expression (yellow) and Bodipy^558/568^C12-stained LD (red) in neuronal RNAi sensitive worms fed with either empty vector (EV) or *atgl-1* RNAi (*ATGL-* RNAi). Scale=100 µm. **o**, Quantification of Bodipy^558/568^C12 fluorescence in the anterior gut of EV (n=28) vs. ATGL-RNAi (n=42) worms, N=3. **p**, Quantification of Oil Red O staining (ORO) in EV (n=26) vs. ATGL-RNAi (n=43) worms, N=3. **q-u**, ORO staining in EV and ATGL-RNAi fed or fasted worms n=92-103, N=6. Scale=100 µm. (**o,p**) Mann-Whitney, (**u**) Student’s t-test, data are represented as mean ± SEM. See related data in Supplemental Figure 3.

### ATGL functions in arcuate neurons to regulate energy homeostasis in mammals

We next tested whether regulation of neuronal LD was physiologically significant in mammals. In light of the energy homeostasis-related phenotypes we observed across invertebrates due to loss of neuronal ATGL, we targeted ATGL in hypothalamic neurons of the ARC, a key region containing neurons that control energy homeostasis^53^ (Fig. 1n). Cre-mediated knock-out (KO) of ATGL in all ARC neurons in adult mice (ARC^ATGL^KO) (Fig. S4a-b) had no effect on body weight (BW), fat mass, or food intake in chow-fed males or females, but did increase lean mass in males (Fig. 4a-f). Parameters of energy balance including energy expenditure (EE), respiratory quotient (RQ), FA oxidation (FAOx) and glucoregulatory responses were also similar between ARC^ATGL^Ctl and ARC^ATGL^KO males and females (Fig. S4e-p). Therefore, loss of neuronal ATGL in ARC neurons did not affect whole-body energy metabolism in unchallenged conditions, in line with a lack of phenotype in flies with pan-neuronal *dATGL* loss under basal conditions^27^. Because fat breakdown post-fasting was impaired in flies lacking neuronal *dATGL*, indicating abnormal responses to metabolic challenge, we measured ATGL expression in the ARC after fasting and a cold exposure. As we show in liver, ATGL expression was upregulated by fasting and cold in the ARC of male mice (Fig. S4c-d).

**Figure 4.**
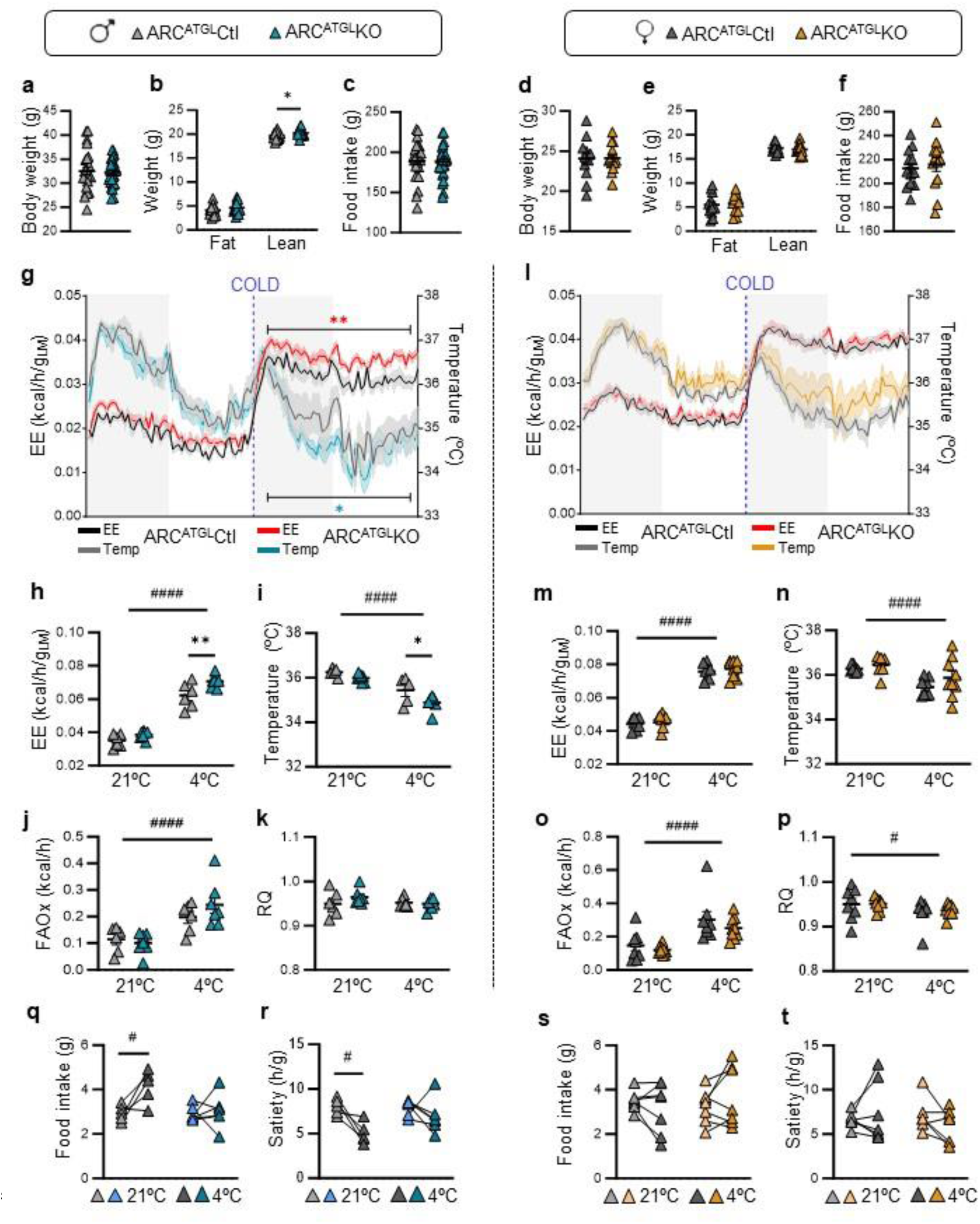
Neuronal ATGL influences whole-body energy homeostasis in mammals. **a,d**, Body weight, **b,e**, fat and lean mass, and **c,f**, cumulative food intake in 16-week-old male and female ARC^ATGL^CRE and ARC^ATGL^KO mice, N=24-28 males and 13 females. **g-n**, Energy expenditure (EE) and body temperature traces with corresponding quantifications. **j,o**, Fatty acid oxidation (FAOx), **k,p**, respiratory quotient (RQ), **q-t**, food intake, and satiety in ARC^ATGL^CRE and ARC^ATGL^KO males and females measured in metabolic cages during 24h at 21 °C or 24h at 4 °C. N=6-7 males and 8-9 females. Data are represented as mean ± SEM. (**a-f**) Student’s t-test, *p<0.05; (**g-t**) Two-way ANOVA: #*p<*0.05, ####*p<*0.0001, time interaction and \**p<*0.05, \*\**p<*0.01, genotype interaction, Sidak post-hoc. See related data in Supplemental Figure 4.

We therefore subjected distinct cohorts of ARC^ATGL^KO mice and controls to either a 16 h fast or cold exposure (4°C) for 24 h. For both ARC^ATGL^KO and ARC^ATGL^Ctl males and females, fasting reduced EE and RQ and increased FAOx (p<0.0001, two-way ANOVA) (Fig. S4e-h,k-n). Because these responses agree with known effects of fasting, loss of ATGL in ARC neurons did not affect fasting-induced metabolic responses. In contrast, while cold exposure caused the expected responses in EE, FAOx, and body temperature in ARC^ATGL^Ctl males and females (Fig. 4g-p) (p<0.0001, two-way ANOVA), male ARC^ATGL^KO mice showed a more pronounced increase in EE with a greater and faster drop in body temperature compared with ARC^ATGL^Ctl mice (Fig. 4g-i, Fig. S4q). These data suggest ATGL loss in ARC neurons impairs energy homeostasis during cold. Indeed, male ARC^ATGL^KO mice showed impaired cold-induced food intake and satiety response compared with ARC^ATGL^Ctl mice (Fig. 4q-r). Plasma FA levels after cold were similar between male ARC^ATGL^KO and ARC^ATGL^Ctl mice (Fig. S4r), suggesting loss of ATGL did not affect peripheral lipolysis. Importantly, ATGL loss in ARC neurons did not impair female physiological responses to cold (Fig.4l-p,s-t; S4s), consistent with the male-specific effect of neuronal *dATGL* loss on *Drosophila* fat breakdown^27^. Our data from multiple models therefore suggest neuronal ATGL plays a conserved and male-specific role in regulating energy homeostasis during metabolic challenges, confirming ATGL-dependent LD regulation in neurons is physiologically significant.

### ATGL functions within AgRP neurons to influence whole-body energy homeostasis

A key step toward identifying the mechanisms by which ATGL acts in neurons to influence energy homeostasis was to identify a homogeneous population of neurons in the ARC that requires ATGL function. Reduced feeding in the ARC^ATGL^KO mice reproduced phenotypes associated with decreased activity of hunger-activated AgRP neurons^54^. We therefore generated a mouse model with a KO of ATGL in AgRP neurons in the NPY-GFP genetic background to easily visualize AgRP neurons (known to co-express NPY); loss of ATGL was validated by RNAscope and qPCR (Fig. S5a-d). A stereological analysis showed that ATGL loss in AgRP neurons did not affect the number of NPY-GFP-positive cells in the ARC (Fig. S5e-g) nor the density of NPY-GFP fibers projecting in the paraventricular nucleus (PVN) in either sex (Fig. S5h-j), ruling out a developmental effect of ATGL loss.

Under *ad libitum* feeding conditions, male AgRP^ATGL^KO mice had reduced BW, perigonadal fat, lean mass, and cumulative food intake (Fig. 5a-c,S5k). These changes cannot be explained by altered growth, as femur length was similar between AgRP^ATGL^CRE and AgRP^ATGL^KO (Fig. S5l). Glucoregulatory responses were also normal in male AgRP^ATGL^KO mice (Fig. S5m-o). However, we observed metabolic changes consistent with the lean phenotype in AgRP^ATGL^KO mice, including increased EE and RQ in fed conditions (Fig. 5g-h,k). During fasting, only increased EE was maintained in AgRP^ATGL^KO males, suggesting the RQ phenotype depends on food availability. Female AgRP^ATGL^KO mice showed a non-significant trend towards lower BW and lean mass in fed conditions (*p* = 0.055) with no other changes in feeding or metabolic parameters in either the fed or fasted state (Fig. 5d-f, i-j,m-n, S5p). AgRP-specific loss of ATGL therefore disrupts energy homeostasis in males during both normal and fasted contexts.

**Figure 5.**
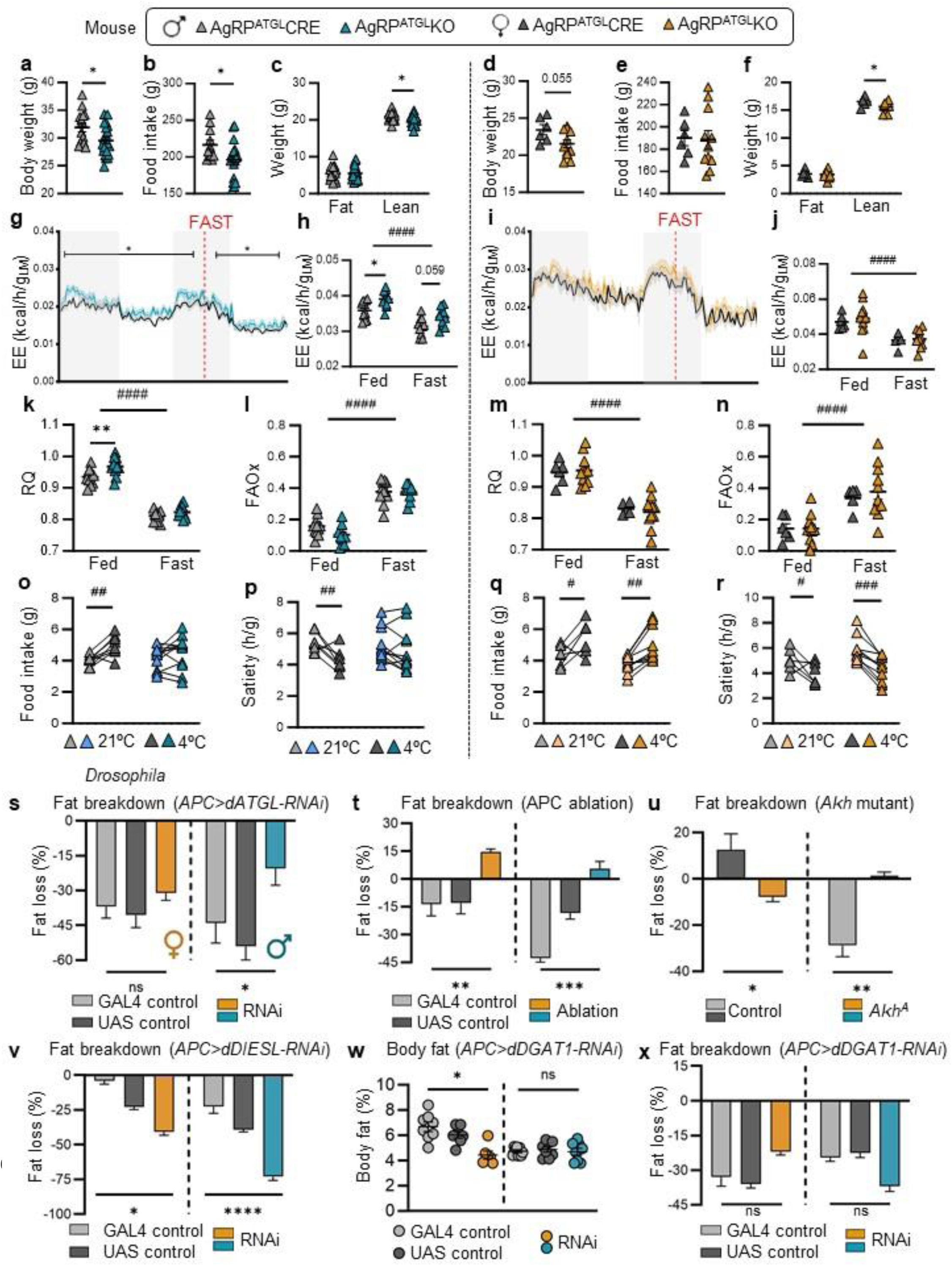
ATGL function within AgRP neurons plays a conserved role in regulating whole-body energy homeostasis. **a,b**, Body weight, **b,e**, cumulative food intake, **c,f**, fat and lean mass in 16- week-old male and female AgRP^ATGL^CRE and AgRP^ATGL^KO mice, N=14-21 males and 6-11 females. **g-j**, EE, **k,m**, RQ and **l,n**, FAOx in AgRP^ATGL^CRE and AgRP^ATGL^KO males and females in *ad libitum* (Fed) or fasted (16h) conditions (Fast). N=10-11 males and 6-11 females. **o-r**, Food intake and satiety in AgRP^ATGL^CRE and AgRP^ATGL^KO males and females over 24h at 21 °C or 24h at 4 °C. N=8-11 males and 6-10 females. Data are represented as mean ± SEM. (**a-f**) Student’s t-test, \**p<*0.05. (**g-r**) Two-way ANOVA: #*p<*0.05, ##*p<*0.01, ####*p<*0.0001, time interaction and \**p<*0.05, \*\**p<*0.01, genotype interaction, Sidak post-hoc. **s**, Whole-body fat breakdown 0-24h post-fasting in female (orange) and male (turquoise) flies with loss of *dATGL* in the adipokinetic hormone (Akh)-producing cells (APC). **t**, Fat breakdown 12-24h post-fasting in flies in which the APC were ablated via overexpression of proapoptotic gene *reaper* (*rpr*). **u**, Fat breakdown 0-12h post-fasting in *Akh* mutant flies (Akh^A^). **v**, Fat breakdown 12-24h post-fasting in flies with APC-specific loss of *dDIESL*. **w**, Body fat in flies with APC-specific loss of *dDGAT1.* **x**, Fat breakdown 12-24h post- fasting in flies with APC-specific loss of *dDGAT1*. Body fat shown as mean +/- SEM. Fat breakdown data expressed as the mean percent body fat loss post-fasting +/- coefficient of error. Two-way ANOVA: ns indicates not significant, \**p<*0.05, \*\**p<*0.01, \*\*\**p<*0.001, \*\*\*\**p<*0.0001, RNAi genotype interaction. See related data in Supplemental Figures 5 and 6.

During cold exposure, male AgRP^ATGL^CRE and AgRP^ATGL^KO mice showed normal metabolic responses to cold (Fig. S5q-t) (p<0.0001, two-way ANOVA); however, AgRP^ATGL^KO males had impaired cold-induced food intake and satiety response compared with AgRP^ATGL^CRE littermates (Fig. 5o-p). Loss of ATGL in AgRP neurons of female mice did not affect cold responses (Fig. 5q-r; Fig. S5u-x). Importantly, the hypophagia and lean phenotypes in AgRP^ATGL^KO males are consistent with reduced AgRP tone^54–56^. AgRP neurons therefore represent one group of neurons in which ATGL function is required to regulate energy homeostasis in basal and challenged conditions. Considering loss of HSL in AgRP neurons had no effect on energy balance^48^, our data suggest a specific role for ATGL as a lipase that supports AgRP function.

### Lipid droplet regulation in *Drosophila* hunger-activated neurons affects whole-body energy homeostasis

Like murine AgRP neurons, the ∼20 *Drosophila* adipokinetic hormone (Akh)-producing cells (APC) are neuroendocrine cells activated during nutrient deprivation to regulate food-seeking behaviours^57,58^. APC also mediate peripheral lipolysis by releasing Akh, a neuropeptide that binds the Akh receptor to stimulate lipolysis^52,59^. We therefore asked whether *dATGL* acts in hunger- activated APC to regulate *Drosophila* whole-body energy homeostasis, as we described for hunger-activated AgRP neurons. We used *Akh-GAL4*, an APC-specific GAL4 driver, to knock down *dATGL* specifically in fly APC. APC-specific *dATGL* loss did not affect body fat in either adult male or female flies (Fig. S6a). Because APC ablation, inhibition, and loss of Akh augment body fat in both sexes^60^, APC-specific *dATGL* loss does not cause APC death or a complete loss of function. Post-fasting, we observed reduced fat breakdown in males but not females with APC-specific loss of *dATGL* over 24 h (Fig. 5s). This reproduced the decreased fat breakdown phenotype caused by reduced APC function or loss of Akh peptide (Fig. 5t,u; Fig. S6b-e), and suggests ATGL-dependent LD lipolysis normally supports APC function. Importantly, impaired fat breakdown was specific to APC-dependent production of Akh, as loss of another APC-derived peptide had no effect on this phenotype in either sex (Fig. S6f,g).

To further assess the relationship between LD regulation and APC function, we used RNAi to knock down additional LD-regulatory genes in the APC. APC-specific loss of *dHSL* had no effect on fat storage (Fig. S6h) or fat breakdown (Fig. S6i,j). This mirrors mouse data showing a specific role for ATGL (Fig. 5) and not HSL^48^ in AgRP neurons, and suggests other neurons mediate the effect of pan-neuronal *dHSL* loss on fat breakdown. We next tested how genes that promote LD esterification affect APC function. APC-specific loss of *dDIESL* caused a strong increase in fat breakdown post-fasting in both sexes (Fig. 5v; S6k,l). APC-specific loss of *dDGAT1* both lowered body fat in females (Fig. 5w) and showed a trend toward faster fat breakdown (Fig. 5x; S6m). Because these data reproduce the lean phenotypes^27^ and rapid fat breakdown caused by genetic activation of APC using bacterial sodium channel NaChBac (Fig. S6n,o), this suggests LD formation normally restrains APC activation. In *Drosophila*, LD formation and degradation by LD-regulatory genes therefore ensures the appropriate function of hunger-activated neurons to maintain energy homeostasis. Importantly, the relationship between neuronal LD regulation and energy homeostasis is strongest in male flies. Given that LD-regulatory genes have similar effects on neuronal LD in *Drosophila* males and females (Fig. 2j-s), and APC influence energy homeostasis in both sexes (Fig. 5t,u; S6b-e,n-o)^60^, this reveals a sex difference in the ability of hunger-activated neurons to maintain function when LD regulation is perturbed.

### Neuron-specific loss of ATGL has profound and sex-specific effects on the lipidome

To uncover how LD regulation ensures appropriate function of hunger-activated neurons in flies and mammals, we used multiple approaches and models to evaluate how loss of ATGL affects neuronal lipid metabolism. In flies, we isolated whole brains from male and female adult flies with neuronal loss of *dATGL* and subjected them to mass spectrometry (MS)-based untargeted lipidomic profiling. We detected 764 lipid features and noted sex differences in multiple lipid classes (Fig. S7a; Supplemental table 1). Loss of neuronal *dATGL* caused sex-specific changes in lipid abundance (Fig. S7b,c): males had significant alterations in 133/764 features and females had significant changes in 64/764 features (Supplemental table 1). This included changes to lipid classes such as neutral lipids, FA, and phospholipids (Fig. 6a,b; Fig. S7d-g; Supplemental table 1). In GT1-7 neurons (male origin), we performed untargeted lipidomics by MS following ATGListatin treatment. In positive ion mode, 1261 lipid features were detected. Of these, 116 were significantly affected by ATGL inhibition including 36 annotated lipids. Principal component analysis revealed a major effect of ATGListatin on lipid (sub)classes (Fig. 6d): component 1 explained 36% and component 2 18% of the variance between groups.

**Figure 6.**
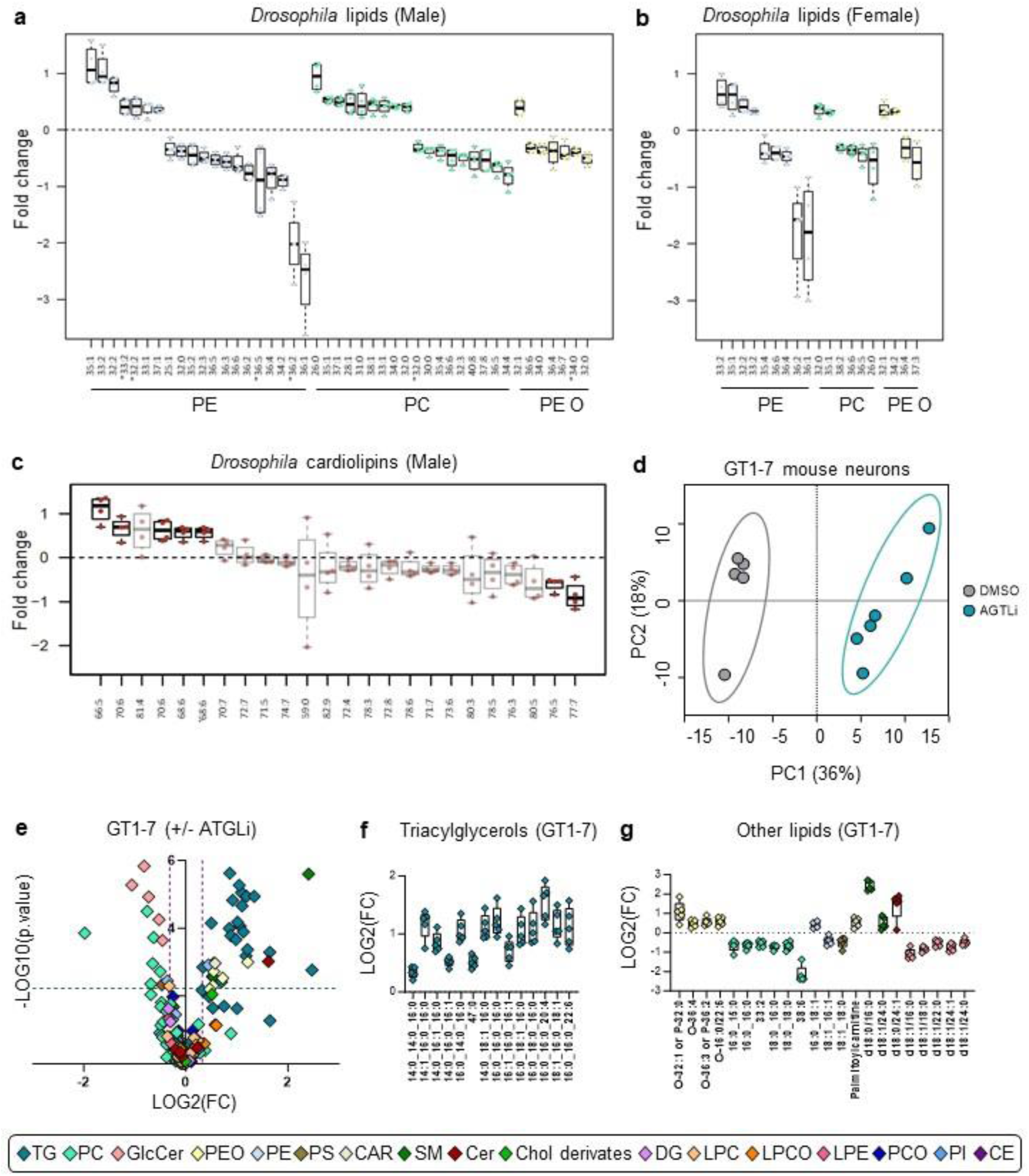
Profound and sex-specific lipid remodeling caused by neuronal loss of ATGL. **a**, Differentially regulated lipid species in brains of *Drosophila* adult males and **b**, females with neuronal loss of *dATGL*. **c**, Differentially regulated cardiolipin species in *Drosophila* adult male brains with neuronal loss of *dATGL*; black boxes indicate significantly altered lipids and grey boxes indicate trends. **d**, Principal component analysis and **e,** volcano plot from LC-QTOF-based lipidomics in GT1-7 cells from the 188 annotated MS features obtained following MS data processing. In the volcano plot, the x axis represents the fold changes of MS signal intensities expressed as log2 for all the features in the ATGListatin group compared with control group. The y axis corresponds to the p values expressed as -log10. The color plots show the annotated lipid entities. N=5-6. **f,g**, Box plots of annotated unique lipids (expressed as log2) significantly discriminating 24h ATGListatin treatment from control group. Multiple t-test; N=5-6. PE= Diacylglycerophosphoethanolamines; PC= Diacylglycerophosphocholines; PEO= 1-alkyl,2- acylglycerophosphoethanolamines; CL= Cardiolipin; Cer= Ceramide; SM= Ceramide phosphocholines (sphingomyelins); GlcCer= Simple Glc series; S= Diacylglycerophosphoserines; CAR= Fatty-acyl-carnitines; PI= Diacylglycerophosphoinositols; PCO= 1-alkyl,2- acylglycerophosphocholines; LPCO= Monoalkylglycerophosphocholines; LPE= Monoacylglycerophosphoethanolamines; DG= Diacylglycerols; PC= Monoacylglycerophosphocholines; CE= Steryl esters; Chol derivates= Cholesterol and derivates; TG= Triacylglycerols. See related data in Supplemental Figures 7 and 8.

ATGL inhibition in GT1-7 neurons affected several lipid (sub)classes, with major increases in TG content (Fig. 6e-g, Fig. S8a-b). Increased TG abundance across multiple species was also observed in *Drosophila* male brains (Fig. S7f); only two TG species were altered in females (Fig. S7g). Overall, ATGL loss caused significant dysregulation of phospholipids in both male fly brains and cultured mouse neurons. For example, loss of *dATGL* significantly reduced levels of two cardiolipin (CL) species in *Drosophila* male brains (Fig. 6c) with a similar trend in 14/24 other CL species (Supplemental table 1); four CL species showed increased abundance. Because CL is enriched on the inner mitochondrial membrane and is essential for optimal function of the electron transport chain^61^, these data suggest neuronal loss of *dATGL* caused mitochondrial defects in male brains. Supporting this, ATGListatin treatment in GT1-7 neurons led to increased palmitoylcarnitine levels, suggestive of dysfunctional FAOx (Fig. 6g).

Loss of neuronal *dATGL* in male flies and ATGListatin treatment of cultured neurons also altered membrane phospholipids including diacylglycerophosphoethanolamines (PE), diacylglycerophosphocholines (PC), diacylglycerophosphoserines (PS), sphingolipids and glucosylceramides, and 1-alkyl, 2-acylglycerophosphocholines (PCO) and 1-alkyl, 2- acylglycerophosphoethanolamines (PEO) (Fig. 6a,e,g). Despite differences in the identity of individual phospholipids dysregulated by loss of ATGL in cultured neurons and *Drosophila* male brains, data from both systems suggest a major remodeling of phospholipid species (see Supplemental table 1, Fig. 6a,g Fig. S7a,d). For example, a notable trend was a disruption in PE and PC levels. In *Drosophila* male brains, the majority of PE species that were significantly dysregulated were lower in abundance (12/19) (Fig. 6a). Considering many PC species were significantly increased, this suggests the PC:PE ratio was increased in *Drosophila* brains. In ATGListatin-treated neurons, we saw higher PE levels and lower PC levels (Fig. 6g), suggesting a lower PC:PE ratio. While these data show opposite trends in PE and PC levels upon loss of ATGL, both high and low PC:PE ratios have been linked with organelle dysfunction and ER stress^62^. Given PC and PE, along with ether lipids and glucosylceramides, contribute to the integrity and function of plasma membranes including lipid rafts and the endomembrane system^63,64^, this suggests ATGL is required to maintain membrane lipid composition and function in neurons.

In line with a lack of whole-body energy homeostasis phenotypes in *Drosophila* females with neuron-specific loss of *dATGL*, there was no decrease in any CL species, fewer PE species with lower abundance, and very few PC species with increased abundance in female flies with neuronal *dATGL* loss (Fig. 6b, Fig. S8c). Because neuron-specific loss of *dATGL* caused an equivalent increase in neuronal LD between males and females (Fig. 2k), this suggests compensatory mechanisms in females maintain neuronal lipid metabolism despite loss of *dATGL*- mediated LD lipolysis. Together, our *Drosophila* and mammalian lipidomic data indicate ATGL controls LD to regulate lipid distribution and utilization in neurons, including lipids that support membrane homeostasis in mitochondria and the ER.

### Neuronal ATGL supports mitochondrial and ER homeostasis to promote neuron function

To determine how dysregulation of lipid distribution and utilization due to altered LD regulation impacts hunger-activated APC and AgRP function, we tested whether loss or inhibition of ATGL in these neurons affected mitochondria and the ER. In flies, APC-specific loss of *dATGL* reduced the number of mitochondria in male but not female APC (Fig. 7a). This aligns with our data showing a male-specific decrease in CL species (Fig. 6c) and male-specific fat breakdown defects (Fig. 5s). We used TEM to examine organelles in male AgRP neurons using GFP immunolabeling (NPY-GFP strain). Despite a similar number of LD (Fig S9a-d), likely due to thin sectioning (70 nm) or low percentage of LD-positive neurons in image frames, and mitochondria in neurons of AgRP^ATGL^CRE and AgRP^ATGL^KO males, mitochondrial morphology (length and aspect ratio) was reduced in AgRP^ATGL^KO males suggesting mitochondrial dysfunction (Fig. 7b-e). Supporting this, our metabolomics dataset shows ATGL inhibition in GT1-7 neurons increased both glycolysis intermediates and the AMP/ATP ratio (Fig. 7f, Fig. S9f-i), whereas amino acids that replenish the TCA cycle (*e.g*., aspartate, glutamine) were reduced, suggesting changes to TCA anaplerotic pathways (Fig. 7f, S9k-l). Together with an increased palmitoylcarnitine level (Fig 6g), this suggests decreased mitochondrial oxidative capacity in AgRP neurons. In agreement, loss of ATGL in AgRP neurons phenocopies whole-body metabolic phenotypes observed in mice with defective FAOx in AgRP neurons^47,49,51^.

**Figure 7.**
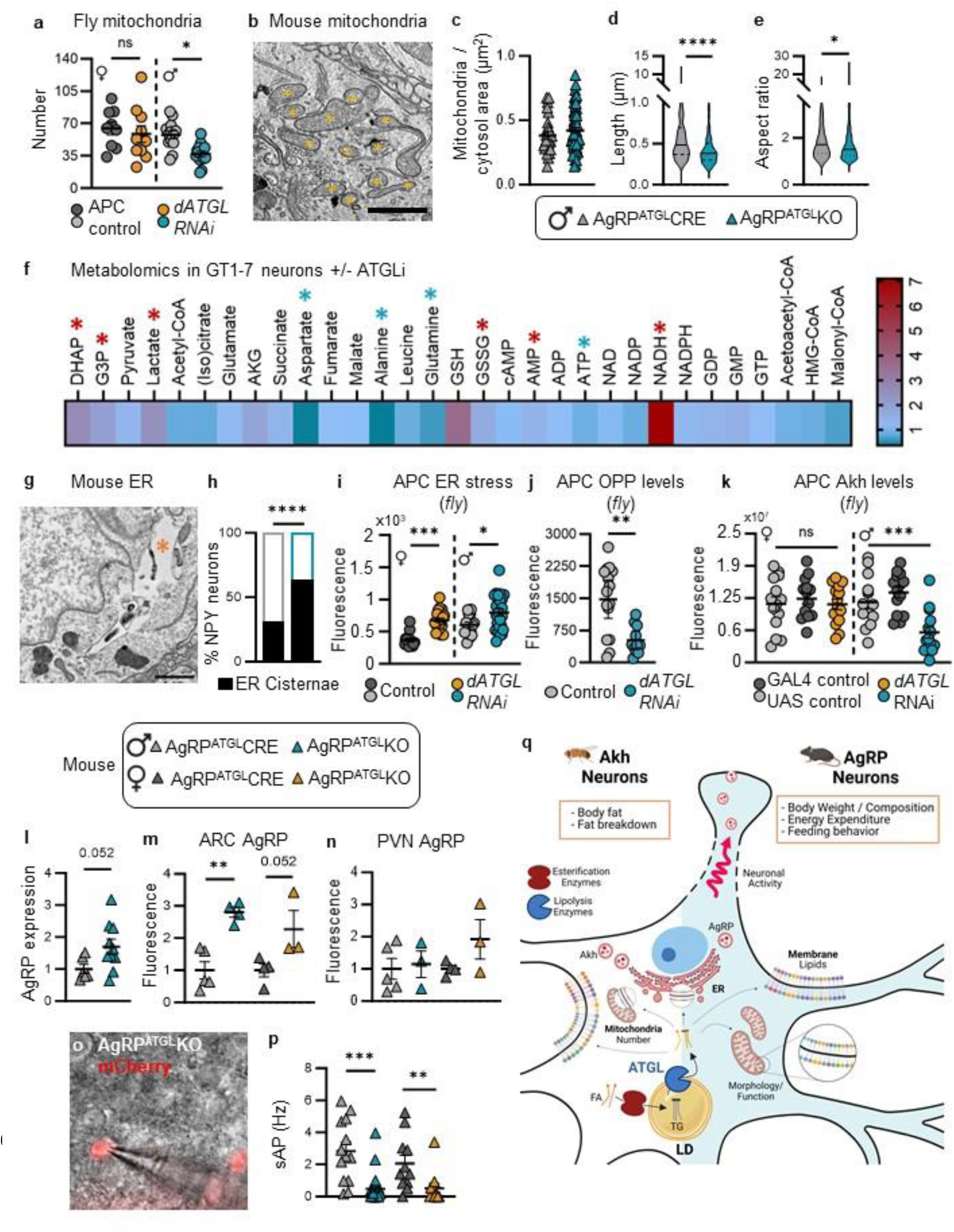
Loss of ATGL is associated with mitochondrial defects and ER stress in hunger-activated neurons. **a**, Mitochondrial number in *Drosophila* female (orange) and male (turquoise) adipokinetic hormone (Akh)-producing cells (APC) with APC-specific loss of *dATGL*. Mean +/- SEM; two-way ANOVA and Tukey post-hoc test. **b**, Electron microscopy of mitochondria in AgRP neurons identified by GFP immunostaining (*yellow). Scale=1 µm. **c**, Mitochondria number, **d**, length, and **e**, aspect ratio (length/width) in male AgRP^ATGL^CRE (n= 2) vs AgRP^ATGL^KO (n=2) mice. N= 26 CRE and 53 KO neurons. (**d-e**) Mean +/- SEM; student t-test. **f**, Relative metabolite levels in response to control and ATGListatin treatment (24h) in GT1-7 neurons. Significant species are annotated by an asterisk (increased red, decreased blue). Mean +/- SEM; multiple t-test; N=7-8. **g**, ER cisternae (*orange) in AgRP neurons. Scale=1 µm. **h**, Percentage of neurons with ER cisternae in AgRP^ATGL^CRE and AgRP^ATGL^KO males. Fisher’s exact test. **i**, Levels of a GFP-based ER stress reporter in *Drosophila* APC. GFP is produced only in contexts where *Xbp1* is spliced in an IRE1-dependent manner ^93^. GFP levels in females (orange) and males (turquoise) with APC- specific loss of *dATGL*. Mean +/- SEM; two-way ANOVA, Tukey post-hoc test. **j**, Nascent protein synthesis in APC of male controls (grey) and males with APC-specific *dATGL* loss (turquoise). Mean +/- SEM; Student’s *t*-test. **k**, Akh protein levels in the APC of adult females (orange) and males (turquoise) with APC-specific loss of *dATGL*. Mean +/- SEM. Two-way ANOVA, Tukey post-hoc test. **l**, AgRP mRNA level in the ARC of AgRP^ATGL^CRE vs AgRP^ATGL^KO males. N=6-10. **m,n**, AgRP immunofluorescence in the ARC (**m**) and the PVN (**n**) from AgRP^ATGL^CRE and AgRP^ATGL^KO males and females (fold change from controls). Mean +/- SEM, (**l**) Student’s t-test, (**m**) one-way ANOVA, Sidak post-hoc test; N=4-5 males and 3-4 females. **o-p**, Spontaneous action potentials (sAP) of AgRP neurons in male and female AgRP^ATGL^CRE vs AgRP^ATGL^KO mice. N=13 vs 25 males; 11 vs 11 females. Kruskal-Wallis test * indicates *p<*0.05, ** indicates p<0.01, *** indicates p<0.001, **** indicates p<0.0001, ns: not significant. **q**, Graphical abstract. See related data in Supplemental Figure 9.

Beyond mitochondria, TEM analysis of AgRP neurons showed a greater percentage of neurons with ER cisternae in male AgRP^ATGL^KO mice (Fig. 7g,h). Because ER cisternae are associated with ER stress^65–67^, this suggests ER stress is elevated in AgRP neurons of AgRP^ATGL^KO mice. Supporting this mouse data, RNAseq on dissected *Drosophila* brains showed upregulation of factors involved in cell and ER stress in both sexes (e.g., *mTerf3*, *Hsp70Bb*) and reduced expression of genes that regulate ER and Golgi function (e.g., *GM130*, *CG14715*) (Supplemental table 2). We also detected higher levels of a *Drosophila* ER stress reporter in APC lacking *dATGL* in both sexes (Fig. 7i) and showed reduced levels of newly-synthesized protein in male APC in flies with APC-specific loss of *dATGL* (Fig. 7j). Together, these data indicate that APC/AgRP neurons show phenotypes consistent with mitochondrial dysfunction and ER stress.

Changes to mitochondria and the endomembrane system including the ER are associated with altered AgRP expression and firing under negative energy balance^47,49,68–70^. To determine how cellular defects associated with ATGL loss impact APC and AgRP neurons, we used multiple approaches to monitor the function of hunger-activated neurons in flies and mice. In flies, APC-dependent production of Akh directly initiates fat breakdown post-fasting^52,59^. We found significantly lower Akh levels in adult male but not female flies compared with sex-matched controls (Fig. 7k). Neuronal *dATGL* therefore influences whole-body energy homeostasis in male flies by supporting APC mitochondrial and ER function to maintain appropriate Akh levels. In females, energy homeostasis was not disturbed by neuronal loss of *dATGL* because Akh production remained intact, possibly because females upregulated a sugar transporter and acyl-CoA synthetase family member (Supplemental table 2) to provide substrates to sustain cellular lipid metabolism without *dATGL*.

In mice, AgRP mRNA and peptide levels were both increased in the ARC of male AgRP^ATGL^KO mice, with no changes in AgRP peptide level in the PVN where AgRP neurons project (Fig. 7l-n, S9m-p). In AgRP^ATGL^KO females, there was a non-significant trend towards increased AgRP peptide levels in the ARC (*p* = 0.06) and PVN (*p* = 0.1) (Fig. 7m-n). Increased AgRP levels in the ARC align with increased ER cisternae in male AgRP^ATGL^ KO mice (Fig. 7g-h) and reports showing ER stress augments AgRP expression^68–70^. Whole-cell patch-clamp recordings (Fig. 7o) of AgRP neurons in AgRP^ATGL^CRE and KO mice revealed that loss of ATGL reduced action potential frequency in male and female AgRP neurons (Fig. 7p, S9q,r). Because loss of ATGL in male AgRP neurons largely reproduces feeding and metabolic phenotypes associated with the silencing of AgRP neurons^54–56^, our data suggest that reduced AgRP neuron firing explains how neuronal loss of ATGL affects energy homeostasis in males. Why a decrease in AgRP neuron firing in AgRP^ATGL^ KO females was not sufficient to disrupt energy homeostasis remains unclear. Altogether, our data in flies and mice uncovers a functionally significant and male-biased role for LD regulation in supporting mitochondria and ER homeostasis to maintain appropriate function in hunger-activated neurons.

## DISCUSSION

Our data establish that LD are normally present in neurons *in vivo*. Based on this discovery, we identified multiple regulators of neuronal LD and show that this regulation plays a physiologically significant role in maintaining whole-body energy homeostasis. For at least one gene, ATGL, we show LD regulation contributes to neuron function by providing lipids that support ER and mitochondrial homeostasis. Neuronal LD regulation is significant in hunger-activated neurons in male flies and mice, where disrupting LD regulation in these neurons impairs their ability to maintain whole-body energy homeostasis. Taken together, our findings reveal LD as a functionally important organelle in neurons, where LD regulation plays a key role in coordinating neuronal lipid supply and utilization.

Our finding that LD are present in neurons challenges the prevailing view that LD do not form *in vivo* under normal physiological conditions. The reason that LD were not previously observed in neurons is likely related to the difficulty in visualizing neuronal LD *in vivo*: neuronal LD are small and present in relatively few neurons (∼10% of ARC neurons), lowering the probability of finding an LD-positive neuron. This low number of LD-positive neurons agrees with a recent survey of PLIN2 expression in the brain^71^, and may be attributed to rapid LD turnover that does not favor LD accumulation. Supporting this, neuronal loss of lipases *dATGL* and *dHSL* promote LD accumulation, and loss of ATGL causes profound remodeling of the lipidome, suggesting neurons have a significant basal lipolytic rate. Considering neuronal LD are infrequent and smaller than LD in other brain cell types such as ependymocytes^71,72^, and the limited sensitivity of LD imaging methods using dyes, it is not surprising that no studies detected neuronal LD *in vivo*. Adding further complexity, neuronal LD are not uniformly distributed across brain regions. Given that changes in activity alter neuronal LD abundance in cultured mouse neurons^14,17^, between-region disparities in neuron activity may explain this differential LD abundance. Indeed, activity is detected in *Drosophila* mushroom body neurons where LD are abundant even when flies are resting^73^.

Beyond identifying neuronal LD *in vivo*, we advanced our understanding of neuronal LD regulation by identifying genes that influence LD abundance. We uncovered a conserved role for ATGL in restricting LD abundance under basal conditions across models, consistent with recently-published roles for ATGL in regulating neuronal LD abundance in *C. elegans*^41^ and LD lipolysis in neuronal cell line or primary neurons in cooperation with lipase DDHD2^10,14,16,74^. We also show DGAT1 inhibition reduced neuronal LD, in line with DGAT1-dependent LD regulation in mammalian retinal cells^44,75^. The congruence between data from our group and others on neuronal DGAT1 and ATGL, and the equivalent effect of ATGL on neuronal LD across worms, flies, and mammals suggests neuronal LD regulation is highly conserved. Additional LD- regulatory genes we identified in flies may therefore play similar roles in mammals, a possibility to test in future studies. Follow-up studies will also need to test a broader network of LD-regulatory genes to determine similarities and differences in regulation of LD between neuronal and non- neuronal cells^7,8,76^. For example, we found that neuronal LD were not affected by loss of *dSEIPIN*, which is essential for LD biogenesis in many non-neuronal cells^77,78^. Given the known role for SEIPIN in converting small nascent droplets into larger mature LD^79^ and the small size of neuronal LD, this may explain why neuron-specific loss of *dSEIPIN* had no effect on LD abundance. Future studies will similarly determine whether lipophagy influences LD during normal conditions, as it does during stress^12,17^, and to determine the source of substrates used for neuronal LD biogenesis^6,80^.

Our mechanistic studies on one LD-regulatory gene, ATGL, provide fundamental insights into how LD regulation supports neuron function. Beyond TG regulation^10,81–83^, our data reveal a role for ATGL in regulating neuronal phospholipids; specifically phospholipids that support mitochondrial and ER function. These effects align with recent studies suggesting another glycerolipid hydrolase DDHD2^16^ acts on both neutral lipids and phospholipids to support neuronal lipid metabolism and organelle function^10,14,84^, though DDHD2 does not localize to LD^10,74^. A role for ATGL in regulating phospholipids further aligns with phospholipid remodeling due to loss of ATGL in other cell types and systems^41,85,86^, and with known effects of ATGL on ER and mitochondrial homeostasis in endothelial cells^87^, cardiomyocytes^88^, and pancreatic beta cells^89^. ATGL therefore has broad effects on neuronal lipid distribution and organelle homeostasis. Indeed, the diverse cellular processes affected by ATGL likely explain why its loss in AgRP neurons has incomplete phenotypic overlap with precise manipulations of ER stress pathways^69,90^. Considering we show genes in addition to ATGL regulate neuronal LD in *Drosophila*, we propose a broad model in which LD support neuronal lipid distribution and organelle homeostasis to maintain appropriate neuron function under normal physiological conditions. Future studies will need to refine this model, however, to explain why neuronal loss of individual LD-regulatory genes cause distinct energy homeostasis phenotypes even when they have the same effect on neuronal LD abundance (*e.g*. dHSL and dATGL). Follow-up studies will also need to determine whether the relationship between LD abundance and neuron function is similar in other neurons.

Overall, our studies across flies and mice provide strong evidence that LD are a functionally significant organelle in neurons *in vivo* under normal physiological conditions. We show that LD regulation manages the tight coordination between lipid supply and demand to ensure the accurate distribution and utilization of lipids in neurons. Given that lipids support many aspects of neuron function (*e.g*. activity, morphology), and neuronal lipid metabolism is dysregulated in common diseases such as Alzheimer’s disease, our findings have broad implications for our understanding of lipid regulation in neurons across physiological and pathological contexts. Our discovery of sex differences in the cellular and functional consequences of LD dysregulation in neurons also highlights the importance of biological sex effects on neuronal lipid metabolism. More broadly in the brain, deeper insight into the regulation of neuronal lipid metabolism will provide key information to understand the close metabolic coupling between glia and neurons. Indeed, recent studies show that oxidative stress, mitochondrial dysfunction, excitotoxicity, or tauopathy in neurons promotes LD formation in glia^17,18,80,91,92^. Glial LD formation in these contexts is triggered by the neuron-to-glia transfer of lipids^17,18,80^, which are metabolized and neutralized in glia via ꞵ-oxidation^17^. Thus, shedding light on the mechanisms underlying LD formation in neurons will help identify additional mechanisms by which we can resolve defects in the neuron-glia exchange of lipids that is observed in disease^17,18,80^.

## METHODS

### C. elegans

#### Maintenance and strains

*C.elegans* were maintained as previously described^94^. Briefly, worms were maintained on standard NGM plates streaked with OP50 *Escherichia coli*. TU3311 (*uIs60 [unc-119p::YFP + unc- 119p::sid-1]*) were obtained from the *Caenorhabditis* Genetics Center (University of Minnesota, Minneapolis; RRID:WB-STRAIN:WBStrain00035055), which is funded by NIH Office of Research Infrastructure Programs (P40 OD010440). All experiments were performed at 20°C.

#### RNAi experiments

RNA interference (RNAi) treatment was performed by feeding *E. coli* HT115 containing an empty vector (EV) or *atgl-1* RNAi clone from the ORFeome RNAi library (Open Biosystems). Worms were transferred onto RNAi plates enriched with 1 mM isopropyl-ß-D-thiogalactopyranoside (IPTG) at day 1 of adulthood. The *atgl-1* RNAi clone was confirmed by sequencing.

#### Oil red O staining, imaging, and quantification

Oil red O staining was conducted as previously reported but by omitting the freeze-thaw steps and the MRWB-PFA permeabilization^95,96^. Briefly, Oil Red O stock solution was made at a concentration of 10 mM and balanced for at least 2 days on a rocker at RT. The working solution was freshly made at a concentration of 6 mM and filtered. Age- synchronized day 9 adult worms were dehydrated in PBS/isopropanol 60%/0.01% Triton-X for 15 min at RT and then stained overnight at RT with Oil red O working solution. Worms were washed 3 times with PBS/0.01% Triton-X and mounted on slides with mounting media. Oil red O staining was visualized using Zeiss Fluorescent Microscope, using the brightfield, with a ×10 objective. Images were quantified using the Fiji (ImageJ; RRID:SCR_002285) software. Integrated density was used as the primary measure and the relative percentage of the signal was calculated. ORO stain was quantified in 55–153 animals per condition, over 3 different sets of experiments.

#### Bodipy staining

BODIPY™ 558/568 C12 (Invitrogen D3835) was used to perform vital staining in live worms. BODIPY™ was diluted at 5 µM in either EV or *atgl-1* RNAi bacterial suspension. 6-cm plates were streaked and dried in a laminar flow hood for immediate use. After being fed with RNAi (EV or *atgl-1*) from day 1 of adulthood, day 7 adult worms were fed with BODIPY™ 558/568 C12 RNAi suspension (EV or *atgl-1*) for 24h at 20°C, protected from light. The following day, the worms were transferred to an NGM agar plate streaked with OP50 (for non-fasting condition) or a non-streaked plate (for the fasting condition) for an additional 24h. Worms were mounted on slides with 2% agarose pads and immobilized using a 5 mM solution of levamisole diluted in M9. Staining was visualized using Zeiss Fluorescent Microscope, and quantified using the Fiji (ImageJ) software. Only the anterior gut is quantified, representing LD and fat accumulation in worms^97^.

### Drosophila

#### *Drosophila* rearing

Fly strains were maintained on a 12 h:12 h light:dark cycle at 22°C. Larvae were reared on cornmeal-sugar-yeast 2-acid medium ^27,98^ at a density of 50 larvae per 10 mL food. Male and female pupae were distinguished by the presence or absence of sex combs, respectively. Pupae eclosed into single-sex vials; adult flies were transferred into new vials every 2-3 days. For fasting adult flies were transferred to 0.8% agar (w/v) in 1X PBS for either 12 hr or 24 h. All experiments used 5- to 6-day-old male and female flies.

#### *Drosophila* strains

We used the following strains from Bloomington *Drosophila* Stock Center: *w^11^*^18^ *(3605), UAS- dHSL-RNAi* (65148), *UAS-dPLIN2-RNAi* (32846), *UAS-dDIESL-RNAi* (67895), *UAS-dSREBP-RNAi* (34073), *UAS-dDGAT1-RNAi* (65963), *y^1^,v^1^;P{y[+t7.7]=CaryP}Msp300[attP40]* (36304), *y^1^,v^1^;P{y[+t7.7]=CaryP}attP2* (36303), *UAS-reaper* (5823), *UAS-NaChBac* (9468), *UAS-Lst-RNAi* (60400), *UAS-kir2.1* (6595), *elav-GAL4 (458)*. We used the following strains from Vienna *Drosophila* Resource Center: *UAS-dATGL-RNAi* (37880), *UAS-dHSL-RNAi*#2 (109336), *UAS- dPLIN1-RNAi* (30884), *UAS-dPLIN1-RNAi*#2 (106891), *UAS-dPLIN2-RNAi*#2 (102269), *UAS- dAGPAT-RNAi* (48593), *y,w^11^*^18^*;P{attP,y^+^,w^3’^}* (60100). We received *UAS-GFP-LD (2.6)* and *UAS-GFP-LD (3.4)* from M. Welte, *UAS-mRFP* from D. Allan, *nsyb-GAL4* from M. Gordon, *Akh-GAL4* from J. Park, *Akh^rev^* and *Akh^A^* from Ronald Kuhnlein. All GAL4 strains were backcrossed into *w^11^*^18^ background for 10 generations.

#### Body fat and storage measurements

Five- to six-day-old flies were anesthetized, weighed, and snap-frozen at -80°C (storage time <4 weeks). Triglyceride concentration was measured as described^99,100^, with minor modifications as in^27^. Percent body fat was calculated using triglyceride concentration and body weight. One biological replicate consists of a group of 5 flies; each experiment contained 4 biological replicates and every experiment was repeated twice (n=8).

#### Whole-brain and APC sample and slide preparation

For both whole-brain and APC dissections, adult flies were anesthetized briefly and individually dissected in cold PBS. Tissues were fixed at room temperature for 40 min in 4% PFA (Electron Microscopy Sciences 15710) and washed twice in 1 ml cold PBS. Samples were incubated in Hoechst 33342 (1:500 in PBS; Invitrogen H3570) for 30 min, washed again in PBS, and mounted in a saturated sucrose solution (70% w/v). Slides were kept at 4°C until imaging. Imaging was carried out no more than 30 h after mounting. N=10-25 for all APC and whole-brain samples.

#### Protein synthesis assay and Akh quantification

Akh was detected in the APCs using an anti-Akh primary antibody as described^57^ with AlexaFluor488 (1:200) goat anti-rabbit secondary antibody. The Click-iT Plus OPP Protein Synthesis Assay kit was used to measure nascent protein synthesis in APC. Briefly, APC-containing tissue marked by mRFP was dissected in *Drosophila* Schneider’s medium and immediately transferred to a solution containing OPP reagent (1:2000 in Schneider’s medium), and incubated for 30 min at room temperature. OPP solution was removed and the APC-containing tissue was washed 3 times with Schneider’s medium and fixed with 4% PFA. OPP was detected according to manufacturer’s instructions (Molecular Probes C10456).

#### Fluorescent dyes

To measure APC mitochondrial mass, tissue containing the APC was isolated in cold PBS and transferred to a solution with MitoView Green (1:500 in PBS; Biotium 70054) for 30 min before fixing with 4% PFA. Subsequent fixation and/or washes and mounting as described above.

#### Image acquisition and quantification

All tissues were imaged within 48 h of mounting; images were acquired on a Leica TSC SP5 inverted confocal microscope system. Images were processed using Fiji image analysis software^101^. LD counts and mitochondrial number (MitoView dye) were obtained using a custom Fiji counting macro. Briefly, a three-dimensional image containing lipid droplets marked by GFP-LD were first cleaned using “Median (3D)” function in Fiji^101^. Background was then removed using the ’Subtract Background’ function with a size parameter of 8. To ensure that each image has consistent intensities for the lipid droplets, the intensity of the processed image was standardized by applying a look up table (LUT); values for the LUT applied were the display range found using the “Enhance Contrast” function with the saturated parameter set to 0.1 on a maximum projected image of the median filtered and background-removed image. The lipid droplets were detected in the processed image using “3D Maxima Finder” function with “radiusxy=3 radiusz=6 noise=150” as arguments. Fluorescence for the Xbp-1 ER stress reporter, OPP assay, and Akh peptide levels in the APC was quantified by measuring the sum of fluorescence across 3 optical sections of the APC; fluorescence for Xbp-1 reporter, and OPP assay was normalized to APC size. For all image quantification, one biological replicate represents APC from one adult fly.

#### RNA sequencing

One biological replicate consisted of 30 *Drosophila* brains; we collected three replicates per sex and per genotype. Brains were dissected individually and transferred to Trizol; all brains per replicate were pooled into 1 mL Trizol and stored at -80°C. RNA was isolated according to manufacturer’s protocol (Thermo Fisher Scientific 15596018). Sample quality and RNA sequencing was performed at the UBC Biomedical Research Center Sequencing Core, as previously described^102,103^. Differences in gene expression were identified using DESeq2 R package^104^.

#### *Drosophila* Lipidomics

One biological replicate consisted of fifty *Drosophila* brains, and we collected four replicates per sex and per genotype. Brains were dissected individually and transferred to 80% LC-MS grade methanol; all brains per replicate were pooled into 1 ml 80% LC-MS grade methanol and stored at -80°C. The brains were sonicated in an ice-water bath for 30 minutes and stored at -20°C for 4 h for protein precipitation. The solvent was evaporated, and the lysate was resuspended in 375 μL of 80% ice-cold methanol in water. 1 mL of methyl tert-butyl ether was added, and the mixture was shaken for 5 min. Phase separation was induced by adding 275 μL of water. The upper layer containing lipids was transferred to a new tube, dried, and reconstituted in acetonitrile/isopropanol (1:1, v/v) at a volume proportional to bicinchoninic acid assay protein quantification results. Species were only considered differentially regulated if fold-change >0.3 different from controls and unadjusted *p*<0.05. N=4 samples per sex and per genotype. Because we had two control groups, lipid species in *Drosophila* were only considered differentially regulated by neuronal *dATGL* loss if their abundance was different from both control groups.

### Mouse neurons in culture

#### GT1-7 and N46 hypothalamic neurons

GT1-7 cells (a generous gift from Dr Pamela Mellon; RRID:CVCL_0281) and N46 cells (Cedarlane, #CLU138; RRID:CVCL_D45) were grown in DMEM (4.5 g/L glucose, Gibco™, #11965092) containing 10% FBS (Wisent™, #10437-028), 100 mM Sodium pyruvate (only for GT1-7 cells, Gibco™, #11360-070) and 5000 U/mL of Penicillin/Streptomycin (P/S; Gibco™, #15070-063) until 80% confluence.

#### Primary hypothalamic cultures

Brains from NPY-GFP or POMC-GFP P1-P2 mouse pups were harvested and the mediobasal hypothalamus (MBH) was dissected in 10 mL of Lebovitz-15 media (Gibco, #11415-064) and centrifuged at 2000g for 2 min (see description of strains in mouse studies). The pellet was then digested at 37 °C for 15 min using 4.5 U/mL lyophilised papain (Worthington Biochemical, #LS003120), 5 mg/mL D-glucose (Sigma, #G7528-1KG), 0.2 mg/mL L-cystein (MP, #101444) 0.2 mg/mL BSA (Multicell, #800-095-EG) and 20 U/mL DNAse I (Worthington Biochemical, #LS006333) in PBS 1X and centrifuged at 2000g for 2 min. The pellet was then resuspended in Neurobasal A media 1X (Gibco, #10888-022) with 2% B-27 supplement (Gibco, #17504-044), 2% FBS (Wisent™, #10437-028), 1% P/S (Gibco™, #15070-063) and 1% GlutaMax (Fisher, #35050061) and filtered using a 70 μM cell-strainer. Neurons were then allowed to grow on coverslips for 7 days in Neurobasal A media with 2% B-27, 2% FBS, 1% P/S and 1% GlutaMax and 2 μM Cytosine beta-D-arabinofuranoside (AraC; Sigma, #C1768-100MG) to inhibit glial cell growth. Neurons were fixed using 4% paraformaldehyde, stained using 1:1000 HCS LipidTox Neutral Red Dye (Fisher, #H34476) for neutral lipids and 1:1500 Hoechst 33342 for nuclei (Invitrogen, #H3570) and imaged using the Zeiss fluorescent microscope (Carl Zeiss AG).

#### High throughput LD imaging

GT1-7 and N46 cells were plated on 96-well plates (PerkinElmer, #6055302). Cells were treated with DMEM (Multicell, #319-060-CL) in 1% FBS (Wisent™, #10437-028), 10 mM glucose and 0.25 mM oleate (Nu-chek Prep, #S-1120) in 0.27% bovine serum albumin (BSA) (Multicell, #800- 095-EG), 50 μM Atglistatin (Cedarlane, #HY-15859), 5 μM Forskolin (Sigma, #344282-5MG), 100 nM DGAT1 inhibitor A-922500 (Sigma, #A1737-1MG) or 0.1% DMSO for 2.5 or 24 h as described in the figure legends. Cells were fixed using 4% paraformaldehyde and neutral lipids were stained using 1:500 5 mM Bodipy 493/503 (Sigma, #D3922) while nuclei were stained using 1:1500 Hoechst 33342 (Invitrogen, #H3570). Plates were imaged using Operetta High Throughput screening system and analyzed using Harmony High-Content Imaging and Analysis Software. For analysis, one experimental N consisted of the average of 6 wells in which, in every well, 9 areas were randomly selected for quantification.

#### Mass spectrometry (MS)-based Lipidomics

Lipidomic studies were performed on GT1-7 neurons and analyzed by the Montreal Heart Institute Metabolomic Core Facility. MS-based targeted and untargeted lipidomics required 10 million and 5 million cells per condition respectively. Protein quantification was performed using a Bradford assay for MS normalization.

*1- Profile of esterified fatty acids analysis (triglyceride fraction) using GC-MS*

Briefly, GT1-7 and N46 cells were treated with DMEM containing 1% FBS, 10 mM glucose and 0.1% DMSO or 50 µM ATGListatin for 24h, counted, harvested in screw-cap tubes of 10 million cells and flash frozen using liquid nitrogen. Quantitative profiling was performed using gas chromatography-MS (GC-MS) analysis using as previously described^105^. In brief, cells were incubated in a mixture of chloroform/methanol (2:1) and supplemented 1:4 with 0.004% butylated hydroxytoluene (BHT) overnight at 4 °C before being filtered, dried under nitrogen gas, and re- suspended in a mixture of hexane/chloroform/methanol (95:3:2). The triglycerides fraction was obtained after separation on an aminoisopropyl column, dried under nitrogen gas following the addition of specific internal standards and resuspended in hexane/methanol with 0.004% BHT. Esterified FA were analyzed as their methyl esters (FAMES) following a direct trans-esterification with acetyl chloride and neutralized by incorporating 6% potassium carbonate. Thereafter, the upper hexane phase is injected onto a 7890B GC coupled with a 5977 mass selective detector (Agilent Technologies, Santa Clara, CA, USA) equipped with a capillary column (J&W Select FAME CP7420; 100 m × 250 µm inner diameter; Agilent Technologies, Santa Clara, CA, USA). The analysis was operated in positive chemical ionization mode and ammonia was used as reagent gas. Chromatographic conditions were fixed as follows: injection at 270°C in a split mode, high-purity helium used as carrier gas and a temperature gradient beginning at 190°C for 25 min and increased by 1.5 C/min up to 236°C. FA were analyzed as their [M+NH3]+ and concentrations calculated using internal standards and standard curves.

*2- Untargeted Lipidomics*

GT1-7 neurons were plated on glass plates and treated with DMEM containing 1% FBS, 10 mM glucose and 0.1% DMSO or 50 µM ATGListatin for 24h, counted, harvested in 15 mL Falcon tubes of 5 million cells and flash frozen using liquid nitrogen. Lipid species were then extracted and analyzed as previously described^106,107^. Briefly, samples were injected (from 0.6 to 1.1 µl according to protein concentration) into a 1290 Infinity high resolution HPLC coupled with a 6530 Accurate Mass quadrupole time-of-flight (LC-QTOF, Agilent Technologies) system equipped with a dual electrospray ionization (ESI) source. Elution was assessed using a Zorbax Eclipse plus column (C18, 2.1 x 100 mm, 1.8 µm, Agilent Technologies) maintained at 40 °C using an 83 min chromatographic gradient of solvent A (0.2% formic acid and 10 mM ammonium formate in water), and B (0.2% formic acid and 5 mM ammonium formate in methanol/acetonitrile/methyl tert-butyl ether [MTBE], 55:35:10 [v/v/v]). Samples were analyzed in positive ionization mode. MS data was processed using the Mass Hunter Qualitative Analysis software package (version B.06.00, Agilent Technologies Inc.) and an in-house bioinformatics pipeline leading to a list of MS signal features characterized by a mass, a retention time, and a signal intensity. Lipid annotation was performed using data alignment with an in-house database previously validated using MSMS. MSMS was used to confirm the latter annotations and identify, when possible, additional unknown MS features.

#### Metabolomics

GT1-7 cells were plated to 80% confluence on T25 flasks and treated with DMEM containing 1% FBS, 10 mM glucose and 0.1% DMSO or 50 μM ATGListatin for 24h, after which flasks were rinsed and flash-frozen in liquid nitrogen. Quantification of metabolic species was performed by the CRCHUM Metabolomics core facility, as previously described ^108^. Briefly, samples were quantified using Liquid Chromatography with tandem mass spectrometry (LC-MS-MS), normalized to protein levels (Bradford protein assay), and a blank condition (no cells) was subtracted.

#### Mouse studies

All animal care and experimental procedures were conducted in accordance with guidelines of the Canadian Council on Animal Care and approved by the institutional animal care committee at CRCHUM (protocol #CM19018TAs and CM230041TAs). Mice were housed on a 12h dark-light cycle (dark from 10am to 10pm) at 21-23°C, in a pathogen free environment. Standard irradiated chow diet (Teklad) and water were provided *ad libitum* or otherwise mentioned. For all studies, age- and sex-matched littermates were used and individually housed and animals were in experimental designs from 8 to maximum 20 weeks old.

ATGL^fl/fl^ mice in which exon 1 is flanked by loxP sequences were kindly donated by Dr Grant Mitchell ^109^ and maintained at least 6 generations on the C57BL/6J genetic background (C57BL/6J, 000664; RRID:IMSR_JAX:000664). AgRP-IRES-Cre were previously generated ^110^ and obtained homozygous from the Jackson laboratory [Agrp^tm1(cre)Lowl/J^, 012899] (129S6/SvEvTac background)(RRID:IMSR_JAX:012899). NPY-GFP reporter mice were obtained from the Jackson laboratory [NPY^hrGFP(1Lowl/J)^, 006417] (B6/FVB-Tg background)(RRID:IMSR_JAX:006417). Male NPY-GFP [B6.FVB-Tg(Npy-hrGFP)1Lowl/J, 006417] and POMC-eGFP mice [C57BL/6J-

Tg(Pomc-EGFP)1Low/J, 009593](RRID:IMSR_JAX:009593) were purchased from The Jackson Laboratory (6–10 weeks old) and bred with C57BL/6J WT females from the same genetic background to produce experimental animals.

#### Metabolic challenges

Naive WT (C57Bl/6J) male mice were purchased at 7-8 weeks old from Jackson for the expression profile experiments. They were acclimated to a reversed cycle 2 to 3 weeks before any experiments. Fasted mice were food deprived for 16h starting during the second half of the dark cycle. Cohorts of animals were exposed for 24h to 21 °C (controls), 30°C (thermoneutrality) or 4°C (cold) in CLAMS metabolic cages.

ATGL KO in ARC neurons (ARC^ATGL^CTL vs ARC^ATGL^KO)

Eight- to 9-week-old ATGL^fl/fl^ males and females underwent stereotaxic surgery to knock out (KO) invalidate ATGL in the arcuate nucleus (ARC) using Cre-expressing viruses as previously described ^111^. Briefly, ATGL^fl/fl^ mice were kept under anesthesia with isoflurane and received bilateral viral injections of either AAV9-hSyn-Cre-2A-tdTomato-SV40pA (Viroveck, 2.18E13 vg/ml) or AAV9-hSyn-tdTomato-SV40pA (Viroveck, 2.24E13 vg/ml). 200 nL/side were simultaneously injected (0.5 nL/sec) using neurosyringes (Hamilton, #65457-01) placed at a 10° angle, to the following coordinates AP: bregma-1.4 mm ; lateral: sinus+1.2 mm ; depth: dura-5.9 mm. Five min after the end of the injection, syringes were removed, the wound closed and mice recovered in thermoneutrality incubators for at least 2h. Animals were kept at least 4 weeks after surgery before experimentation. Accuracy of ARC viral injections were validated using tdTomato. Only mice harboring more than 20% reduction of ATGL expression in the ARC were included in the study.

ATGL KO in AgRP neurons (AgRP^ATGL^CRE vs AgRP^ATGL^KO):

The Cre-Lox strategy was used to excise exon 1 of the ATGL floxed gene in AgRP neurons. The Cre locus was maintained heterozygous, to avoid potential Cre toxicity. Briefly, AgRP^Cre/+^ mice were bred with NPY-GFP mice. Male AgRP^Cre/+^:NPY^GFP/+^ were bred with female ATGL^fl/fl^ mice to generate AgRP^Cre/+^:NPY^GFP/+^:ATGL^fl/+^ mice which were then crossed with ATGL^fl/+^ or ATGL^fl/fl^ to generate experimental mice : AgRP^Cre/+^:ATGL^+/+^ (AgRP^ATGL^CRE) and AgRP^Cre/+^:ATGL^fl/fl^ (AgRP^ATGL^KO). Mice carrying or not the NPY^GFP/+^ transgene were included in experimentations. Mice harboring significant ectopic ATGL recombination (detected by genomic qPCR) were not included in experimentations.

#### Metabolic cages (CLAMS)

Respiratory quotient (RQ), energy expenditure (EE), food consumption were monitored using indirect calorimetry in Comprehensive Lab Animal Monitoring System metabolic cages (CLAMS, Columbus Instruments International; RRID:SCR_016718se) as previously described^112^. Animals were single-housed in CLAMS apparatus in a dark/light cycle matching their housing conditions during 24h for acclimation, followed by measurements. Energy expenditure was normalized by lean mass. Fatty acid oxidation (FAOx) was calculated from RQ and EE (not normalized by lean mass) as FAOx = (EE X (1-RER))/0.3.

#### Feeding behavior analysis

The CLAMS system weighs hoppers with food (± 0.01g) every second and detects “not eating” when weight is stable and “eating” if unstable. Single feeding events (bouts) are calculated as the weight difference between “eating” and “not eating” events. They are recorded as feeding vectors with a start time, their duration (s), and the amount of food consumed (g). Data for single mice were extracted using Oxymax software then analyzed and compiled with MATLAB (MathsWorks© R2021a; RRID:SCR_001622). As previously reported, meals consist of the sum of single feeding events (bouts) separated by an inter-meal interval (IMI)^113–115^. As feeding bouts have three characteristics (start time, duration, size), three parameters had to be determined to define a meal. A meal is the sum of bouts > 0.03 g and > 10 sec, occurring within < 5 min. On a defined period of time (24 h for example), following parameters were thus calculated using MATLAB program: cumulative food intake (the sum of all meals (g)), and the average meal size (g), with the average IMI (s), to calculate satiety defined as the ratio of meal size (g) / IMI(s).

#### Body Composition

Body composition (fat and lean mass) was measured by magnetic resonance imaging (echoMRI). Brown adipose tissue (BAT), inguinal, intraperitoneal (perigonadal) and subcutaneous (inguinal) fat pads were collected and weighed at sacrifice.

#### Glucose Tolerance Test

Mice were food deprived 5 h before the test. A bolus of glucose (Dextrose, 1.5 g/kg) was administered via an intraperitoneal injection, and glycemia was measured from blood sampled from the tail vein using an Accu-chek Performa glucometer at T0 (before injection), 15, 30, 60, and 90 min. Blood samples were collected via a capillary for insulin assays.

#### Plasma Insulin

Insulin assays were performed by the cell physiology platform of the CRCHUM using commercially available ELISA kits.

#### Tissue Collection

Fresh brain microdissection and tissue collection was done after mice were deeply anesthetized with isoflurane. Tissues were weighed, flashed frozen and stored at -80°C. For cytomorphologic experiments, perfusions were performed. Mice were deeply anesthetized with excess Ketamin/xylazin and transcardially perfused with 1x PBS followed by 4% PFA. Brains were extracted, post-fixed for 2 h in 4%PFA, put in sucrose overnight and stored at -80°C. Using a microtome (Leica, SM2000R), 30μm brain coronal sections were made and kept in antifreeze at -20°C before any immuno-histochemistry (IHC) or hybridization *in situ* (RNAScope).

#### Gene Expression

Total RNA from frozen tissue was extracted using the Trizol method as previously described^116^. Total RNA concentration and purity were determined using the Nanodrop 2000. cDNA synthesis and qPCR were performed as described^116^. Briefly, 1 µg of total RNA was retro-transcribed with M-MuLV reverse transcriptase (Invitrogen, #28025013) using random hexamers and diluted 1:10 prior to quantification by qPCR (QuantiFast SYBR Green PCR kit, Qiagen, #28025013) using a Corbett Rotor-Gene 6000 (Qiagen) with primers (1 µM) described in **Table 2**. qPCR was quantified using the standard curve method. Gene expression was normalized to the expression of a stable housekeeping gene or the geometric mean of many (determined by NormFinder). Gene expression is represented as fold change from mean of control groups, after normalization.

**Table 1.**
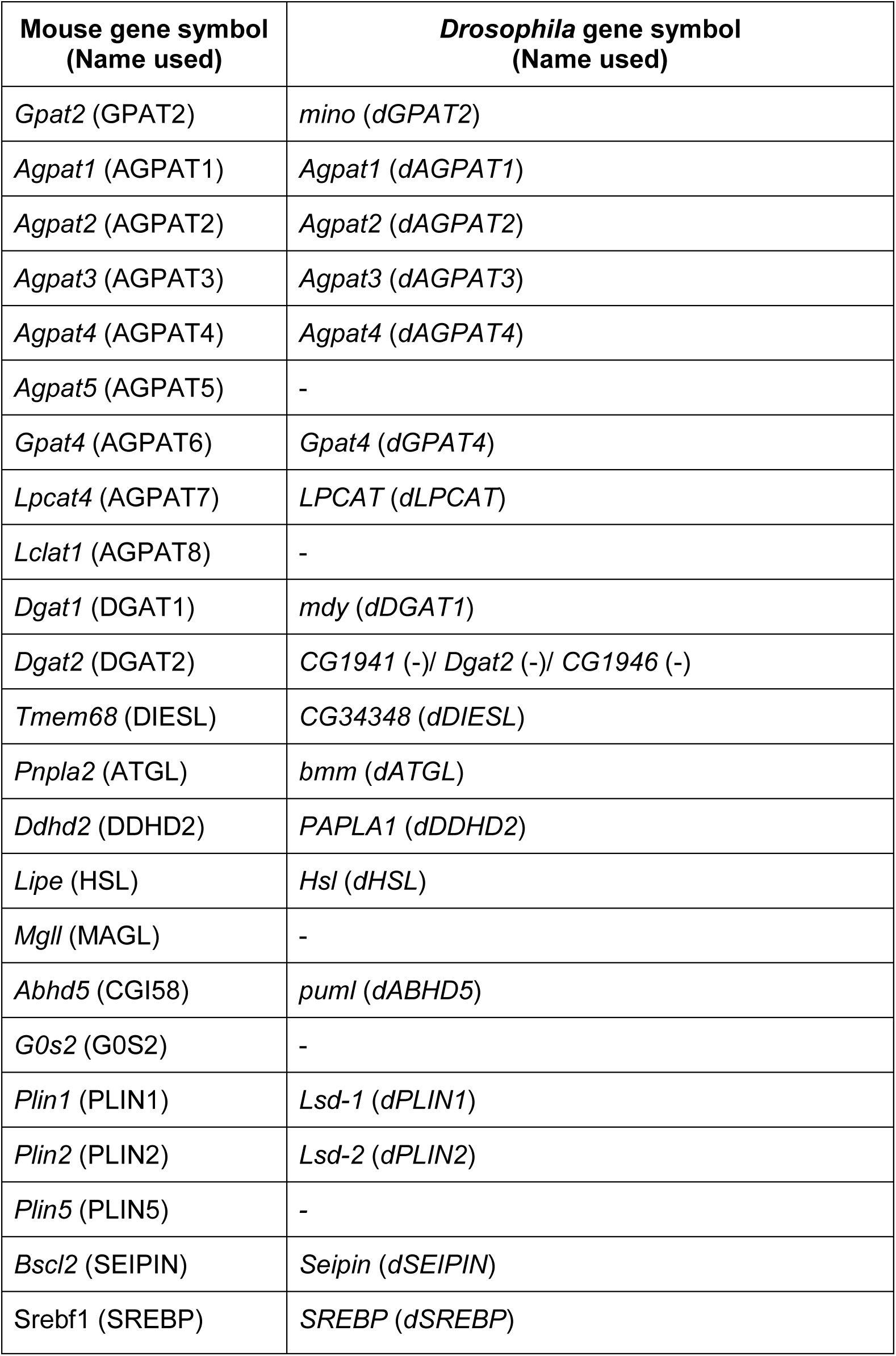
Species equivalence of genes and lipid droplet-associated proteins.

**Table 2.**
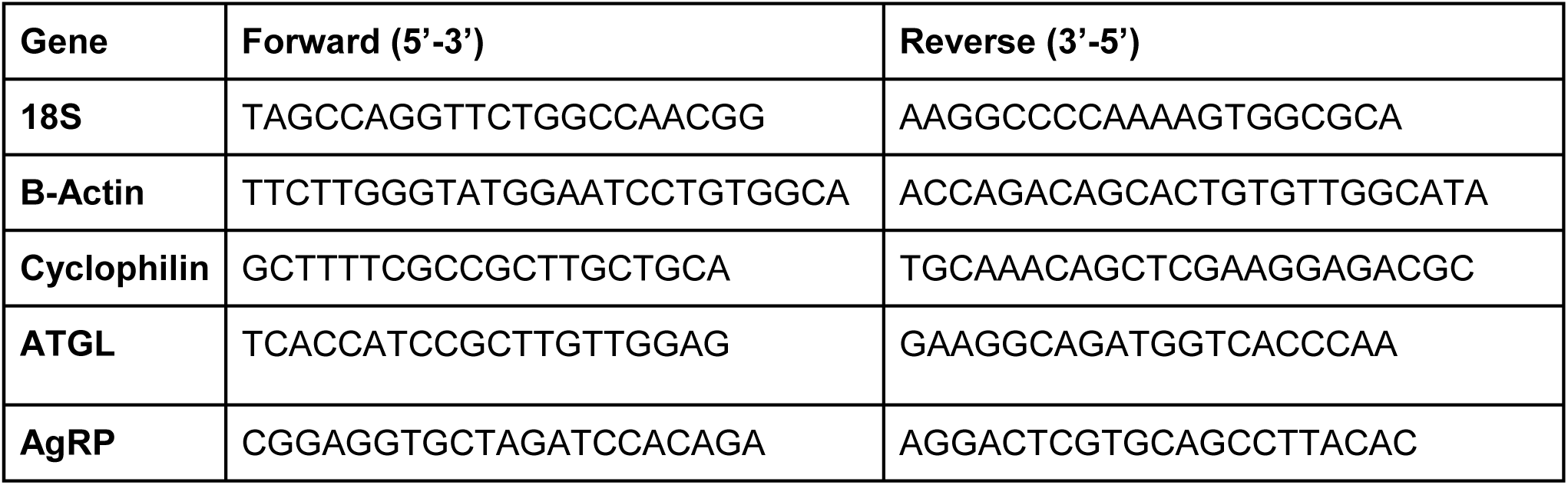
qPCR primers.

#### RNAScope

Brain sections (30 µm) were washed with 1X PBS, mounted on microscope glass slides, and dried for 30 min at 60°C. Following the company’s instructions (ACD, Multiplex Fluorescent V2 Assay, #323100), on the first day, all sections were dehydrated, incubated with hydrogen peroxide, steamed for 5 min with antigen retrieval, and treated with Protease III for 15 min. Pnpla2 (#469441-C1) and AgRP (#400711-C2) probes were incubated 2 h at 40°C as indicated (c2 probe diluted at 1:50 in the c1-probe). A negative control brain section was done for every sample using a negative probe (ACD, #320871). Slides were kept ON, in 5x Saline Sodium Citrate protected from light, at RT. On the second day, after washing in wash buffer, sections were amplified 3 times as indicated by the company. Fluorescent signal was developed using opal dye diluted at 1:1500, sections were counterstained with DAPI and mounted with ProLong Gold Antifade Mountant (eLife, #P36930) and stored at 4°C. Zeiss fluorescent microscope (Carl Zeiss AG; RRID:SCR_013672) with an Apototome was used for imaging. Quantification was done using Fiji software^101^.

#### Immunohistochemistry

Brain sections (30 µm) were successively permeabilized 5 min using Triton solution (0.01X), blocked 2 h with 2% goat serum, incubated with primary antibodies ON at 4°C and then 2 h at RT with secondary antibodies. Sections were mounted with DAPI incorporated in the mounting media (Vectashield, VECTH1200) and imaged with a Zeiss fluorescent microscope (Carl Zeiss AG; RRID:SCR_013672). Primary antibodies: anti-AGRP, 1:500 (EPR18155-110, Abcam #254558, RRID:AB_3076273); anti-GFP, 1:500 (life technologies 33-2600; RRID:AB_2533111). Secondary antibodies: Alexa Fluor 546–goat anti-rabbit IgG (A-11035; RRID:AB_2534093) and Alexa Fluor 488–goat anti-mouse IgG (A-11001; RRID:AB_2534069) (1:1000; Life Technologies)

#### Electrophysiology

<u>Stereotaxic surgeries:</u> 7-week-old AgRP^ATGL^CRE and AgRP^ATGL^KO mice underwent stereotaxic surgery to selectively induce the expression of the red mCherry fluorescent protein in AgRP neurons. Briefly mice were kept under anesthesia with isoflurane and received bilateral viral injections of AAV2-CAG-DIO-mCherry-WPRE-bGHpA (Viroveck, 1.31E13 vg/ml). 200nL/side were simultaneously injected (0.5nl/sec) using neurosyringes (Hamilton, #65457-01) placed at a 10° angle, to the following coordinates AP: bregma-1.4 mm; lateral: sinus +1.2 mm; depth: dura-5.8 mm. 5 min after the end of the injection, syringes were removed, the wound closed and mice recovered in thermoneutrality incubators for at least 2h.

<u>Arcuate nucleus slices:</u> Coronal arcuate nucleus slices (300 µm thick) were obtained from AgRP^ATGL^CRE or AgRP^ATGL^KO mice 3 to 4 weeks after the surgeries. Animals were deeply anesthetized with isoflurane and brains were rapidly removed. Then the brain was cut with a vibratome VT1200 (Leica, Germany) in ice-cold N-methyl-D-glucamine (NMDG) cutting ACSF containing (in mM): 119.9 NMDG, 2.5 KCl, 25, 1 CaCl2, 1.4 NaH2P04, 26 NaCO3 and 20 D-glucose saturated with 95% O2 and 5 CO2. Slices containing ARC were transferred to 95% O2 and 5 CO2 saturated NMDG solution at 33°C followed by ASCF containing (in mM): 130 NaCl, 2.8 KCl, 1.25 NaH2PO4, 1.2 MgCl2, 2.5 CaCl2, 26 NaCO3 and 2.5 D-glucose for 1 h at room temperature prior to recordings.

<u>Electrophysiology:</u> Slices were transferred to a recording chamber where they were perfused with ACSF (2 ml/min). Whole-cell recordings were achieved from the soma of mCherry expressing ARC neurons. Borosilicate patch pipettes (2-5 MΩ) were filled with an internal solution containing (in mM): 105 K-gluconate, 30 KCl, 10 phosphocreatine, 10 HEPES, 4 ATP-Mg, 0.3 GTP-Tris, 0.3 EGTA (adjusted to pH 7.2 with KOH; 290-300 mOsmol). Recordings were made at room temperature. Data were acquired using a Multiclamp 700B amplifier (Molecular Devices) and digitized using a Digidata 1440A digitizer and pClamp/Clampfit 10.7 (Molecular Devices). Recordings were low pass-filtered at 2 kHz and digitized at 20 kHz. Access resistance (Ra) was 10-25 MΩ and regularly monitored during experiments, data were excluded if Ra variations were above 20% throughout the experiment. Input resistances (Rin) were calculated by measuring the slope of the linear portion of the current-voltage (I-V) curve (-120mV to -70mV). Spontaneous firing frequencies were obtained from I=0 current-clamp mode recordings for at least 2 min.

#### Electron Microscopy

<u>Tissue preparation:</u> Adult mice were injected and anesthetized with excess Ketamin/xylazin and transcardially perfused successively with cold solutions at a very slow rate: 50 mL PBS, 100 mL fixative solution (2% glutaraldehyde (EMS cat. #16320) + 4% paraformaldehyde (EMS cat. #15713-S), and 100 mL 4% PFA. Brains were extracted, post-fixed 30 min in 4% PFA, washed with PBS and freshly sliced using a vibratome (Leica VT1000S, vibration 10, speed 3) at 50 μm. Sections were stored in anti-freeze solution at -20°C until further use.

<u>Immunohistochemistry on Brain Slices:</u> Fifty μm sections containing the ARC of 8-week-old AgRP^ATGL^CRE or AgRP^ATGL^KO mice on the NPY-GFP background were processed for immunostaining against GFP prior to EM processing. Successively, sections were: quenched 5 min with 0.3% H2O2 (Fisher Scientific, Ottawa, Canada cat# H325500) in PBS, incubated 30 min in 0.1% NaBH4 in PBS, washed in PBS, blocked 1 h, RT in a solution containing 10% fetal bovine serum (FBS; Jackson ImmunoResearch Labs, Baltimore, USA cat# 005-000-121), 3% bovine serum albumin (Sigma-Aldrich, Oakville, Canada cat# A7906-500G), and 0.05% Triton X-100 (Millipore-Sigma, Oakville, Canada cat# X100-1L) in PBS. Next, sections were incubated overnight at 4°C in blocking buffer solution with the chicken anti-GFP antibody (1:5000; Aves Labs, Davis, USA cat# GFP-1020; RRID:AB_10000240). The following day, sections were washed with Tris-buffered saline (TBS; 50 mM, pH 7.4) and incubated for 90 min at RT with biotinylated donkey anti-chicken polyclonal secondary antibody (1:300; Jackson ImmunoResearch, Baltimore, USA, cat# 703-066-155; RRID:AB_2340355) in TBS containing 0.05% Triton X-100. Afterwards, sections were washed in TBS and incubated for 1h at RT in a 1:100 avidin-biotin complex (Vector Laboratories, Newark, USA, cat# PK-6100) solution in TBS. Lastly, 0.05% 3,3’-diaminobenzidine (Millipore Sigma, Oakville, USA, cat# D5905-50TAB) activated with 0.015% H2O2 diluted in Tris buffer (0.05 M, pH 8.0) was used to reveal the staining for 90s.

<u>Sample Preparation and Imaging by Transmission Electron Microscopy:</u> The protocol for EM sample preparation was recently detailed in ^117^. Briefly, brain sections were washed in phosphate buffer (PB) and incubated 1h in a solution containing equal volumes of 3% potassium ferrocyanide (Sigma-Aldrich, Ontario, Canada, cat# P9387) with 4% osmium tetroxide (EMS, Pensylvannia, USA, cat# 19190) in PB. After washing with 100% PB, 50%PB/50%ddH2O, 100% ddH2O for 5 min each, sections proceed to be incubated in a filtered (0.45 μm filter) 1% thiocarbohydrazide solution (diluted in MilliQ water; Sigma- Aldrich, Ontario, Canada, cat# 223220) for 20 min followed by a second incubation in 2% aqueous osmium tetroxide (diluted in MilliQ water) for 30 min. Then, sections were dehydrated in increasing concentrations of ethanol for 5 min each (2 × 35%, 50%, 70%, 80%, 90%, 3 × 100%) and washed 3 x 5 min with propylene oxide (Sigma- Aldrich, #cat 110205-18L-C). Sections were next embedded overnight at RT in Durcupan resin (20g component A, 20g component B, 0.6g component C, 0.4g component D; Sigma Canada, Toronto, cat# 44610). The following day, the resin-infiltrated sections were flat-embedded onto fluoropolymer films (ACLAR®, Pennsylvania, USA, Electron Microscopy Sciences, cat# 50425- 25) surrounded by resin and kept at 55°C in a convection oven for 72h to allow for polymerization. Next, tissue sections containing the region of interest were excised and glued to a resin block- face. For TEM imaging, using a Leica ARTOS 3D ultramicrotome, 2-3 74 nm thick sections were collected (4 levels, ∼ 8-10 μm apart) on copper grids (EMS, cat# G150-Cu) and imaging on TEM - 120kv JEOL 1400-Flash equipped with a 4k Gatan OneView digital camera. Using the software for GMS3 images of neurons were randomly acquired in the ARC at a resolution of 5 nm per pixel and exported as .tif file format.

#### Single cell RNA sequencing

The RNA Drop-seq single-cell dataset utilized for the preliminary assessment of transcriptional landscape in mice hypothalamic neurons was sourced from a public GEO dataset, using published annotations^38^. Re-analysis began from the merged raw counts file (mm10), and filtering of low-quality cells and doublets reproduced the original standard processing (800 < Features < 6000; Mitochondrial RNA (%) < 20). Data integration was performed in accordance with guidelines outlined in Seurat v4.1.0 vignettes^118^, but experimental groups were analyzed separately. Presented data focused on the initial three batches comprising 21 male mice (7350 cells), as well as the last batch, which contained two male (358 cells) and two female (676 cells) mice on a chow diet. Following standard Seurat v4 processing and re-clustering, original cell-type annotations were validated using the FindMarkers function with updated markers literature, such as the Azimuth database^118^. These annotations were used to subset neurons (Male=4445; Female=425) as well as specific AgRP (Male=977; Female=41) and POMC (Male=403; Female=22) populations for further analysis.

### Statistical analysis and data presentation

GraphPad Prism (10.2.2; RRID:SCR_002798) was used for the majority of data analysis and graph preparation. QQ plots, Shapiro-Wilk and Kolmogorov-Smirnov tests were used to test normality of residuals. If data was normally-distributed we used unpaired parametric tests (two-sided t-test and ANOVA); if data was not normally distributed we used an unpaired nonparametric test (Mann-Whitney test or Kruskall-Wallis). For ANOVA Tukey or Sidak post-hoc tests were used to perform multiple comparisons between groups. Two-way ANOVA were used to detect sex:genotype, genotype:time, and sex:diet interactions; three-way ANOVA were used to detect sex:genotype:time interactions.

### Data and code availability statement

Fly lipidomic and transcriptomic data are available in Supplemental tables. The other datasets generated and/or analyzed in the current study are available from the corresponding authors (without any restrictions) on reasonable request. Single-cell RNA seq and Drop-seq data (mouse neurons) are available at GEO accession codes GSE90806 and GSE93374 respectively. RNA-seq data (fly brains) are available at GEO accession code GSE270119.

## ACKNOWLEDGEMENTS

We thank Dr. Michael Welte for the *UAS-GFP-LD* strains^28^ and Dr. Jae Park for the *Akh-GAL4* strain and anti-Akh antibody^57^. Stocks obtained from the Bloomington *Drosophila* Stock Center (NIH P40OD018537) were used in this study. We thank the TriP at Harvard Medical School (NIH/NIGMS R01-GM084947) for providing transgenic RNAi fly stocks and/or plasmid vectors used in this study. Transgenic fly stocks and/or plasmids were also obtained from the Vienna *Drosophila* Resource Center (VDRC, https://stockcenter.vdrc.at/control/main). We acknowledge critical resources and information provided by FlyBase^119^. This study was supported by operating grants to E.J.R. from the Canadian Institutes of Health Research (CIHR, PJT-153072 and PJT- 183786), Michael Smith Foundation for Health Research (16876), a CIHR Sex and Gender Science Chair in Genetics (GS4-171365), and infrastructure purchased with funds from the Canadian Foundation for Innovation (JELF-34879). We acknowledge the UBC Biomedical Research Center for support with our *Drosophila* RNA sequencing dataset. We acknowledge that our research takes place on the traditional, ancestral, and unceded territory of the Musqueam people; a privilege for which we are grateful. We are grateful to Dr Grant Mitchell (U. Montréal) for the ATGL-floxed mice. We thank Colby Sandberg and Mohammadparsa Khakpour (U. Victoria) for processing EM samples. We are grateful to Dr Xavier Fioramonti (U. Bordeaux) for his help in optimizing hypothalamic electrophysiology studies. We thank the cellular imaging, small animal phenotyping and imaging, cell physiology, and the metabolomics core facilities at CRCHUM. This work was supported by grants to TA from the National Natural Sciences and Engineering Research Council (NSERC, RGPIN/04798), the CIHR (PJT153035) and by a grant to MET from the Canadian Foundation for Innovation (CFI, 39965). MT, SF, MR, CMR and TA were supported by a salary award from Fonds de Recherche Québec-Santé (FRQS). MET holds a Canada Research Chair (Tier II) in *Neurobiology of Aging and Cognition*. RM and DM were supported by doctoral fellowships from FRQS, Diabète Québec, Université de Montréal, the Neuroscience Department and CRCHUM. AL was supported by doctoral fellowships from ALS Canada and FRQS, and a MITACS postdoctoral fellowship. SA and JV were supported by MSc and doctoral fellowships from CIHR, respectively.

## AUTHOR CONTRIBUTIONS

***Drosophila***. CJM, JDF, LWW, NY, SP performed *Drosophila* fat storage and fat breakdown assays, CMC and CJM performed *ex vivo* and immunohistochemistry experiments on *Drosophila* APCs; CJM, SH, NS dissected *Drosophila* brains for LD counts. CJM, SH, CFC dissected samples for *Drosophila* RNAseq and lipidomics. TH and AH performed unbiased lipidomics and CFC, YHX, JJYX, CASC and GF analyzed and prepared figures for *Drosophila* lipidomics data. CMC and CJM contributed to figure design, CMC performed all statistical analysis, prepared all Supplemental Tables and Files, and finalized all *Drosophila* figures and data. CJM, CMC, JDF, CFC and EJR conceptualized and interpreted data from *Drosophila* experiments. ***C. elegans* and Mouse**. AL, RM and JAP carried out experiments in *C. elegans*. DM and RM performed studies on cultured mouse neurons. RM, DM, DR, KB helped with mouse colonies, genotyping, RNAscope, histology, qPCR, stereotaxic injections and phenotyping studies. CD and MR carried out LC-MS/MS for lipidomics. RM, JV, PK, MET and BL carried out EM studies. SA and MT performed scRNAseq data analysis. BR, LRDH, AB and CMR carried out electrophysiological recordings and data analysis. RM, DM, AJP, JV, MET, BL, MR, SF, CMR and TA contributed to conceptualization, experimental design for cultured neuron and mouse studies, data analysis and interpretation. **Manuscript.** RM, DM, TA, EJR drafted manuscript text; RM, DM, BL, CMC, CMR, SF, TA, EJR edited manuscript.

## SUPPLEMENTAL FIGURES

**Figure S1.**
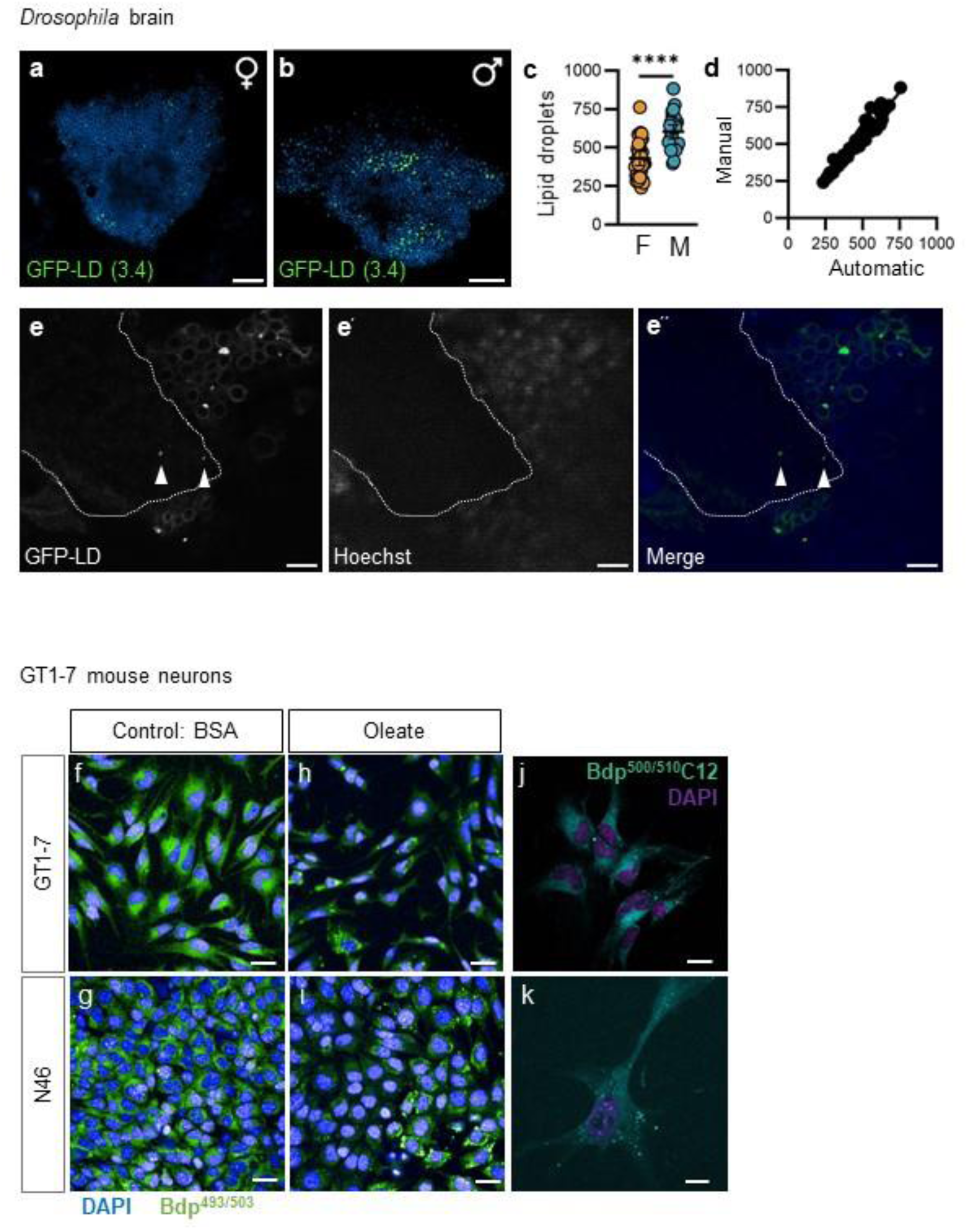
Neuronal lipid droplets are present under normal physiological conditions in flies and cultured hypothalamic neurons. **a,** Z-projection of confocal images from mushroom body of 5-day-old female and **b**, male animals with pan-neuronal expression of an independent *UAS-GFP-LD(3.4)* line. Green punctae represent neuronal lipid droplets (LD). (Scale=20 µm). **c**, Number of mushroom body LD in female and male brains in *elav>GFP-LD(2.6)* animals. Student’s t-test, \*\*\*\**p<*0.0001. **d**, Graph of LD number in *elav>GFP-LD(2.6)* females showing correspondence between manual LD counts and LD counts derived from our automatic counting script (R^2^=0.9405). **e**, Z-projection of confocal image of the mushroom body in an *elav>GFP-LD(2.6)* male. Hoechst indicates region with neuronal cell bodies; Hoechst-negative region indicates region with no cell bodies. Arrowheads indicate GFP-positive punctae corresponding to LD within axons. Scale=5 µm. **f,h**, Representative images of Bodipy^493/503^-stained LD in GT1-7 and **g**,**i**, N46 neurons. **f,g**, vehicle (BSA) and **h,i**, 24h Oleate treatment. Scale=40 µm. **j,k**, GT1-7 neurons treated with Bodipy^500/510^- C12 (**j**, 20 µM, 2h and **k**, 2 µM, 16h ; scale=20 µm).

**Figure S2.**
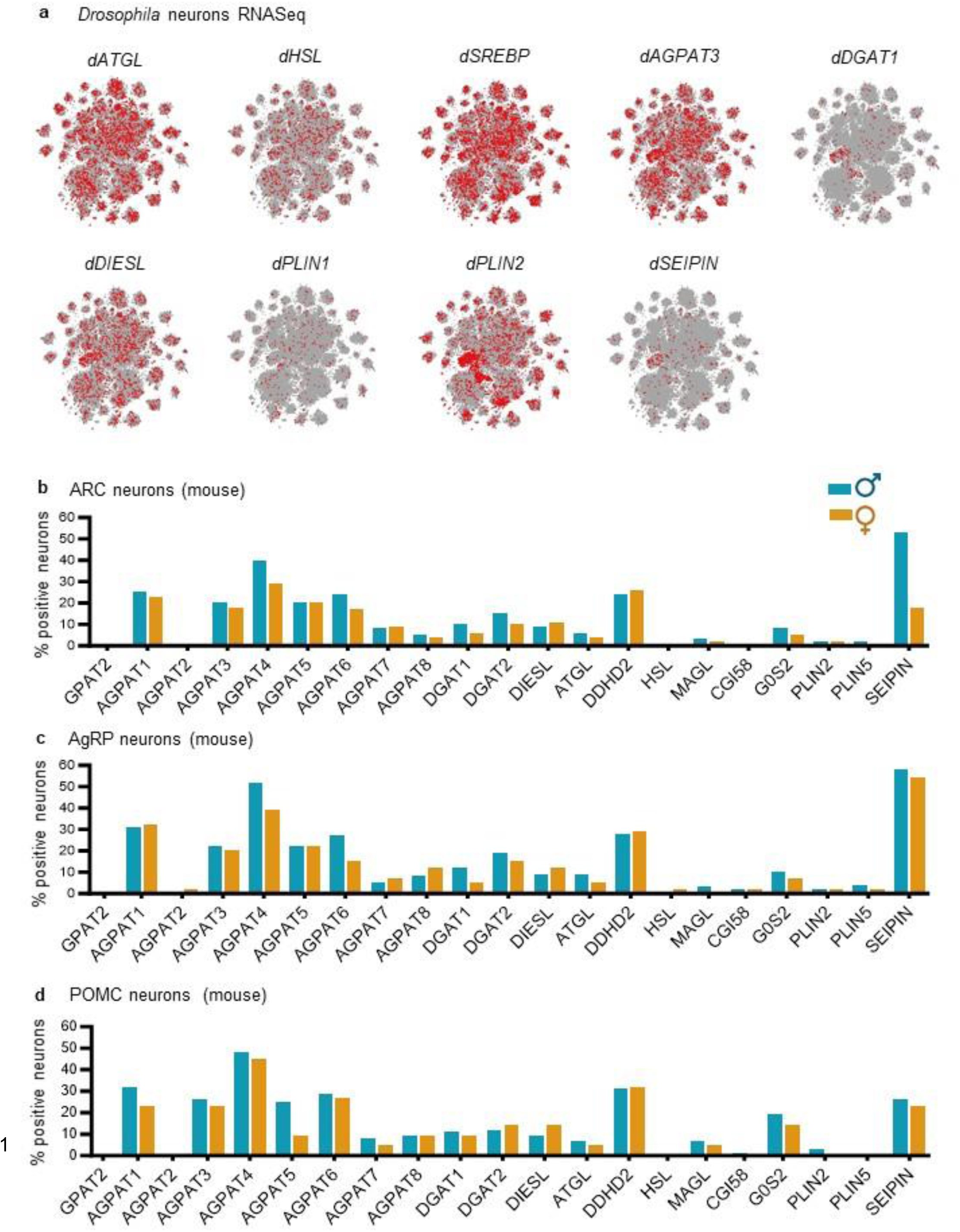
Lipid droplet-regulatory genes are expressed in *Drosophila* neurons and in mouse hypothalamic neurons. **a**, Seurat T-SNE of single-cell RNAseq data showing neurons in which mRNA from lipid droplet(LD)-regulatory genes were detected in *Drosophila*. Based on 10X cross-tissue data from neurons in Fly Cell Atlas^1^, visualized using SCope tool^2^. **b-d**, LD-regulatory gene expression profile analyzed from Single cell RNA sequencing (available dataset: GSE90806 in all ARC neurons (**b**) from 21 male (4445 cells) and 2 female mice (425 cells), (**c**) AgRP neurons (977 male cells, 41 female cells) and (**d**) POMC neurons (403 male cells, 22 female cells). See Table 1.

**Figure S3.**
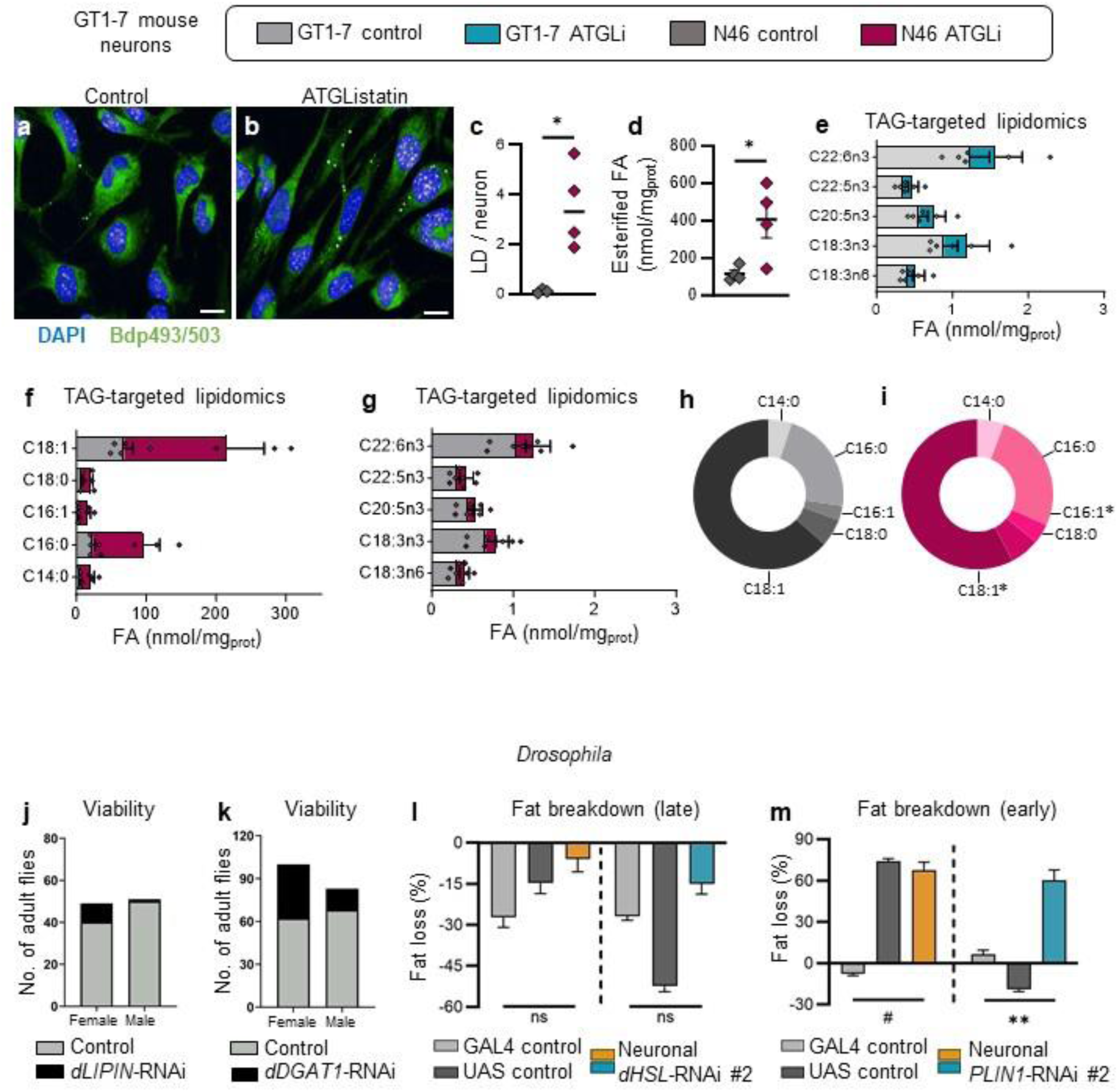
Lipid droplet-regulatory genes influence lipid droplet number in cultured hypothalamic neurons and energy homeostasis in flies. **a**, **b**, Representative images of Bodipy-stained lipid droplets (LD) in GT1-7 neurons treated with vehicle or ATGListatin (24h), scale=20 µm. **c**, Number of LD in N46 neurons treated with vehicle or ATGListatin (24h). N=3-4. **d**, Total amount of FA esterified into TG in N46 neurons treated with vehicle (DMSO) or ATGListatin (24h). **e**, Profile of polyunsaturated FA esterified into TG in GT1- 7 cells treated with vehicle (DMSO) or ATGListatin. C22:6n3, Docosahexaenoic acid; C22:5n3, Docosapentaenoic acid; C20:5n3, Eicosapentaenoic acid; C18:3n3, α-Linolenic acid; C18:3n6, ʎ- Linolenic acid. N= 3 independent experiments. **f,g**, Profile of FA esterified into TG in N46 cells treated with vehicle (DMSO) or ATGListatin. C14:0, Myristic acid; C16:0, Palmitic acid; C16:1, Palmitoleic acid; C18:0, Stearic acid; C18:1, Oleic acid. N= 3. **h,i**, Relative proportion of FA esterified into TG in N46 neurons treated with vehicle (DMSO) or ATGListatin (24h). N=3. Data are represented as mean ± SEM. Student’s t-test (**c,d**), multiple t-test (**h,i**) ; \**p<0.05*. Number of viable adults with pan-neuronal loss of *dLIPIN* (**j**) and *dDGAT1* (**k**). *Drosophila* fat breakdown between 12-24h (late) and 0-12h (early) post-fasting in females (orange) and males (turquoise) with pan-neuronal loss of *dHSL* (**l**) and *dPLIN1* (**m**) using independent RNAi lines. Fat breakdown data expressed as the mean body fat loss over a given period post-fasting +/- coefficient of error. Two-way ANOVA: ns indicates not significant, **p<0.01 RNAi genotype interaction, #*p*<0.05 control genotype interaction.

**Figure S4.**
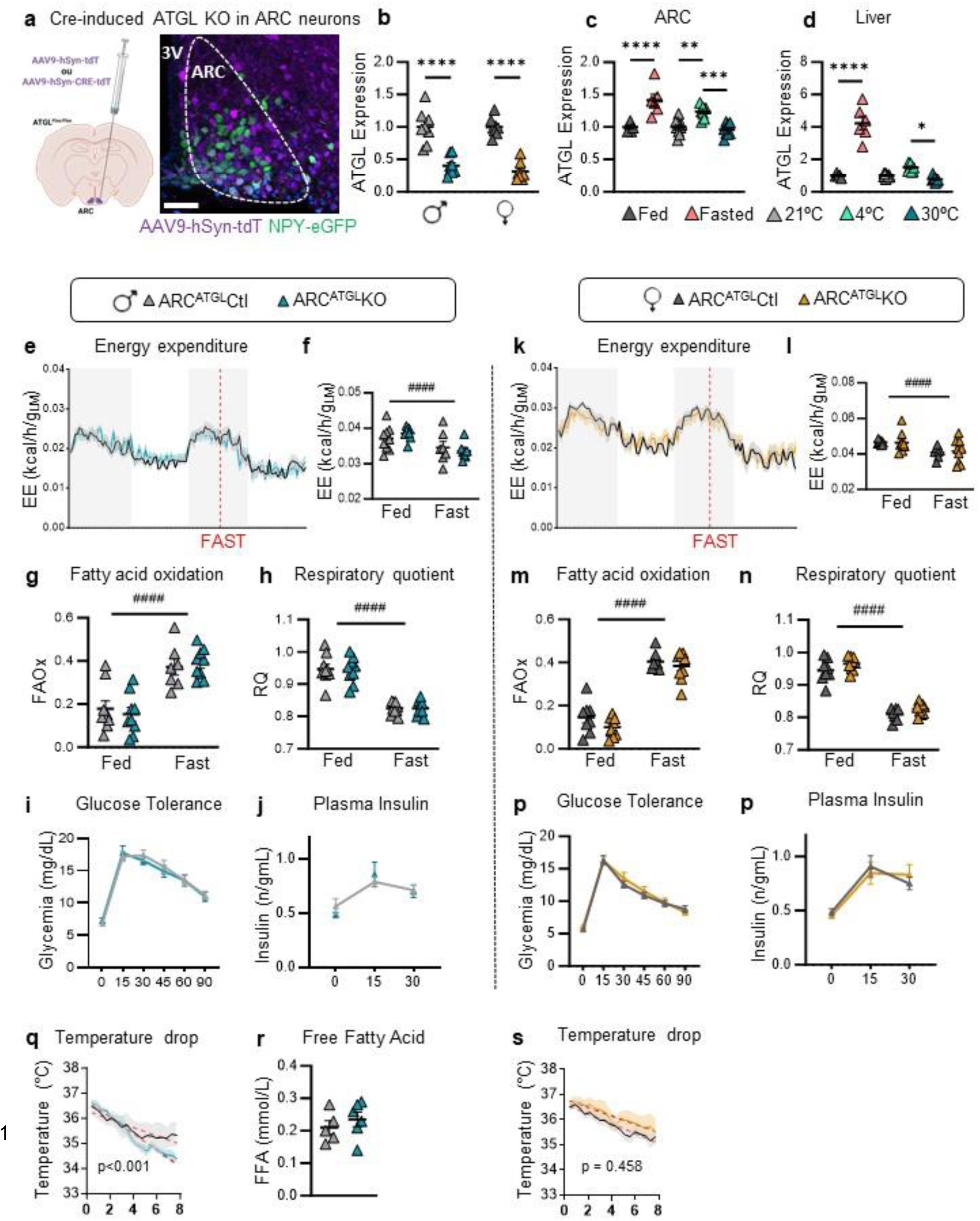
Loss of ATGL in ARC neurons does not impair metabolic responses to a fast or glucose homeostasis. **a**, Stereotaxic injection of AA9-hSyn-tdTomato or -CRE expressing viruses in the ARC nucleus of NPY-GFP reporter mice or ATGL floxed mice. NPY neurons (green) expressing the tdTomato protein 4 weeks post virus injection. Scale=50 µm. **b**, ATGL mRNA level (qPCR) in the ARC of male and female ATGL floxed mice injected with AA9-hSyn-tdTomato (ARC^ATGL^Ctl) or AA9-hSyn-CRE-tdTomato (ARC^ATGL^KO) viruses. N=8/group. **c**, ATGL mRNA level in the ARC and **d**, liver of male mice in fed (N=5) or fasted (16h) conditions (N=6) at 21°C, and in fed conditions after 24h at 21°C (N=11), 4 °C (N=8) or 30 °C (N=8). **e-n**, EE, FAOx and RQ in ARC^ATGL^CRE and ARC^ATGL^KO males and females in *ad libitum* fed (Fed) or fasted (16h) conditions (Fast). N=9 males and 8-9 females. **i, o**, Blood glucose and **j**, plasma insulin levels measured during glucose tolerance tests in male and **p**, female ARC^ATGL^CRE and ARC^ATGL^KO mice. N=10-11 males and 11-12 females. **q,s**, Drop in body temperature during the first 8h of cold exposure (4°C) in male and female ARC^ATGL^CRE and ARC^ATGL^KO mice. **r**, Plasma free fatty acid in male ARC^ATGL^CRE and ARC^ATGL^KO mice (N=5-6) after cold exposure. Data are represented as mean ± SEM. **b**, Student’s t-test; **c,d**, One-way ANOVA, Sidak post-hoc test; **e-n**, Two-way ANOVA, ####*p<*0.0001 time interaction, \**p<*0.05, \*\**p<*0.01, \*\*\**p<*0.001, \*\*\*\**p<*0.0001, genotype interaction, Sidak post-hoc test. **q-s**, Linear regression analysis, males *p<*0.0003.

**Figure S5.**
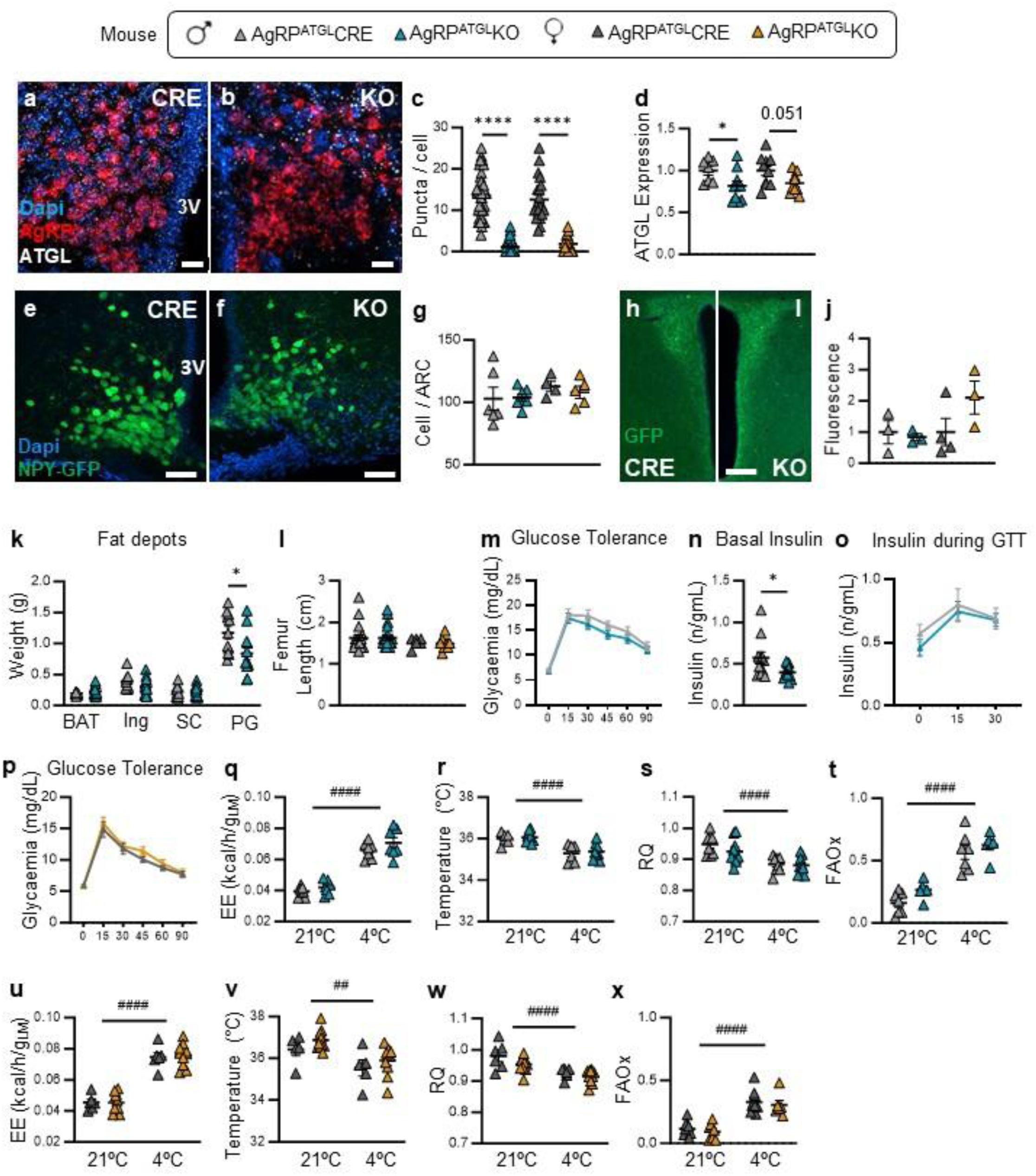
Validation and phenotyping of mice with AgRP-specific loss of ATGL. **a,b**, *In situ* detection of ATGL mRNA (white) by RNAScope in AgRP neurons (red) and **c**, ATGL puncta number in AgRP neurons of AgRP^ATGL^CRE and AgRP^ATGL^KO mice. N=20 cells/mouse, 2 Cre vs 2 KO males, 1 Cre vs 1 KO female, scale=20 µm. **d**, ATGL expression (qPCR) in ARC microdissections of male and female AgRP^ATGL^CRE and AgRP^ATGL^KO mice (N= 8-10 males and 10-10 females). **e,f**, NPY-positive neurons in the ARC of male and female AgRP^ATGL^CRE and AgRP^ATGL^KO mice (on NPY-GFP genetic background) and **g**, quantification. N= 6-7 males and 4- 5 females. Average count of a minimum of 2 ARC sections/mouse, scale=50 µm. **h,i**, GFP immunofluorescence in the PVN of males and females AgRP^ATGL^CRE and AgRP^ATGL^KO mice and **j**, its quantification. N=3-3 males and 3-4 females, scale=200 µm. **k**, Fat depot weight and **l**, femur length in males and females AgRP^ATGL^CRE and AgRP^ATGL^KO mice (N=19-21 males and 6-11 females). **m**, Blood glucose in males and **p**, females and **n,o**, plasma insulin levels during glucose tolerance tests in male AgRP^ATGL^CRE and AgRP^ATGL^KO mice (N=11-14 males and 6-11 females). Student’s t-test, \**p<*0.05. **q-t**, EE, body temperature, RQ and FAOx in male and **u-x**, female AgRP^ATGL^CRE vs AgRP^ATGL^KO mice during 24h at 21 °C or at 4 °C. N=7-7 males and 6-11 females. Data are represented as mean ± SEM. **a-n**, Student’s t-test, \**p<*0.05, \*\*\*\**p<*0.0001; Two-way ANOVA, ##*p<*0.01, ####*p<*0.0001, time interaction, Sidak post-hoc test.

**Figure S6.**
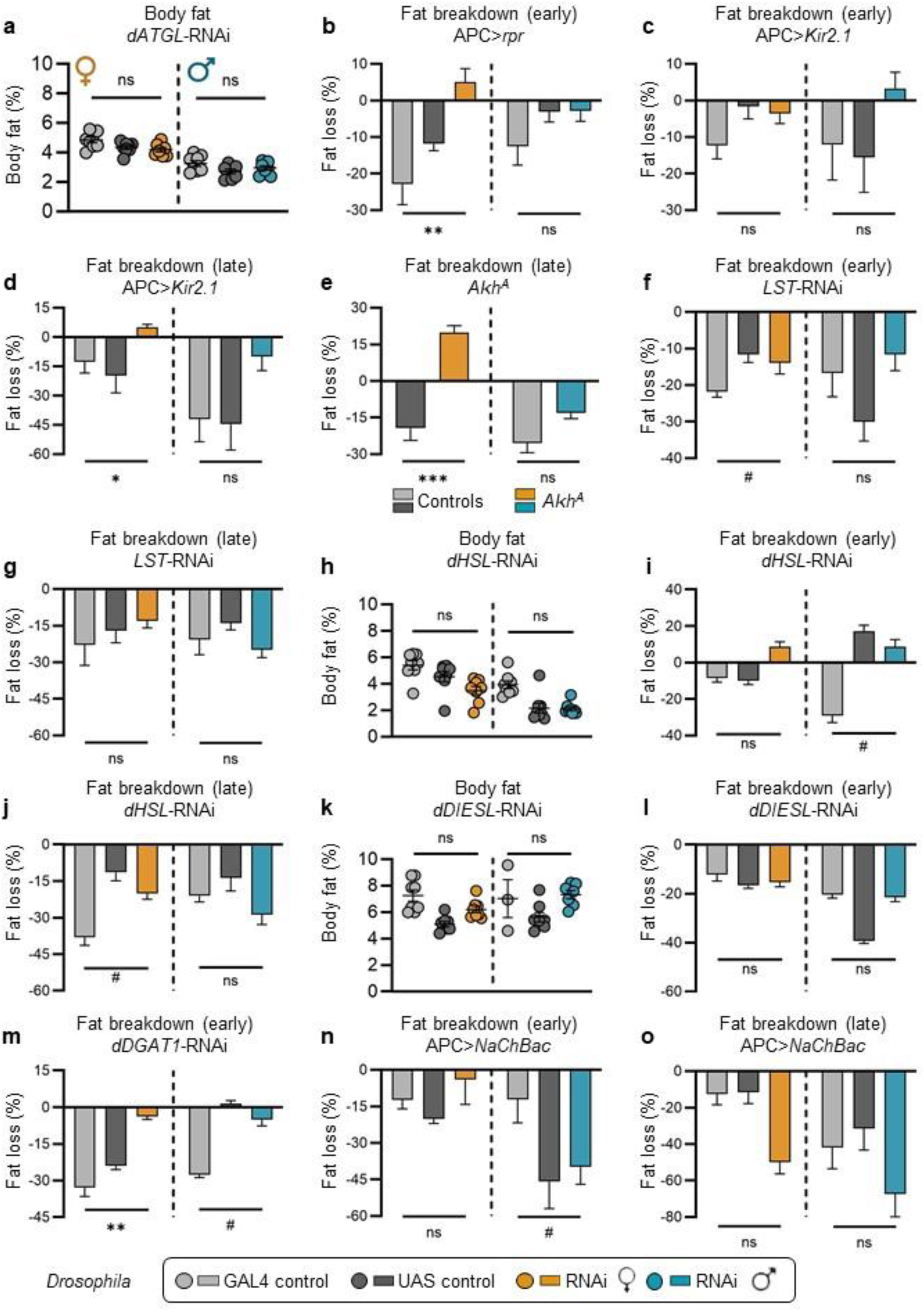
Genetic manipulation of adipokinetic hormone (Akh)-producing cells or Akh levels affects fat breakdown. **a**, Body fat in *Drosophila* females (orange) and males (turquoise) with adipokinetic hormone (Akh) cell (APC)-specific loss of *dATGL*. **b**, Fat breakdown 0-12h post-fasting in flies with APC-specific overexpression of proapoptotic gene *reaper* (*rpr*). **c**, Fat breakdown 0-12h and **d**, 12-24h post-fasting in flies with APC-specific overexpression of inwardly-rectifying potassium channel *Kir2.1*. **e**, Fat breakdown 12-24h post-fasting in *Akh* mutant flies. **f**, Fat breakdown 0-12h and **g**, 12-24h post-fasting in flies with APC-specific loss of *Limostatin* (*Lst*). **h**, Body fat and **i**, fat breakdown at 0-12h and **j**, 12-24h in flies with APC-specific loss of *dHSL*. **k**, Body fat and **l**, fat breakdown 0- 12h post-fasting in flies with APC-specific loss of *dDIESL*. **m**, Fat breakdown 0-12h post-fasting in flies with APC-specific loss of *dDGAT1*. **n**, Fat breakdown 0-12h and **o**, 12-24h post-fasting in flies with APC-specific overexpression of bacterial sodium channel *NaChBac*. Percent body fat expressed as mean +/- SEM. Fat breakdown data expressed as the mean percent body fat loss post-fasting +/- coefficient of error. Two-way ANOVA: ns indicates not significant, \**p<*0.05, \*\**p<*0.01, *** *p<*0.001, RNAi genotype interaction; #*p<*0.05 control genotype interaction.

**Figure S7.**
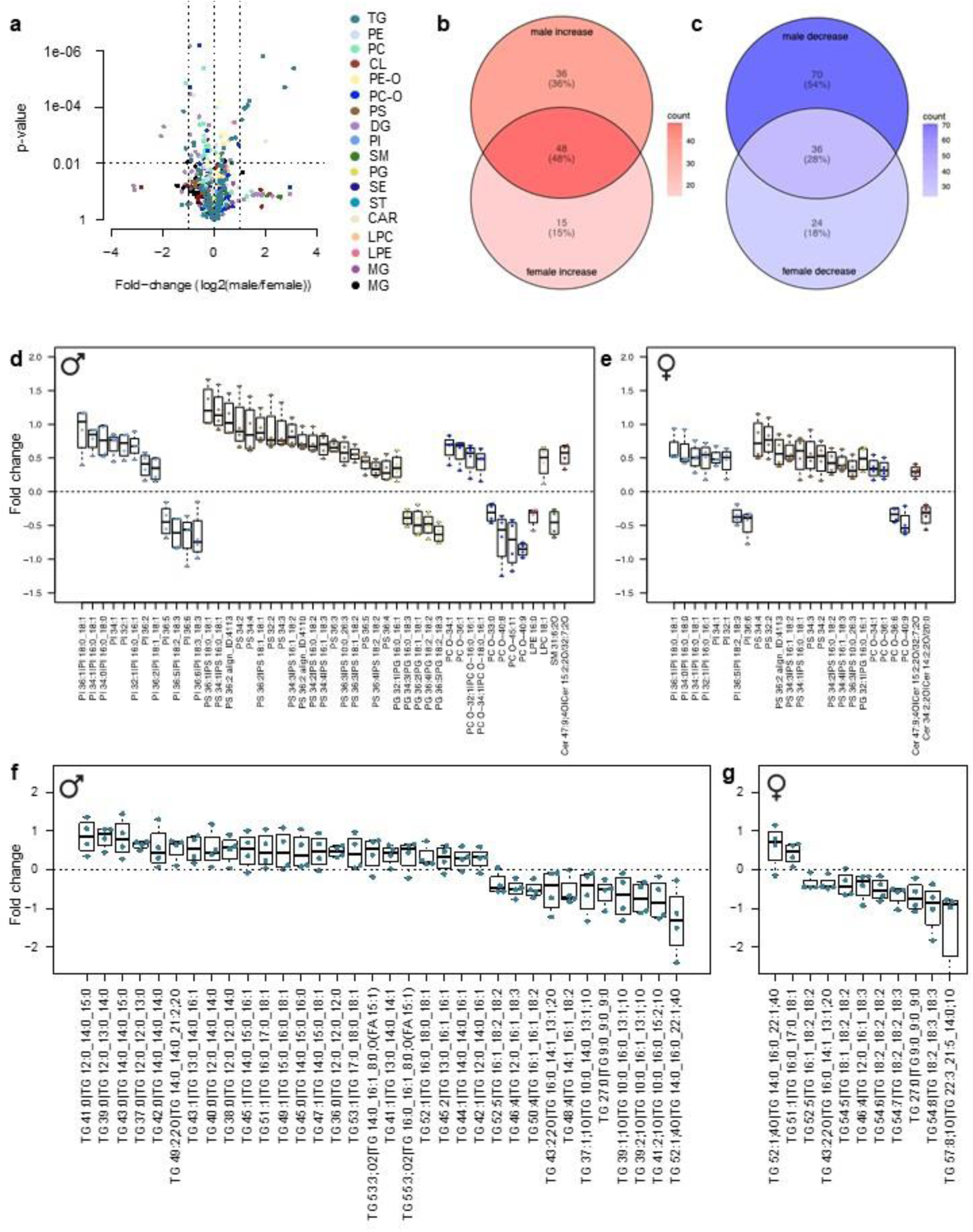
Unbiased lipidomic analysis of *Drosophila* male and female brains with and without neuronal dATGL. **a**, Volcano plot showing lipid classes that are differentially regulated between *Drosophila* male and female brains. Positive fold-change indicates male-biased lipids; negative fold-change indicates female-biased lipids. **b**, Venn diagram indicating lipid species that are upregulated in *Drosophila* female brains, male brains, or both upon neuronal loss of *dATGL*. **c**, Venn diagram showing shared- or uniquely-downregulated genes in response to neuronal loss of *dATGL*. **d**, Differentially-regulated lipid species in *Drosophila* male and **e**, female brains with neuronal loss of *dATGL*. Only lipids with fold-change >0.3 and unadjusted p<0.05 are shown. TG=Triacylglycerols; PE= Diacylglycerophosphoethanolamines; PC= Diacylglycerophosphocholines; CL= Cardiolipin; PEO= 1-alkyl,2- acylglycerophosphoethanolamines; PCO= 1-alkyl, 2-acylglycerophosphocholines; DG= Diacylglycerols; CAR= Fatty acyl carnitines; MG= monoacylglycerols; CE= Steryl esters; ST= sterols; PI= Diacylglycerophosphoinositols; PS= Diacylglycerophosphoserines; PG= phosphatidylglycerol; LPE= Monoacylglycerophosphoethanolamines; LPC= Monoacylglycerophosphocholines; SM= Ceramide phosphocholines (sphingomyelins); Cer= Ceramide. **f**, TG levels in *Drosophila* male and **g**, female brains with neuronal loss of *dATGL*; all TG species detected are shown, only 3 TG species were differentially regulated in female brains compared with 12 differentially regulated TG species in males (see Supplemental table 1).

**Figure S8.**
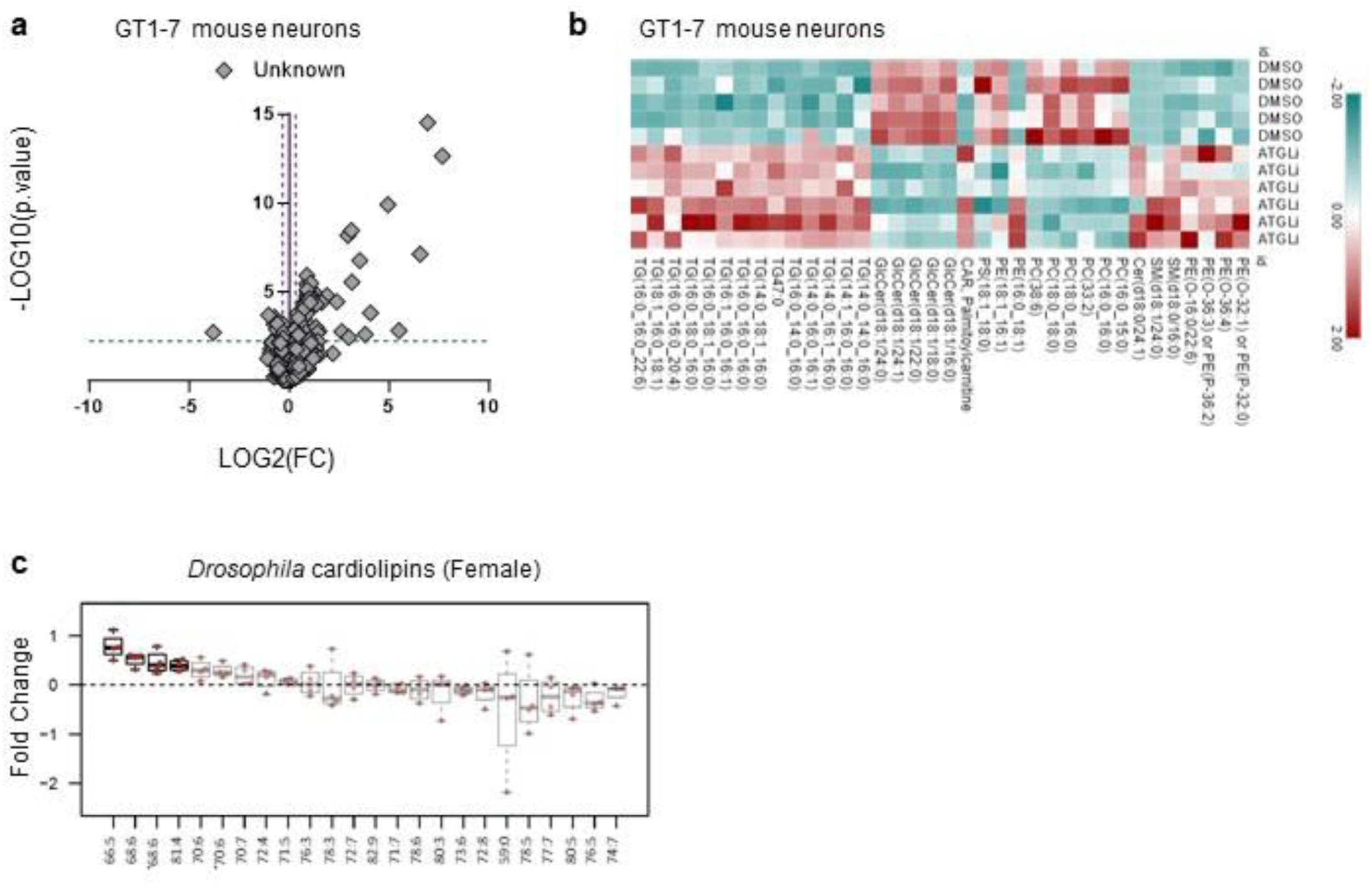
Lipid remodeling in cultured hypothalamic neurons in response to ATGL inhibition. **a**, Volcano plot of 1073 unidentified lipid features (untargeted lipidomics) in GT1-7 neurons treated with vehicle or ATGListatin during 24h. The x-axis represents the fold changes of MS signal intensities for all the features in the treated group compared with the untreated one and expressed as log2. The y axis corresponds to the p values expressed as -log10. N=6. **b**, Heatmap of annotated unique lipids (untargeted lipidomics) in GT-7 neurons treated with vehicle or ATGListatin for 24h. Heatmap was generated using the online Morpheus software (https://software.broadinstitute.org/morpheus). N=5-6. **c**, Differentially regulated cardiolipin species in *Drosophila* adult female brains with neuronal loss of *dATGL*.

**Figure S9.**
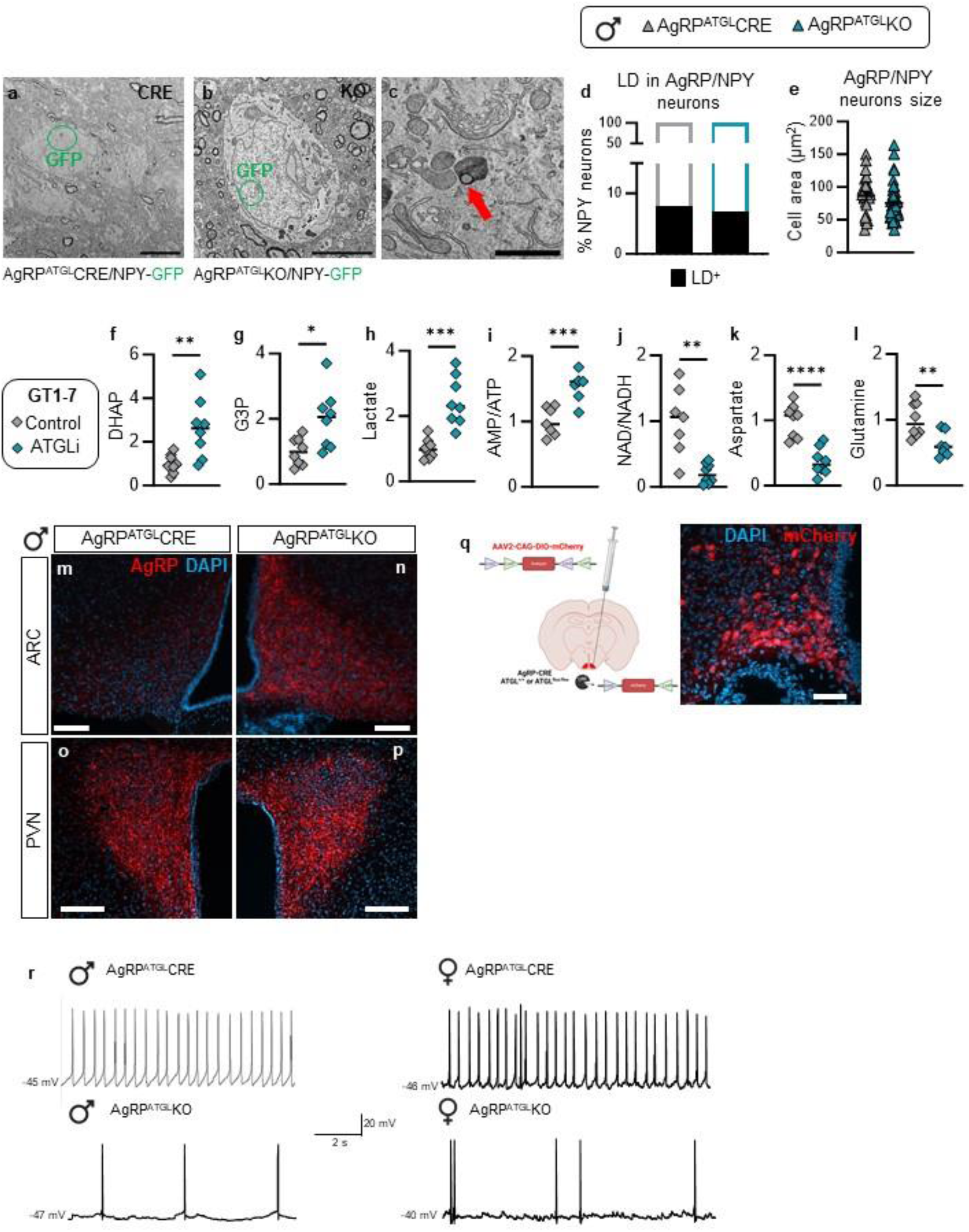
Loss or inhibition of ATGL affects metabolism and activity of hunger-activated neurons. **a-b**, EM identification of AgRP neurons using GFP immunostaining in AgRP^ATGL^CRE and AgRP^ATGL^KO mice on the NPY-GFP genetic background. Scale=5 µm. **c**, Representative EM image of an LD (red arrow) in AgRP neurons. Scale=1 µm. **d**, Percentage of LD-positive AgRP neurons and **e**, AgRP neuron area in male AgRP^ATGL^CRE and AgRP^ATGL^KO mice. N=26 CRE and 53 KO cells KO from 2 CRE and 2 KO mice. **f-l**, Relative metabolite and amino acid levels or ratios in GT1-7 neurons treated with DMSO or ATGListatin during 24h. N=7-8. Student’s t.test, \**p<*0.05, \*\**p<*0.01, *** *p<*0.001. **m-p**, Representative images of AgRP immunostaining in **m-n**, ARC and **o-p**, PVN. Scale=100 µm. **q**, Stereotaxic injections of Cre-dependent mCherry viruses (AAV2-CAG-DIO-mCherry) in the ARC of male and female AgRP^ATGL^CRE vs AgRP^ATGL^KO mice for electrophysiological recordings. mCherry fluorescence in AgRP. Scale=50 µm. **r**, Traces of action potentials in males and females AgRP^ATGL^CRE and AgRP^ATGL^KO.

