## Supplemental Table 1 for "Neuronal lipid droplets play a conserved and sex-biased role in maintaining whole-body energy homeostasis"

| **Differentially Regulated Lipids in Female brains with Neuronal loss of *dATGL*** | | |
|---|---|---|
| **Lipid identity** | **log2FC (RNAi/controls)** | **p-value** |
| PC 35:1\|PC 17:0_18:1 | 0.320048 | 7.05E-06 |
| PE O-34:2\|PE O-18:1_16:1 | 0.327179 | 4.16E-05 |
| PC 36:2\|PC 18:1_18:1 | -0.05164 | 0.000155 |
| PC 36:5\|PC 18:2_18:3 | -0.21404 | 0.000544 |
| PC 38:2 align_ID:4336 | -0.3051 | 0.000585 |
| PE 34:3 | 0.15448 | 0.000933 |
| CAR 20:0 | -0.36111 | 0.000971 |
| Cer 48:11;4O\|Cer 13:1;2O/35:10;2O | -0.12972 | 0.001147 |
| PI 34:1 | 0.509527 | 0.001288 |
| PE 36:2;O\|PE 18:2_18:0;O | -1.62206 | 0.00137 |
| CL 68:6\|CL 16:1_18:2_16:1_18:2 | 0.520995 | 0.001389 |
| PC 36:4 | -0.18219 | 0.001752 |
| PE 33:2 | 0.318708 | 0.001914 |
| PE 36:5\|PE 18:2_18:3 | -0.27341 | 0.001937 |
| DG 41:5 | 0.199905 | 0.002019 |
| PC 38:1 align_ID:4355 | 0.270406 | 0.002129 |
| Cer 34:1;2O\|Cer 14:1;2O/20:0 | -0.17623 | 0.002278 |
| PC O-36:4 | -0.2563 | 0.002428 |
| PC 38:1\|PC 20:0_18:1 | 0.180642 | 0.002438 |
| CAR 18:2 | -0.48043 | 0.002441 |
| PC 36:5 | -0.42263 | 0.002522 |
| PC 36:1\|PC 18:0_18:1 | 0.141552 | 0.002666 |
| CAR 22:0 | -0.33301 | 0.002992 |
| PC 33:1\|PC 16:0_17:1 | 0.188551 | 0.003353 |
| TG 50:4\|TG 16:1_16:1_18:2 | -0.19977 | 0.003878 |
| Cer 46:9;4O\|Cer 15:2;2O/31:7;2O | 0.181106 | 0.004239 |
| PS 34:3\|PS 16:1_18:2 | 0.591902 | 0.004381 |
| PE 32:2 | 0.438595 | 0.004434 |
| ST 28:3;O align_ID:1222 | 0.176061 | 0.004536 |
| PC O-34:2\|PC O-16:0_18:2 | -0.20224 | 0.004719 |
| PC 32:0 | 0.370502 | 0.004969 |
| PC O-36:6 | -0.34031 | 0.005091 |
| PC 30:0\|PC 14:0_16:0 | -0.19099 | 0.005161 |
| PC 36:6\|PC 18:3_18:3 | -0.35014 | 0.005214 |
| PS 34:4\|PS 16:1_18:3 | 0.432167 | 0.005943 |
| Cer 47:9;4O\|Cer 15:2;2O/32:7;2O | 0.300397 | 0.006255 |
| TG 54:7\|TG 18:2_18:2_18:3 | -0.6419 | 0.006714 |
| HexCer 34:2;2O | -0.32743 | 0.006843 |
| SE 29:1/18:2 | -0.37551 | 0.006896 |
| Cer 34:2;2O\|Cer 14:2;2O/20:0 | -0.33895 | 0.007497 |
| PE 36:4;2O\|PE 18:2_18:2;2O | -0.45466 | 0.00763 |

| | | |
|---|---|---|
| PI 36:1\|PI 18:0_18:1 | 0.641467 | 0.007822 |
| PS 32:2 | 0.794611 | 0.008135 |
| PE O-32:1\|PE O-16:0_16:1 | 0.361761 | 0.00832 |
| PC 32:0\|PC 16:0_16:0 | -0.18186 | 0.008365 |
| PI 36:5\|PI 18:2_18:3 | -0.35512 | 0.008529 |
| PG 32:1\|PG 16:0_16:1 | 0.488922 | 0.00909 |
| PE O-34:1\|PE O-18:0_16:1 | 0.165506 | 0.009831 |
| PE 36:6\|PE 18:3_18:3 | -0.39666 | 0.010097 |
| CL 66:5\|CL 16:1_16:1_16:1_18:2 | 0.799818 | 0.011333 |
| PI 34:1\|PI 16:0_18:1 | 0.521545 | 0.011442 |
| PC 34:0 | 0.250798 | 0.012387 |
| FA 30:2 | -0.49493 | 0.012401 |
| PI 32:1 | 0.468575 | 0.013195 |
| CAR 18:1 | -0.22608 | 0.013255 |
| PC O-36:1 | 0.336188 | 0.013341 |
| TG 54:8\|TG 18:2_18:3_18:3 | -0.89248 | 0.013365 |
| Cer 49:10;4O\|Cer 14:2;2O/35:8;2O | -0.11905 | 0.013576 |
| PE O-40:3\|PE O-22:0_18:3 | -0.27484 | 0.014067 |
| PC 30:1\|PC 14:0_16:1 | 0.212705 | 0.014932 |
| PE O-32:1 | 0.254892 | 0.015117 |
| LPE-N (FA)33:2\|LPE-N (FA 15:0)18:2 | 0.223518 | 0.015422 |
| PE 36:1;O\|PE 18:0_18:1;O | -1.62637 | 0.016067 |
| PC 36:4\|PC 18:2_18:2 | -0.06439 | 0.016186 |
| PC 34:4\|PC 18:1_16:3 | -0.14377 | 0.016616 |
| PC 37:8 | -0.28384 | 0.016869 |
| PE 33:2;2O\|PE 16:1_17:1;2O | 0.68526 | 0.017024 |
| TG 48:4\|TG 14:1_16:1_18:2 | -0.29646 | 0.017055 |
| PC O-32:1\|PC O-16:0_16:1 | 0.260211 | 0.017232 |
| Cer 34:0;2O\|Cer 14:0;2O/20:0 | -0.16743 | 0.017286 |
| PI 34:0\|PI 16:0_18:0 | 0.594327 | 0.018209 |
| DG 48:11 | -0.21506 | 0.018394 |
| PI 32:1\|PI 16:0_16:1 | 0.511758 | 0.019367 |
| PC O-34:1 | 0.355102 | 0.019924 |
| PE-Cer 36:0;3O | 0.273048 | 0.019941 |
| PC 34:0\|PC 16:0_18:0 | -0.16347 | 0.019979 |
| PE 33:1\|PE 16:0_17:1 | 0.25584 | 0.019988 |
| FA 28:2 | -0.63917 | 0.020085 |
| PE O-38:3\|PE O-20:0_18:3 | -0.15348 | 0.020595 |
| PC O-36:2\|PC O-18:0_18:2 | -0.18149 | 0.020835 |
| PS 36:2 align_ID:4113 | 0.601385 | 0.020898 |
| MG 36:3 | -0.32386 | 0.022064 |
| PE 35:4\|PE 17:2_18:2 | -0.35907 | 0.022194 |
| CL 81:4\|CL 17:0_28:0_18:2_18:2 | 0.39641 | 0.022275 |
| PC 33:2 | 0.18279 | 0.022686 |
| PE O-37:3\|PE O-19:0_18:3 | -0.54631 | 0.022728 |
| PS 34:4 | 0.794637 | 0.022941 |

| | | |
|---|---|---|
| PG 32:0\|PG 16:0_16:0 | 0.290265 | 0.023254 |
| PC 33:0 | 0.157076 | 0.025956 |
| PI 36:6 | -0.4597 | 0.026051 |
| PS 34:2\|PS 16:0_18:2 | 0.44537 | 0.026077 |
| PE O-36:7\|PE O-18:4_18:3 | -0.2786 | 0.026237 |
| DG 26:0\|DG 12:0_14:0 | 0.271665 | 0.02684 |
| PC 36:3\|PC 18:1_18:2 | 0.038332 | 0.026877 |
| PC 33:1 | 0.208117 | 0.027098 |
| PE O-36:4\|PE O-18:1_18:3 | -0.30798 | 0.028361 |
| FA 16:0 align_ID:295 | -0.29121 | 0.028755 |
| CL 70:6\|CL 16:1_18:1_18:2_18:2 | 0.292181 | 0.029328 |
| TG 54:6\|TG 18:2_18:2_18:2 | -0.48456 | 0.029482 |
| PS 34:2 | 0.509602 | 0.029943 |
| PE-Cer 37:3;3O | -0.26469 | 0.030642 |
| DG 36:5\|DG 18:2_18:3 | -0.40247 | 0.031531 |
| PC O-37:4 | 0.163662 | 0.032165 |
| PE P-34:1\|PE P-18:0_16:1 | 0.238928 | 0.03291 |
| PC 26:0 align_ID:2773 | -0.5817 | 0.03515 |
| PE-Cer 34:0;3O | 0.229763 | 0.03521 |
| PE 34:3\|PE 16:1_18:2 | 0.161299 | 0.037449 |
| PE O-36:5 | -0.25328 | 0.037508 |
| CAR 18:0 | -0.31369 | 0.037863 |
| DG 27:0 | -0.54542 | 0.03834 |
| CL 68:6\|CL 16:1_16:1_18:2_18:2 | 0.471797 | 0.038494 |
| PE 35:1;2O\|PE 18:0_17:1;2O | 0.602603 | 0.0385 |
| PS 36:5 | 0.296119 | 0.038522 |
| PC O-40:9 | -0.47153 | 0.038851 |
| FA 39:0 | -0.22205 | 0.039374 |
| Cer 48:9;4O\|Cer 17:3;2O/31:6;2O | 0.157951 | 0.042211 |
| PS 34:3 | 0.563063 | 0.04319 |
| PS 34:1\|PS 16:0_18:1 | 0.563483 | 0.043773 |
| FAHFA 26:0;O\|FAHFA 16:0/10:0;O | -0.69404 | 0.043926 |
| FAHFA 30:0;O\|FAHFA 16:0/14:0;O | -0.97809 | 0.044045 |
| PC 32:1\|PC 16:0_16:1 | 0.042298 | 0.044213 |
| DG 44:7 | 0.139461 | 0.044542 |
| Cer 36:2;2O\|Cer 14:2;2O/22:0 | -0.17368 | 0.044825 |
| PC O-36:2 | -0.09931 | 0.047865 |
| PG 34:3\|PG 16:1_18:2 | 0.28432 | 0.048684 |
| DG 39:5 | 0.150404 | 0.048805 |
| PS 36:3\|PS 10:0_26:3 | 0.347222 | 0.049092 |
| PS 35:2;O\|PS 18:1_17:1;O | -0.29143 | 0.049208 |
| PE 36:1\|PE 18:0_18:1 | 0.070437 | 0.050564 |
| PC O-34:1\|PC O-18:0_16:1 | 0.196305 | 0.050627 |
| PS 36:1\|PS 18:0_18:1 | 0.643225 | 0.051205 |
| PI 36:3 | 0.194753 | 0.051934 |
| PE 32:1\|PE 16:0_16:1 | 0.23091 | 0.052681 |

| | | |
|---|---|---|
| PE O-35:3\|PE O-17:0_18:3 | -0.21991 | 0.054943 |
| PC O-36:5 | -0.25445 | 0.056024 |
| TG 51:1\|TG 16:0_17:0_18:1 | 0.443624 | 0.056152 |
| PI 36:2 | 0.285329 | 0.05723 |
| PE 32:2\|PE 16:1_16:1 | 0.346314 | 0.057902 |
| CL 70:6\|CL 16:0_18:2_18:2_18:2 | 0.319682 | 0.057962 |
| PC O-34:8 | 1.967896 | 0.060189 |
| TG 46:4\|TG 12:0_16:1_18:3 | -0.39494 | 0.061462 |
| PE-Cer 36:5;2O | -0.20434 | 0.061921 |
| PI 34:2 | 0.233638 | 0.0654 |
| PS 36:3 | 0.344436 | 0.065657 |
| PC O-35:1 | 0.219101 | 0.066303 |
| PE O-32:0\|PE O-18:0_14:0 | -0.18843 | 0.06891 |
| PC O-33:2 align_ID:3465 | -0.56072 | 0.069358 |
| PS 36:2 align_ID:4110 | 0.451311 | 0.071174 |
| PC 40:2 | -0.16409 | 0.071343 |
| MG 21:1 | -0.45023 | 0.072 |
| PC O-32:3 | -0.18561 | 0.072269 |
| PE O-34:2 | 0.18479 | 0.072289 |
| PI 36:6\|PI 18:3_18:3 | -0.47115 | 0.072782 |
| PS 35:1\|PS 10:0_25:1 | -0.22154 | 0.072841 |
| PI 34:3 | 0.33031 | 0.074223 |
| PE O-36:1\|PE O-18:0_18:1 | 0.093803 | 0.075207 |
| DG O-42:4\|DG O-22:3_20:1 | -0.68515 | 0.075937 |
| PS 36:4\|PS 18:2_18:2 | 0.213132 | 0.076263 |
| PE 36:3;2O\|PE 18:1_18:2;2O | -0.18566 | 0.076721 |
| PG 32:2\|PG 16:1_16:1 | 0.361923 | 0.07691 |
| CL 76:5\|CL 24:0_16:1_18:1_18:3 | -0.2956 | 0.077406 |
| TG 52:4\|TG 16:1_18:1_18:2 | -0.2748 | 0.077893 |
| PC O-35:3 align_ID:3728 | -1.70862 | 0.078227 |
| PS 36:3\|PS 18:1_18:2 | 0.329916 | 0.07915 |
| PC 36:0 | -0.15602 | 0.080127 |
| PC 36:2 align_ID:4298 | -0.1225 | 0.080846 |
| TG 52:5\|TG 16:1_18:2_18:2 | -0.35197 | 0.081089 |
| PE O-38:3 | -0.09524 | 0.083862 |
| PE 35:1\|PE 17:0_18:1 | 0.144844 | 0.08404 |
| DG 52:10 | -0.18475 | 0.085282 |
| SM 61:6;2O | -0.1469 | 0.086117 |
| PC 36:0\|PC 18:0_18:0 | -0.1832 | 0.08669 |
| PE 38:3\|PE 20:0_18:3 | -0.14289 | 0.08674 |
| PC O-38:5 | -0.15258 | 0.086842 |
| Cer 41:1;4O | -0.34476 | 0.088142 |
| PI 36:3\|PI 18:1_18:2 | 0.214695 | 0.088669 |
| PE O-36:3\|PE O-18:0_18:3 | -0.09998 | 0.090006 |
| PI 32:2 | 0.272265 | 0.090477 |
| PC 38:2\|PC 20:0_18:2 | -0.25728 | 0.092811 |

| | | |
|---|---|---|
| PE O-38:1\|PE O-20:0_18:1 | 0.155665 | 0.093261 |
| DG 50:7 | -0.18375 | 0.094629 |
| SE 29:1/16:2 | -0.25724 | 0.095109 |
| TG 54:4\|TG 18:1_18:1_18:2 | -0.26874 | 0.095602 |
| PE 34:1 | 0.147465 | 0.098041 |
| PE 34:1\|PE 16:0_18:1 | 0.083002 | 0.098639 |
| PC O-37:5 | -0.10489 | 0.101875 |
| PS 36:2\|PS 18:1_18:1 | 0.437476 | 0.103368 |
| PI 36:2\|PI 18:1_18:1 | 0.256617 | 0.105763 |
| Cer 38:0;2O\|Cer 14:0;2O/24:0 | -0.16177 | 0.106572 |
| TG 53:1\|TG 17:0_18:0_18:1 | 0.292913 | 0.107174 |
| PI 36:5 | -0.20237 | 0.109529 |
| PC 32:3 | -0.12707 | 0.109627 |
| PG 34:1\|PG 16:0_18:1 | 0.292987 | 0.109796 |
| PC 40:6 | -0.21644 | 0.110117 |
| TG 52:1;4O\|TG 14:0_16:0_22:1;4O | 0.683621 | 0.110983 |
| PE O-36:6\|PE O-18:4_18:2 | -0.18327 | 0.111782 |
| PS 36:4 | 0.246317 | 0.113331 |
| TG 54:5\|TG 18:1_18:2_18:2 | -0.35295 | 0.120301 |
| CL 73:6 | -0.10853 | 0.120956 |
| PG 36:2\|PG 18:1_18:1 | -0.16573 | 0.121803 |
| Cer 36:1;2O\|Cer 14:1;2O/22:0 | -0.09971 | 0.12219 |
| PE P-36:5\|PE P-18:3_18:2 | 0.145956 | 0.123434 |
| NAE 22:4 align_ID:1089 | -0.69407 | 0.130113 |
| PE 36:2;2O\|PE 18:0_18:2;2O | -0.19706 | 0.130499 |
| PG 36:5\|PG 18:2_18:3 | -0.25172 | 0.130816 |
| PE O-34:0\|PE O-18:0_16:0 | -0.15229 | 0.131838 |
| FA 20:0 | -0.52266 | 0.131856 |
| FA 18:3 align_ID:341 | -0.33275 | 0.131929 |
| PS 36:6 | 0.192846 | 0.133512 |
| FA 28:1 | -0.3395 | 0.134734 |
| SE 28:1/18:2 | -0.62469 | 0.135072 |
| PC O-37:8 | -0.1817 | 0.13551 |
| Cer 40:1;2O\|Cer 16:1;2O/24:0 | -0.26966 | 0.137129 |
| TG 43:2;2O\|TG 16:0_14:1_13:1;2O | -0.35774 | 0.139829 |
| NAE 20:0 | -0.34438 | 0.141096 |
| PC 35:4 | -0.21411 | 0.145406 |
| TG O-37:0\|TG O-13:0_12:0_12:0 | -0.35783 | 0.146173 |
| Cer 49:11;4O\|Cer 13:2;2O/36:9;2O | -0.16939 | 0.148701 |
| SM 39:4;3O | 0.221343 | 0.152479 |
| NAE 22:5 | -0.80285 | 0.153766 |
| PE 30:1 | -0.17272 | 0.155354 |
| FA 42:5 | -0.51236 | 0.156092 |
| ST 28:4;O align_ID:1213 | -0.21543 | 0.157205 |
| CL 70:7\|CL 16:1_18:2_18:2_18:2 | 0.202065 | 0.158017 |
| PC 38:6 | 0.359423 | 0.15865 |

| | | |
|---|---|---|
| PC 28:1 | 0.28326 | 0.158744 |
| PE-Cer 32:1;2O\|PE-Cer 14:1;2O/18:0 | -0.17049 | 0.160331 |
| Cer 28:0;4O | -0.20309 | 0.162471 |
| TG 57:8;1O\|TG 22:3_21:5_14:0;1O | -1.19876 | 0.162737 |
| SM 39:6;3O | -0.59899 | 0.163872 |
| DG 35:1\|DG 17:0_18:1 | 0.312249 | 0.164139 |
| NAE 20:2 | -0.79983 | 0.16778 |
| DG 34:3\|DG 16:1_18:2 | -0.11921 | 0.168935 |
| NAE 20:1 | -0.70527 | 0.172728 |
| FA 24:0 align_ID:559 | -0.24759 | 0.174249 |
| PC 37:2 | -0.14959 | 0.178062 |
| DG O-33:1\|DG O-17:0_16:1 | -1.24829 | 0.178949 |
| PE 37:3\|PE 19:1_18:2 | -0.14071 | 0.18099 |
| DG O-42:3\|DG O-15:3_27:0 | -0.24812 | 0.182361 |
| TG 47:5;1O\|TG 14:1_18:2_15:2;1O | 0.127628 | 0.182616 |
| PC O-33:2 align_ID:3472 | 0.125306 | 0.182876 |
| DG 42:9 | -0.76387 | 0.185525 |
| DG 32:2 | -0.97333 | 0.186908 |
| PA 36:3\|PA 18:1_18:2 | 0.203237 | 0.187693 |
| SM 44:1;3O | -0.09694 | 0.18829 |
| DG 28:2\|DG 14:1_14:1 | -0.20234 | 0.188318 |
| MG 18:2 | -0.21701 | 0.188409 |
| MG 26:6 | -0.57674 | 0.189282 |
| CAR 16:0 | -0.22629 | 0.190564 |
| Cer 36:0;2O\|Cer 14:0;2O/22:0 | -0.1381 | 0.190726 |
| PE 25:1;O\|PE 10:0_15:1;O | -0.25898 | 0.191312 |
| FA 22:0 | -0.24452 | 0.191573 |
| TG 27:0\|TG 9:0_9:0_9:0 | -0.65909 | 0.192504 |
| CL 71:5 | 0.066338 | 0.192578 |
| MG 16:0 | -0.32822 | 0.19309 |
| PE 36:4\|PE 18:2_18:2 | -0.06223 | 0.19372 |
| PI 34:2\|PI 16:1_18:1 | 0.182063 | 0.194044 |
| FA 30:0 | -0.21895 | 0.195957 |
| PC 35:4;O\|PC 18:2_17:2;O | 0.489628 | 0.196032 |
| FA 9:0 align_ID:135 | 0.250795 | 0.197679 |
| PC 34:2;O\|PC 18:1_16:1;O | -0.27487 | 0.200922 |
| DG 21:0 | -0.24599 | 0.203109 |
| TG 39:1;1O\|TG 10:0_16:0_13:1;1O | 0.328534 | 0.203289 |
| TG 46:3\|TG 14:1_16:1_16:1 | -0.17872 | 0.203668 |
| DG 44:9 | 0.153934 | 0.203876 |
| SM 43:8;2O | -0.87013 | 0.204168 |
| TG 39:0\|TG 12:0_13:0_14:0 | 0.4162 | 0.204242 |
| NAE 18:4 | -0.33447 | 0.206052 |
| PC 40:8 | -0.21488 | 0.206874 |
| PE 27:1;O\|PE 10:0_17:1;O align_ID:1107 | -0.20305 | 0.20769 |
| PC 37:6 | -0.18357 | 0.209219 |

| | | |
|---|---|---|
| PC 35:3\|PC 17:1_18:2 | -0.11841 | 0.209913 |
| PE 27:1;O\|PE 10:0_17:1;O align_ID:1108 | -0.16995 | 0.211267 |
| TG 52:6\|TG 16:1_18:2_18:3 | 0.363174 | 0.212553 |
| PE O-34:0 | -0.10437 | 0.212841 |
| PC 33:3 | 0.097052 | 0.214193 |
| PE O-37:2 | 0.148525 | 0.214645 |
| SM 43:1;3O | -0.16067 | 0.21467 |
| Cer 32:1;2O\|Cer 14:1;2O/18:0 | -0.11199 | 0.215608 |
| Cer 34:1;3O\|Cer 18:1;2O/16:0;(2OH) | -0.23982 | 0.215741 |
| TG 53:2\|TG 17:0_18:1_18:1 | 0.175302 | 0.215964 |
| PA 36:4\|PA 18:2_18:2 | 0.220194 | 0.219386 |
| Cer 44:1;3O\|Cer 18:1;2O/26:0;O | -0.30575 | 0.219854 |
| Cer 42:1;3O\|Cer 18:1;2O/24:0;(2OH) | -0.2808 | 0.22146 |
| Diisodecyl phthalate | -0.38006 | 0.221662 |
| MG 38:1 | -0.29446 | 0.223649 |
| FA 32:0 | -0.31893 | 0.228018 |
| CL 74:7 | -0.59805 | 0.229757 |
| TG 39:2;2O\|TG 14:0_15:2_10:0;2O | -0.30343 | 0.232278 |
| PE-Cer 34:1;3O | -0.16758 | 0.235608 |
| PE 37:1 | 0.147391 | 0.235711 |
| SM 39:2;3O | 0.086126 | 0.236072 |
| ST 28:4;O align_ID:1101 | -0.19273 | 0.236366 |
| FA 27:0 align_ID:640 | -0.24111 | 0.237646 |
| PE P-36:6\|PE P-18:3_18:3 | -0.0993 | 0.23793 |
| PC O-41:3 | 0.347946 | 0.238249 |
| PE 38:2\|PE 20:0_18:2 | -0.04579 | 0.23907 |
| NAOrn 18:2;O | -0.2414 | 0.239235 |
| PE 34:2;2O\|PE 16:0_18:2;2O | -0.13192 | 0.239532 |
| PI 34:3\|PI 16:1_18:2 | 0.151811 | 0.240274 |
| Cer 44:1;3O\|Cer 26:0;2O/18:1;O | -0.21491 | 0.245941 |
| PE 40:3 | -0.15883 | 0.246051 |
| PE 36:4 | 0.074807 | 0.246271 |
| PS 36:6\|PS 18:3_18:3 | 0.187155 | 0.246324 |
| PC O-38:2 | 0.068061 | 0.249247 |
| PE 32:0\|PE 16:0_16:0 | -0.07665 | 0.249962 |
| PE-Cer 38:2;2O | -0.15233 | 0.250961 |
| PC 37:1 | 0.145683 | 0.251343 |
| SM 44:7;2O\|SM 14:1;2O/30:6 | 0.089847 | 0.251354 |
| CE 18:1 | -0.31536 | 0.252057 |
| ST 27:2;O | -0.09867 | 0.253841 |
| PC 32:2\|PC 16:1_16:1 | 0.08673 | 0.255268 |
| DG O-35:1\|DG O-19:0_16:1 | -0.67577 | 0.255449 |
| PC 31:0\|PC 15:0_16:0 | 0.131082 | 0.257281 |
| TG 41:0\|TG 12:0_14:0_15:0 | 0.39398 | 0.260267 |
| DG 30:0 | -0.84929 | 0.26063 |
| PG 36:2\|PG 18:0_18:2 | -0.09842 | 0.260772 |

| | | |
|---|---|---|
| PE O-36:3\|PE O-18:1_18:2 | -0.07424 | 0.261998 |
| NAE 22:4 align_ID:1090 | -0.55372 | 0.268079 |
| DG 36:4\|DG 18:2_18:2 | -0.11252 | 0.269802 |
| TG 55:3;O2\|TG 16:0_16:1_8:0;O(FA 15:1) | -0.47956 | 0.270311 |
| DG 32:0\|DG 16:0_16:0 | -0.28138 | 0.272738 |
| PE 34:2;3O\|PE 16:1_18:1;3O | -0.18826 | 0.275817 |
| PC O-36:9 | 0.364558 | 0.277233 |
| Cer 43:1;4O | -0.20745 | 0.277549 |
| PE 36:2 | 0.069048 | 0.279126 |
| TG 49:1\|TG 15:0_16:0_18:1 | 0.291495 | 0.27993 |
| Cer 35:1;2O\|Cer 14:1;2O/21:0 | -0.63492 | 0.280924 |
| PC 35:1\|PC 18:0_17:1 | 0.126356 | 0.281167 |
| LPC 18:3/0:0 | -0.30512 | 0.282801 |
| PS 36:5\|PS 18:2_18:3 | 0.13369 | 0.283483 |
| PC 26:0 align_ID:2772 | 0.213605 | 0.285287 |
| PE O-37:0 | -0.06976 | 0.285692 |
| PC 34:3 | -0.05483 | 0.285962 |
| Cer 44:1;3O\|Cer 19:0;2O/25:1;O | -0.22788 | 0.287183 |
| FA 18:1 align_ID:352 | -0.2022 | 0.287233 |
| Cer 38:1;2O\|Cer 14:1;2O/24:0 | -0.08273 | 0.288241 |
| DG 37:5 | 0.142819 | 0.292884 |
| FA 24:0 align_ID:557 | -0.14945 | 0.295065 |
| TG 37:0\|TG 12:0_12:0_13:0 | 0.275728 | 0.295722 |
| SM 36:2;3O align_ID:3629 | 0.0858 | 0.299516 |
| FA 25:0 | -0.33084 | 0.302699 |
| TG 41:1\|TG 13:0_14:0_14:1 | 0.303325 | 0.304858 |
| FA 27:0 align_ID:639 | -0.31493 | 0.307262 |
| Cer 44:1;4O\|Cer 18:1;3O/26:0;(2OH) | -0.18198 | 0.30905 |
| PE 38:1\|PE 20:0_18:1 | 0.083576 | 0.310171 |
| TG 47:1\|TG 14:0_15:0_18:1 | 0.271984 | 0.310216 |
| FA 21:0 align_ID:460 | -0.26347 | 0.310984 |
| Cer 46:0;3O\|Cer 24:0;2O/22:0;O | -0.25731 | 0.311648 |
| PC O-36:1\|PC O-18:0_18:1 | 0.17177 | 0.311882 |
| SM 40:6;3O | -0.49247 | 0.312552 |
| DG 49:10 | -0.43793 | 0.314131 |
| FA 16:0 align_ID:301 | -0.28849 | 0.316193 |
| SE 28:1/18:1 | -0.153 | 0.31648 |
| TG 56:4\|TG 20:0_18:2_18:2 | -0.31139 | 0.316852 |
| Cer 42:1;4O | -0.19897 | 0.317009 |
| FA 16:1;O | -0.35257 | 0.319884 |
| PC 35:1 | 0.15041 | 0.322105 |
| CL 71:7 | -0.07915 | 0.32387 |
| TG 43:0\|TG 14:0_14:0_15:0 | 0.352205 | 0.324424 |
| FA 23:0 | -0.24262 | 0.324695 |
| TG 53:3;O2\|TG 14:0_16:1_8:0;O(FA 15:1) | -0.33552 | 0.329584 |
| PG 35:3 | -0.15366 | 0.330457 |

| | | |
|---|---|---|
| PE 35:0 | 0.057398 | 0.333136 |
| DG 52:6 | -0.07866 | 0.333476 |
| PC O-37:1 | 0.121179 | 0.334172 |
| FA 20:1 | -0.37632 | 0.334229 |
| FA 22:1 | -0.35244 | 0.336207 |
| PE 40:2 | -0.0919 | 0.342622 |
| Cer 42:1;3O\|Cer 21:1;2O/21:0;O | -0.22104 | 0.344778 |
| PC O-33:2 align_ID:3468 | -0.05094 | 0.344907 |
| PC 35:2\|PC 17:0_18:2 | 0.044644 | 0.345543 |
| FA 26:0 | -0.2091 | 0.346095 |
| PE 36:2\|PE 18:0_18:2 | 0.021357 | 0.346207 |
| FA 28:0 | -0.24669 | 0.348232 |
| PE 27:0 | -0.11525 | 0.350662 |
| PC O-39:9 | -0.31468 | 0.351882 |
| PC O-35:4 | -0.0379 | 0.353677 |
| PE-Cer 32:2;2O\|PE-Cer 14:2;2O/18:0 | -0.10443 | 0.354129 |
| FA 18:0 align_ID:361 | -0.10299 | 0.354551 |
| FA 36:0 | -0.16688 | 0.357519 |
| Cer 42:0;2O | -0.19368 | 0.35785 |
| PE 34:2\|PE 16:0_18:2 | 0.029107 | 0.358493 |
| PE-Cer 34:0;2O\|PE-Cer 14:0;2O/20:0 | 0.075628 | 0.358617 |
| PE 32:3\|PE 14:0_18:3 | -0.12721 | 0.360593 |
| ST 27:1;O | -0.30124 | 0.360972 |
| PE-Cer 36:2;2O\|PE-Cer 14:2;2O/22:0 | -0.07775 | 0.36135 |
| PC O-32:1 | 0.060791 | 0.363017 |
| FA 24:1 | -0.30558 | 0.363969 |
| Cer 42:1;2O\|Cer 18:1;2O/24:0 | -0.19558 | 0.364185 |
| Cer 64:13;4O\|Cer 46:9;3O(FA 18:3) | 0.095517 | 0.364284 |
| PE O-30:0\|PE O-16:0_14:0 | 0.094819 | 0.364711 |
| Cer 46:1;3O\|Cer 20:1;2O/26:0;O | -0.17197 | 0.366059 |
| PC 39:3 | 0.384411 | 0.367536 |
| PC 37:4 | -0.13889 | 0.367553 |
| TG 52:3\|TG 16:1_18:1_18:1 | -0.06709 | 0.368774 |
| ST 29:1;O align_ID:1448 | 0.782701 | 0.368917 |
| PE 38:1 | 0.054619 | 0.369293 |
| PC 31:2;O\|PC 16:1_15:1;O | -0.1167 | 0.372929 |
| DG 50:6 | -0.06297 | 0.373392 |
| PC 37:5 | -0.13225 | 0.374694 |
| FA 34:0 | -0.42267 | 0.376034 |
| FA 19:0 | -0.24596 | 0.380832 |
| PC 34:1\|PC 16:0_18:1 | 0.007949 | 0.383046 |
| TG 50:1;2O\|TG 17:0_17:0_16:1;2O | 0.095231 | 0.386267 |
| PE 34:2 | 0.059452 | 0.387886 |
| PE 34:0\|PE 16:0_18:0 | -0.05076 | 0.388871 |
| FA 19:2 | -0.44594 | 0.389982 |
| PE O-35:2\|PE O-17:0_18:2 | -0.06123 | 0.39139 |

| | | |
|---|---|---|
| PE 36:5 | -0.31156 | 0.392404 |
| Cer 46:1;4O | -0.19239 | 0.393691 |
| DG 32:2\|DG 16:1_16:1 | -0.27177 | 0.393921 |
| Cer 42:0;3O\|Cer 18:0;3O/24:0 | -0.19486 | 0.396115 |
| LPC 16:0 | -0.19124 | 0.397346 |
| FA 19:1 | -0.38727 | 0.399802 |
| CL 72:8\|CL 18:2_18:2_18:2_18:2 | -0.15098 | 0.401438 |
| FA 20:3 | -0.4489 | 0.401985 |
| DG 29:0\|DG 14:0_15:0 | -0.22627 | 0.402645 |
| FA 18:1;O | -0.28525 | 0.403081 |
| FA 17:0 | -0.27842 | 0.405597 |
| DG 40:5 | -0.38146 | 0.405993 |
| DG 20:0 | -0.3488 | 0.406497 |
| FA 18:3 align_ID:342 | -0.23858 | 0.408171 |
| NAE 9:0 | -0.13534 | 0.40876 |
| PC O-36:3 | -0.05133 | 0.408921 |
| CAR 24:1 | -0.29354 | 0.409173 |
| FA 18:2 align_ID:347 | -0.24939 | 0.410611 |
| CL 77:7\|CL 39:3_38:4 | -0.20399 | 0.41225 |
| PE O-40:1\|PE O-22:0_18:1 | 0.090202 | 0.417191 |
| TG 51:2\|TG 17:0_16:1_18:1 | 0.157024 | 0.418197 |
| TG 37:1;1O\|TG 10:0_14:0_13:1;1O | 0.207291 | 0.422565 |
| PC O-38:8 | -1.01346 | 0.423539 |
| DG 32:5 | 0.268189 | 0.424033 |
| FA 20:2 | -0.40141 | 0.424376 |
| PG 34:2\|PG 16:0_18:2 | 0.082304 | 0.424493 |
| FA 22:4 | -0.25787 | 0.424503 |
| DG 44:4 | -0.30496 | 0.4256 |
| CL 72:4\|CL 39:1_33:3 | 0.137735 | 0.426184 |
| DG 31:0\|DG 15:0_16:0 | -0.26344 | 0.426982 |
| DG 36:3\|DG 18:1_18:2 | 0.124544 | 0.427068 |
| TG 43:1\|TG 13:0_14:0_16:1 | 0.213335 | 0.427248 |
| Cer 54:11;4O\|Cer 12:2;2O/42:9;2O | 0.058129 | 0.428431 |
| CL 80:5\|CL 18:0_26:0_18:2_18:3 | -0.23401 | 0.429628 |
| FA 32:1 | -0.23583 | 0.43118 |
| PE-Cer 32:1;3O | -0.07322 | 0.43272 |
| Cer 44:0;3O\|Cer 20:0;3O/24:0 | -0.16539 | 0.435713 |
| PI 36:4\|PI 18:2_18:2 | -0.08204 | 0.437227 |
| Dioctyl phthalate | -0.2576 | 0.438006 |
| Cer 48:10;4O\|Cer 12:2;2O/36:8;2O | 0.014249 | 0.439375 |
| DG 26:1\|DG 12:0_14:1 | -0.09146 | 0.440552 |
| TG 39:1\|TG 12:0_13:0_14:1 | 0.204936 | 0.441627 |
| PC 33:2\|PC 15:0_18:2 | -0.02225 | 0.441945 |
| DG 32:1\|DG 16:0_16:1 | -0.14398 | 0.443157 |
| DG 28:2 | -0.36149 | 0.445089 |
| Cer 42:1;3O\|Cer 18:1;2O/24:0;O | -0.13348 | 0.445613 |

| | | |
|---|---|---|
| PG 34:3\|PG 16:0_18:3 | -0.09147 | 0.4457 |
| PC 34:4 | -0.12741 | 0.450326 |
| PG 36:3\|PG 18:1_18:2 | 0.072199 | 0.451032 |
| TG 49:2\|TG 15:0_16:1_18:1 | 0.15984 | 0.451068 |
| FA 34:1 | -0.18841 | 0.451107 |
| FA 18:1 align_ID:355 | -0.26023 | 0.451514 |
| TG 51:3\|TG 15:0_18:1_18:2 | -0.10487 | 0.452256 |
| DG 33:0\|DG 16:0_17:0 | -0.23497 | 0.45599 |
| SE 28:2/18:1 | -0.1094 | 0.457732 |
| PE-Cer 36:1;2O\|PE-Cer 18:0;2O/18:1 | -0.06433 | 0.45782 |
| TG 44:3\|TG 14:1_14:1_16:1 | -0.14279 | 0.461013 |
| PE-Cer 37:2;3O\|PE-Cer 14:1;2O/23:1;O | -0.06006 | 0.463639 |
| Dodecylbenzenesulfonic acid | -0.27642 | 0.467914 |
| PE-Cer 38:5;2O | -0.1109 | 0.468057 |
| DG 30:1\|DG 14:0_16:1 | -0.1499 | 0.472171 |
| DG 34:0\|DG 16:0_18:0 | -0.09525 | 0.476535 |
| FA 18:0 align_ID:367 | -0.28694 | 0.477969 |
| FA 18:2 align_ID:346 | -0.11163 | 0.478102 |
| DG 18:0 | -0.3163 | 0.478179 |
| PE O-38:2\|PE O-20:0_18:2 | -0.03152 | 0.480496 |
| PE-Cer 38:0;2O\|PE-Cer 14:0;2O/24:0 | -0.08486 | 0.481398 |
| PE-Cer 34:2;2O\|PE-Cer 14:2;2O/20:0 | -0.0592 | 0.485006 |
| TG 41:2;1O\|TG 10:0_16:0_15:2;1O | 0.172295 | 0.48744 |
| PC O-45:11 | 0.205064 | 0.492838 |
| DG 36:2\|DG 18:0_18:2 | -0.06422 | 0.494924 |
| PE 36:3\|PE 18:0_18:3 | -0.02313 | 0.494926 |
| DG O-36:2\|DG O-18:0_18:2 | -0.06653 | 0.496799 |
| PC O-39:4 | 0.301401 | 0.497174 |
| SM 42:9;2O | -0.25623 | 0.498162 |
| CAR 24:0 | -0.08425 | 0.49844 |
| PC O-32:2 | -0.25848 | 0.498547 |
| ST 29:1;O align_ID:1223 | 0.109848 | 0.499642 |
| DG 30:2\|DG 14:1_16:1 | -0.17292 | 0.499686 |
| TG 47:3\|TG 13:0_16:1_18:2 | -0.13792 | 0.500441 |
| TG 54:3\|TG 18:0_18:1_18:2 | -0.13004 | 0.501266 |
| Cer 43:0;3O | -0.14128 | 0.505224 |
| PC O-32:0 | -0.04204 | 0.505796 |
| Cer 42:0;4O\|Cer 18:0;3O/24:0;(2OH) | -0.14367 | 0.506298 |
| PC 37:3 | 0.057607 | 0.507083 |
| FA 34:4 | -0.31204 | 0.512305 |
| FA 15:0 | -0.27133 | 0.51294 |
| ST 27:1;O;S | -0.22309 | 0.513406 |
| Cer 40:1;2O\|Cer 18:1;2O/22:0 | -0.10608 | 0.513855 |
| PE-Cer 33:1;2O | -0.08676 | 0.515623 |
| MG 37:1 | 0.389484 | 0.516138 |
| PC 38:1 align_ID:4354 | -0.22186 | 0.516752 |

| | | |
|---|---|---|
| PE 30:2;O\|PE 16:1_14:1;O | -0.09127 | 0.519163 |
| PE O-36:2\|PE O-18:0_18:2 align_ID:1391 | 0.027968 | 0.519631 |
| FA 17:1 | -0.32666 | 0.520956 |
| FA 11:0 | -0.12029 | 0.520983 |
| PE O-36:2\|PE O-18:0_18:2 align_ID:1392 | 0.056803 | 0.521695 |
| DG 38:1\|DG 20:0_18:1 | -0.16059 | 0.522023 |
| PC 38:6\|PC 16:0_22:6 | -0.10432 | 0.524015 |
| PE-Cer 36:1;2O\|PE-Cer 14:1;2O/22:0 | 0.050527 | 0.527694 |
| DG 41:8 | 0.490812 | 0.529481 |
| FA 18:1 align_ID:353 | -0.20583 | 0.52954 |
| TG 45:0\|TG 14:0_15:0_16:0 | 0.14149 | 0.533578 |
| PC 38:5 | -0.09188 | 0.534044 |
| Cer 46:0;3O\|Cer 22:0;3O/24:0 | -0.16351 | 0.535035 |
| PG 36:1\|PG 18:0_18:1 | 0.083054 | 0.535126 |
| PC 32:2 | 0.036439 | 0.535668 |
| PC 35:3 | -0.0223 | 0.536497 |
| FA 30:1 | -0.17861 | 0.537769 |
| TG 62:1\|TG 14:0_16:0_32:1 | -0.1169 | 0.538692 |
| Cer 44:0;2O | -0.17419 | 0.538856 |
| PE 34:3\|PE 16:0_18:3 | -0.03509 | 0.539852 |
| Cer 34:1;4O | -0.17827 | 0.540548 |
| PE 35:0\|PE 17:0_18:0 | 0.070361 | 0.540739 |
| DG 49:13 | -0.3313 | 0.541079 |
| PG 36:4\|PG 18:2_18:2 | -0.07707 | 0.542171 |
| TG 45:1\|TG 14:0_15:0_16:1 | 0.160329 | 0.543139 |
| TG 56:1\|TG 18:0_20:0_18:1 | -0.1366 | 0.546316 |
| MG 18:1 | -0.22254 | 0.547686 |
| TG 60:1\|TG 14:0_14:0_32:1 | -0.08702 | 0.550201 |
| CL 78:6\|CL 24:0_18:2_18:2_18:2 | -0.08312 | 0.550702 |
| DG 36:5 | -0.2726 | 0.55165 |
| ST 29:2;O | -0.14583 | 0.55181 |
| FA 15:1 | -0.35121 | 0.553148 |
| Cer 42:1;3O\|Cer 19:0;2O/23:1;O | -0.09671 | 0.55346 |
| TG 56:2\|TG 14:0_18:1_24:1 | 0.100139 | 0.55375 |
| TG O-50:2\|TG O-18:0_16:1_16:1 | 0.174763 | 0.553897 |
| TG 40:0\|TG 12:0_14:0_14:0 | 0.100827 | 0.556499 |
| DG 42:10 | -0.23129 | 0.560619 |
| DG 43:10 | -0.2431 | 0.562415 |
| DG 31:3 | -0.25451 | 0.564527 |
| TG 38:0\|TG 12:0_12:0_14:0 | 0.129559 | 0.564954 |
| TG 56:3\|TG 20:0_18:1_18:2 | -0.1335 | 0.567444 |
| TG 50:1\|TG 16:0_16:0_18:1 | 0.1067 | 0.569558 |
| PE-Cer 38:3;2O | -0.09946 | 0.571013 |
| FA 10:0 | 0.055739 | 0.572513 |
| PC 35:2 | -0.07264 | 0.572951 |
| FA 14:0 | -0.25239 | 0.573124 |

| | | |
|---|---|---|
| DG 34:2\|DG 16:0_18:2 | -0.09938 | 0.57396 |
| TG 54:2\|TG 18:0_18:1_18:1 | -0.08744 | 0.578155 |
| PE 36:1;2O\|PE 18:1_18:0;2O | 0.042663 | 0.579708 |
| PC O-37:3 | -0.03858 | 0.580886 |
| PE 36:3;O\|PE 18:2_18:1;O | -0.16663 | 0.584125 |
| PE 38:0\|PE 18:0_20:0 | -0.07416 | 0.586768 |
| FA 16:2 | -0.33963 | 0.588557 |
| PE O-36:3 | -0.06911 | 0.589423 |
| SM 31:6;2O | -0.13788 | 0.597345 |
| ST 28:4;O align_ID:1098 | -0.12167 | 0.600628 |
| TG 49:0\|TG 15:0_16:0_18:0 | -0.0878 | 0.600631 |
| TG 36:0\|TG 12:0_12:0_12:0 | 0.134671 | 0.601058 |
| DG 24:0 | -0.08042 | 0.602351 |
| FA 32:2 | -0.17118 | 0.602391 |
| LPC 16:1 | 0.277338 | 0.603583 |
| TG 48:0\|TG 14:0_16:0_18:0 | -0.09136 | 0.603922 |
| DG 26:1 | -0.22389 | 0.60456 |
| PC 38:2 align_ID:4335 | -0.28177 | 0.606155 |
| FA 42:0 | -0.07113 | 0.606775 |
| PG 34:4 | -0.07024 | 0.60746 |
| TG 39:2;1O\|TG 10:0_16:1_13:1;1O | 0.12319 | 0.610552 |
| TG 47:0\|TG 15:0_16:0_16:0 | -0.10268 | 0.611206 |
| TG 42:0\|TG 14:0_14:0_14:0 | 0.102353 | 0.612149 |
| FA 26:1 | -0.09728 | 0.616 |
| TG 34:0\|TG 10:0_12:0_12:0 | -0.20144 | 0.619349 |
| PE-Cer 34:0;2O | -0.03319 | 0.619497 |
| PE-Cer 32:0;2O\|PE-Cer 14:0;2O/18:0 | 0.061198 | 0.619998 |
| SM 63:7;2O | -0.03145 | 0.621031 |
| DG 33:6 | -0.62883 | 0.621362 |
| PE O-36:0\|PE O-18:0_18:0 | -0.04914 | 0.625101 |
| PE 36:2\|PE 18:1_18:1 | -0.05672 | 0.625344 |
| PE O-36:4 | 0.028139 | 0.627109 |
| TG 48:3\|TG 16:1_16:1_16:1 | -0.06904 | 0.627576 |
| PE-Cer 36:0;2O\|PE-Cer 14:0;2O/22:0 | 0.042905 | 0.628766 |
| TG 52:2\|TG 16:0_18:1_18:1 | 0.067805 | 0.62892 |
| TG 48:1\|TG 14:0_16:0_18:1 | 0.10851 | 0.629702 |
| PE-Cer 33:1;3O\|PE-Cer 14:0;2O/19:1;O | -0.06391 | 0.630539 |
| CL 78:5\|CL 18:0_24:0_18:2_18:3 | -0.19785 | 0.631421 |
| LPE 18:1 align_ID:1741 | 0.2475 | 0.634218 |
| FA 16:1 | -0.2378 | 0.636901 |
| PE 36:3 | 0.044966 | 0.636904 |
| TG O-38:0\|TG O-14:0_12:0_12:0 | -0.14588 | 0.646434 |
| TG 62:2\|TG 14:0_16:1_32:1 | 0.102375 | 0.648692 |
| PE O-40:2\|PE O-22:0_18:2 | 0.032349 | 0.651115 |
| PE 34:4 | 0.082862 | 0.654762 |
| PE O-34:3 | -0.02134 | 0.656251 |

| | | |
|---|---|---|
| TG 46:0\|TG 14:0_16:0_16:0 | -0.08362 | 0.657722 |
| DG 28:1\|DG 14:0_14:1 | -0.06457 | 0.659469 |
| PC 28:0 | -0.02376 | 0.661462 |
| PC 32:4\|PC 16:1_16:3 | 0.039975 | 0.661598 |
| ST 28:3;O align_ID:1116 | -0.08285 | 0.66178 |
| NAE 4:0 | -0.13886 | 0.664841 |
| PC 36:2 align_ID:4091 | -0.15472 | 0.664988 |
| PC O-29:3 | 0.15985 | 0.665331 |
| PC 38:7 | -0.08886 | 0.665347 |
| TG 58:3\|TG 16:1_18:1_24:1 | -0.09793 | 0.666387 |
| FA 21:0 align_ID:459 | -0.08818 | 0.666599 |
| TG 28:0\|TG 8:0_10:0_10:0 | -0.43873 | 0.669297 |
| TG 38:1\|TG 12:0_12:0_14:1 | 0.113412 | 0.673965 |
| FA 13:0 | -0.22589 | 0.676105 |
| PC 30:2 | 0.049017 | 0.677455 |
| TG 49:2;2O\|TG 14:0_14:0_21:2;2O | -0.17172 | 0.67894 |
| TG 40:1\|TG 12:0_12:0_16:1 | 0.091016 | 0.679864 |
| LPE 16:1 | 0.267179 | 0.683223 |
| FA 14:1 | -0.25268 | 0.683981 |
| PC O-40:8 | -0.06454 | 0.684742 |
| PE 32:3 | -0.04886 | 0.688353 |
| PI 34:4\|PI 16:1_18:3 | 0.059486 | 0.6892 |
| PC O-35:3 align_ID:3729 | -0.03052 | 0.691658 |
| Norethisterone acetate | -0.16981 | 0.6917 |
| PE 36:0\|PE 18:0_18:0 | -0.04025 | 0.693393 |
| TG 24:0\|TG 8:0_8:0_8:0 | -0.36247 | 0.694822 |
| PC O-35:0 | 0.043262 | 0.694845 |
| TG 44:0\|TG 14:0_14:0_16:0 | 0.080957 | 0.695666 |
| PC O-38:9 | 0.33727 | 0.695928 |
| Cer 34:0;3O\|Cer 14:0;3O/20:0 | -0.03002 | 0.69957 |
| PE-Cer 37:1;2O | -0.0415 | 0.700058 |
| MGDG 35:2 | -0.11516 | 0.702388 |
| FA 9:0 align_ID:131 | -0.12796 | 0.703559 |
| CL 76:3\|CL 16:0_24:0_18:1_18:2 | 0.065159 | 0.705825 |
| DG 26:4 | -0.1151 | 0.706338 |
| PE 30:1\|PE 14:0_16:1 | 0.074363 | 0.708945 |
| PE-Cer 34:1;2O\|PE-Cer 14:1;2O/20:0 | -0.0261 | 0.710071 |
| CL 59:0 | -0.19073 | 0.710207 |
| TG 50:2\|TG 16:0_16:1_18:1 | 0.064455 | 0.710703 |
| PC 36:3 | 0.028396 | 0.713012 |
| Cer 48:0;3O | -0.08601 | 0.716818 |
| TG 26:0\|TG 8:0_8:0_10:0 | -0.33338 | 0.717641 |
| PE P-36:4\|PE P-18:2_18:2 | -0.03358 | 0.718822 |
| TG O-40:0\|TG O-14:0_12:0_14:0 | -0.22127 | 0.720513 |
| SM 42:8;2O | 0.150482 | 0.723462 |
| PI 36:4 | 0.02819 | 0.725292 |

| | | |
|---|---|---|
| PC O-38:4 | 0.431084 | 0.726517 |
| LPC 18:0/0:0 | -0.0843 | 0.729578 |
| PC O-33:1 | -0.00788 | 0.735292 |
| DGGA 32:0\|DGGA 16:0_16:0 | 0.031449 | 0.736472 |
| Cer 47:5;4O\|Cer 29:2;3O(FA 18:2) | -0.40962 | 0.74001 |
| TG O-42:0\|TG O-16:0_12:0_14:0 | 0.079718 | 0.741334 |
| ST 28:4;O align_ID:1100 | -0.05197 | 0.741832 |
| PE O-40:5\|PE O-22:3_18:2 | -0.05862 | 0.742654 |
| CL 80:3\|CL 18:0_26:0_18:1_18:2 | -0.09394 | 0.742886 |
| FA 29:0 | -0.06931 | 0.743512 |
| NAE 5:0 | 0.084363 | 0.746465 |
| TG 58:2\|TG 16:0_18:1_24:1 | -0.07776 | 0.747488 |
| LPC 18:2/0:0 | -0.11726 | 0.747592 |
| LPE 18:1 align_ID:793 | 0.171161 | 0.748486 |
| PE-Cer 38:1;2O\|PE-Cer 14:1;2O/24:0 | 0.021488 | 0.749194 |
| DG 35:5 | -0.08803 | 0.749502 |
| NAE 22:6 | 0.156033 | 0.749942 |
| SM 39:8;3O | -0.03679 | 0.74999 |
| TG 54:1\|TG 18:0_18:0_18:1 | -0.04584 | 0.753509 |
| PC 35:5 | 0.055703 | 0.757862 |
| LPC 18:1/0:0 | 0.10003 | 0.760409 |
| FA 18:0 align_ID:365 | 0.058415 | 0.760512 |
| PE O-34:1\|PE O-17:0_17:1 | 0.017743 | 0.761277 |
| PE 35:3\|PE 17:1_18:2 | 0.024295 | 0.762083 |
| 4-Imidazoleacrylic acid | -0.18525 | 0.764263 |
| PC O-33:0 | -0.02633 | 0.764771 |
| Cer 40:1;4O | -0.27959 | 0.765152 |
| PE 40:1\|PE 22:0_18:1 | -0.03733 | 0.768302 |
| TG 64:2\|TG 16:0_16:1_32:1 | -0.1041 | 0.768404 |
| DG 28:0\|DG 14:0_14:0 | 0.040532 | 0.768435 |
| PE-Cer 30:1;2O | -0.0565 | 0.76852 |
| TG 50:0\|TG 16:0_16:0_18:0 | -0.04552 | 0.775734 |
| PE 40:2\|PE 22:0_18:2 | -0.03013 | 0.777401 |
| TG 46:1\|TG 14:0_16:0_16:1 | 0.048067 | 0.780963 |
| SM 41:0;3O | 0.017591 | 0.786645 |
| LPC 18:1 | 0.096935 | 0.789255 |
| PC O-38:10 | -0.11659 | 0.790918 |
| SM 40:7;2O | 0.103693 | 0.790922 |
| TG 46:2\|TG 14:0_16:1_16:1 | -0.03608 | 0.794847 |
| PC 34:2\|PC 16:1_18:1 | -0.00253 | 0.795382 |
| PC 33:1\|PC 15:0_18:1 | -0.00684 | 0.797851 |
| FA 34:3 | -0.09239 | 0.799867 |
| LPC 18:2 | -0.08552 | 0.805121 |
| DG 34:4\|DG 16:1_18:3 | 0.022231 | 0.805341 |
| NAE 19:0 | -0.06307 | 0.80574 |
| PE O-35:2 | -0.01135 | 0.806877 |

| | | |
|---|---|---|
| PE O-34:3\|PE O-16:1_18:2 | 0.025059 | 0.814011 |
| PE 34:4\|PE 16:1_18:3 | 0.032004 | 0.814057 |
| PE 35:2;O\|PE 18:1_17:1;O | 0.033476 | 0.824592 |
| PC O-35:2 | -0.0096 | 0.824664 |
| TG 42:1\|TG 12:0_14:0_16:1 | 0.035917 | 0.827815 |
| PC 30:3 | -0.01609 | 0.828659 |
| TG 50:3\|TG 16:1_16:1_18:1 | 0.024547 | 0.830282 |
| TG 44:2\|TG 12:0_16:1_16:1 | -0.03139 | 0.83037 |
| PC 34:4\|PC 16:1_18:3 | -0.01657 | 0.830622 |
| DG 46:10 | 0.018544 | 0.83198 |
| TG 52:1\|TG 16:0_18:0_18:1 | 0.035064 | 0.833926 |
| DG 24:0\|DG 12:0_12:0 | -0.03031 | 0.835084 |
| PC 34:3\|PC 16:1_18:2 | 0.009586 | 0.840221 |
| LPC 16:0/0:0 | -0.04256 | 0.840819 |
| DG 50:5 | -0.01449 | 0.841699 |
| PC 36:1 | 0.06366 | 0.844086 |
| DG 22:1 | 0.229064 | 0.845795 |
| DG O-35:2\|DG O-21:1_14:1 | -0.18123 | 0.846435 |
| PC O-37:2 | 0.015034 | 0.849487 |
| PE O-40:7\|PE O-22:5_18:2 | -0.02234 | 0.855145 |
| TG 52:0\|TG 16:0_18:0_18:0 | 0.033659 | 0.855734 |
| PE O-37:2\|PE O-19:0_18:2 | 0.020049 | 0.860512 |
| ST 28:1;O | 0.233068 | 0.860824 |
| SE 29:1/18:1 | -0.03305 | 0.8612 |
| LPE O-18:0 | -0.0308 | 0.861474 |
| DG O-39:2\|DG O-19:1_20:1 | -0.13888 | 0.868017 |
| DG 31:1\|DG 15:0_16:1 | 0.039653 | 0.881623 |
| LPE 16:0 | 0.039668 | 0.883277 |
| DG 30:3 | 0.05157 | 0.887528 |
| PC O-41:7 | 0.056869 | 0.889632 |
| PC 33:0\|PC 16:0_17:0 | -0.01421 | 0.89007 |
| PE O-34:3\|PE O-16:0_18:3 | 0.006109 | 0.894274 |
| DG 25:1 | 0.055745 | 0.896986 |
| MG 18:0 | 0.018773 | 0.899977 |
| PE 35:2\|PE 17:0_18:2 | -0.00818 | 0.90034 |
| SM 42:7;2O | 0.047724 | 0.901609 |
| TG 45:2;1O\|TG 16:0_16:1_13:1;1O | -0.22355 | 0.901879 |
| TG 48:2\|TG 14:0_16:1_18:1 | 0.019647 | 0.903987 |
| DG 28:4 | -0.05525 | 0.904764 |
| PI 34:4 | 0.013837 | 0.904857 |
| PC O-39:1 | 0.040825 | 0.907689 |
| SM 37:1;2O\|SM 21:0;2O/16:1 | 0.013607 | 0.909227 |
| TG 54:1\|TG 12:0_14:0_28:1 | 0.01901 | 0.909511 |
| TG 45:2\|TG 13:0_16:1_16:1 | 0.021405 | 0.914779 |
| Cer 48:5;4O\|Cer 30:4;3O(FA 18:0) | -0.10507 | 0.916499 |
| LPE 18:2 | -0.03706 | 0.917257 |

| | | |
|---|---|---|
| PC 40:7 | 0.013675 | 0.92054 |
| TG 54:0\|TG 16:0_18:0_20:0 | 0.012418 | 0.924898 |
| FA 16:0 align_ID:298 | -0.01504 | 0.924943 |
| PC O-39:2 | -0.00656 | 0.925527 |
| TG 42:2\|TG 12:0_14:1_16:1 | -0.01975 | 0.926162 |
| SHexCer 33:2;3O | 0.013255 | 0.932926 |
| DG 34:1\|DG 16:0_18:1 | -0.01415 | 0.936279 |
| TG 44:1\|TG 14:0_14:0_16:1 | -0.01144 | 0.936338 |
| PC 34:2 | 0.002548 | 0.9368 |
| LPE O-20:0 | 0.010597 | 0.937447 |
| TG 43:2\|TG 13:0_14:1_16:1 | -0.01927 | 0.938098 |
| FA 12:0 | -0.02779 | 0.938775 |
| CL 78:3\|CL 18:0_26:0_16:1_18:2 | 0.024079 | 0.946435 |
| PE O-37:5\|PE O-19:2_18:3 | 0.007651 | 0.952785 |
| CL 72:7\|CL 18:1_18:2_18:2_18:2 | 0.008913 | 0.957947 |
| DG 41:6 | -0.04586 | 0.958399 |
| DG 48:12 | 0.017253 | 0.967201 |
| TG 50:5\|TG 16:1_16:1_18:3 | -0.01281 | 0.967699 |
| PE-Cer 34:1;3O\|PE-Cer 18:1;2O/16:0;O | 0.003198 | 0.968084 |
| TG 40:2\|TG 12:0_14:1_14:1 | 0.009919 | 0.97064 |
| PE O-38:0\|PE O-18:0_20:0 | -0.0035 | 0.975837 |
| PE O-36:5\|PE O-18:3_18:2 | -0.00361 | 0.977047 |
| DG 33:1\|DG 15:0_18:1 | -0.00725 | 0.978197 |
| DG 36:1\|DG 18:0_18:1 | 0.003433 | 0.982997 |
| PE-Cer 32:0;3O | 0.001775 | 0.984879 |
| PC O-39:7 | 0.008093 | 0.985322 |
| CL 82:9\|CL 28:0_18:3_18:3_18:3 | -0.00316 | 0.988674 |
| PC 31:1 | -0.0003 | 0.989778 |
| LPE 18:0 | -0.00156 | 0.993025 |
| PE O-34:2\|PE O-16:0_18:2 | 0.000127 | 0.99819 |
| Cer 36:0;3O\|Cer 14:0;3O/22:0 | -0.00015 | 0.998424 |
| SM 36:2;3O align_ID:3627 | 0 | 1 |
| NAE 14:0 | NA | NA |
| **Differentially Regulated Lipids in Male brains with Neuronal loss of *dATGL*** | | |
| **Lipid identity** | **log2FC (RNAi/controls)** | **p-value** |
| CAR 22:0 | -0.54715 | 1.68E-06 |
| PC 36:4\|PC 18:2_18:2 | -0.10813 | 1.24E-05 |
| PE O-34:0\|PE O-18:0_16:0 | -0.39202 | 2.43E-05 |
| PE O-32:0\|PE O-18:0_14:0 | -0.50948 | 3.14E-05 |
| PE 36:5\|PE 18:2_18:3 | -0.52496 | 4.91E-05 |
| PG 36:5\|PG 18:2_18:3 | -0.61675 | 6.07E-05 |
| PC 36:5 | -0.66222 | 7.57E-05 |
| PE O-34:0 | -0.35375 | 0.000114 |
| PE 34:2\|PE 16:0_18:2 | -0.08938 | 0.000116 |

| | | |
|---|---|---|
| PS 36:3 | 0.670622 | 0.000138 |
| PE 37:1 | 0.351185 | 0.000333 |
| PC 32:0\|PC 16:0_16:0 | -0.31929 | 0.000378 |
| PG 34:3\|PG 16:0_18:3 | -0.38649 | 0.000546 |
| PE 32:0\|PE 16:0_16:0 | -0.37795 | 0.000563 |
| PC 30:0\|PC 14:0_16:0 | -0.36755 | 0.000585 |
| DG 44:9 | 0.385632 | 0.000672 |
| CL 76:5\|CL 24:0_16:1_18:1_18:3 | -0.60597 | 0.000742 |
| PS 36:3\|PS 18:1_18:2 | 0.563177 | 0.000788 |
| PS 36:2 align_ID:4110 | 0.756556 | 0.001006 |
| PI 34:1\|PI 16:0_18:1 | 0.793035 | 0.00101 |
| PC 35:1\|PC 17:0_18:1 | 0.535885 | 0.00101 |
| CL 77:7\|CL 39:3_38:4 | -0.83739 | 0.001129 |
| PG 36:4\|PG 18:2_18:2 | -0.48363 | 0.001351 |
| PE 34:3\|PE 16:0_18:3 | -0.1349 | 0.001393 |
| PE 32:3\|PE 14:0_18:3 | -0.4716 | 0.001423 |
| PI 34:1 | 0.715413 | 0.001432 |
| PC 32:3 | -0.48812 | 0.001453 |
| PS 34:4\|PS 16:1_18:3 | 0.695011 | 0.001462 |
| PE 34:0\|PE 16:0_18:0 | -0.2184 | 0.001538 |
| CL 68:6\|CL 16:1_18:2_16:1_18:2 | 0.565916 | 0.001676 |
| CL 68:6\|CL 16:1_16:1_18:2_18:2 | 0.563944 | 0.001712 |
| PS 34:2\|PS 16:0_18:2 | 0.735066 | 0.00187 |
| PE 32:2 | 0.801329 | 0.002078 |
| PI 36:5\|PI 18:2_18:3 | -0.59782 | 0.002186 |
| PC 34:0\|PC 16:0_18:0 | -0.28294 | 0.002337 |
| PS 34:3\|PS 16:1_18:2 | 0.856749 | 0.00243 |
| PE 36:3\|PE 18:0_18:3 | -0.13097 | 0.002467 |
| PI 32:1\|PI 16:0_16:1 | 0.686367 | 0.00253 |
| PS 36:2\|PS 18:1_18:1 | 0.946887 | 0.002544 |
| PC 37:8 | -0.52738 | 0.00256 |
| PC 40:8 | -0.52643 | 0.002803 |
| PE 34:3 | 0.281413 | 0.002937 |
| PE 36:6\|PE 18:3_18:3 | -0.57877 | 0.003153 |
| TG 37:0\|TG 12:0_12:0_13:0 | 0.652694 | 0.003264 |
| HexCer 34:2;2O | -0.42074 | 0.003355 |
| PC 36:2\|PC 18:1_18:1 | -0.05115 | 0.003404 |
| TG 50:4\|TG 16:1_16:1_18:2 | -0.5006 | 0.00345 |
| DG 32:5 | -0.80461 | 0.003477 |
| PC 36:1\|PC 18:0_18:1 | 0.226395 | 0.003488 |
| CL 71:7 | -0.2542 | 0.003801 |
| PC 36:5\|PC 18:2_18:3 | -0.2631 | 0.003845 |
| LPE 16:0 | -0.36097 | 0.00395 |
| PI 32:1 | 0.696908 | 0.004013 |
| PS 36:5 | 0.433708 | 0.004052 |
| PE 36:0\|PE 18:0_18:0 | -0.27804 | 0.004139 |

| | | |
|---|---|---|
| PS 34:1\|PS 16:0_18:1 | 1.216664 | 0.004297 |
| PC O-34:1 | 0.663149 | 0.004408 |
| Cer 48:10;4O\|Cer 12:2;2O/36:8;2O | -0.04393 | 0.004415 |
| PC 38:1 align_ID:4355 | 0.431998 | 0.004618 |
| PS 36:1\|PS 18:0_18:1 | 1.297223 | 0.004669 |
| SM 41:0;3O | 0.266872 | 0.005002 |
| PE 36:2;O\|PE 18:2_18:0;O | -2.30204 | 0.005054 |
| PS 36:3\|PS 10:0_26:3 | 0.596489 | 0.00507 |
| PE 33:2 | 0.414701 | 0.005138 |
| PS 36:4\|PS 18:2_18:2 | 0.346615 | 0.00529 |
| PC O-36:1 | 0.609147 | 0.005354 |
| PS 36:2 align_ID:4113 | 1.113292 | 0.005529 |
| DG 32:2 | -1.00931 | 0.005603 |
| Cer 47:9;4O\|Cer 15:2;2O/32:7;2O | 0.550341 | 0.00569 |
| PE 34:2;2O\|PE 16:0_18:2;2O | -0.90334 | 0.00572 |
| PE 33:1\|PE 16:0_17:1 | 0.395206 | 0.006297 |
| PC 33:1 | 0.422521 | 0.006351 |
| PC 35:4 | -0.38208 | 0.006358 |
| PE 36:1;O\|PE 18:0_18:1;O | -2.52737 | 0.006563 |
| CL 73:6 | -0.27371 | 0.006598 |
| PE O-38:3\|PE O-20:0_18:3 | -0.20288 | 0.007129 |
| TG 39:0\|TG 12:0_13:0_14:0 | 0.85653 | 0.007215 |
| CAR 20:0 | -0.39879 | 0.007247 |
| PE 36:4;2O\|PE 18:2_18:2;2O | -0.83087 | 0.007271 |
| PC O-40:9 | -0.85573 | 0.007281 |
| SM 44:7;2O\|SM 14:1;2O/30:6 | 0.195819 | 0.007283 |
| PC 36:6\|PC 18:3_18:3 | -0.48351 | 0.007366 |
| CAR 24:1 | -1.25341 | 0.007442 |
| PC 33:2\|PC 15:0_18:2 | -0.12162 | 0.007614 |
| PE 36:4 | 0.23132 | 0.007708 |
| PC 36:4 | -0.23348 | 0.007769 |
| PE O-32:1 | 0.199885 | 0.007776 |
| PC O-33:0 | -0.30921 | 0.007861 |
| PE-Cer 32:1;2O\|PE-Cer 14:1;2O/18:0 | -0.42767 | 0.00789 |
| PE 36:4\|PE 18:2_18:2 | -0.19966 | 0.007904 |
| CL 66:5\|CL 16:1_16:1_16:1_18:2 | 1.123387 | 0.008393 |
| SM 31:6;2O | -0.45409 | 0.008445 |
| PC 26:0 align_ID:2773 | 0.953746 | 0.008468 |
| PE O-40:3\|PE O-22:0_18:3 | -0.27557 | 0.008605 |
| TG 46:4\|TG 12:0_16:1_18:3 | -0.48284 | 0.008616 |
| PE P-36:4\|PE P-18:2_18:2 | -0.34808 | 0.008842 |
| PE 33:2;2O\|PE 16:1_17:1;2O | 1.083079 | 0.009548 |
| LPE O-18:0 | -0.28735 | 0.010577 |
| PC 33:2 | 0.228684 | 0.010755 |
| TG 39:2;1O\|TG 10:0_16:1_13:1;1O | -0.69383 | 0.010868 |
| PE O-34:3 | -0.11498 | 0.01177 |

| | | |
|---|---|---|
| PE 35:2;O\|PE 18:1_17:1;O | -0.43537 | 0.012039 |
| PC O-45:11 | -0.72359 | 0.012056 |
| PI 34:0\|PI 16:0_18:0 | 0.773465 | 0.012058 |
| LPE 18:0 | -0.26807 | 0.012158 |
| DG 44:7 | 0.273031 | 0.012481 |
| PE 35:1;2O\|PE 18:0_17:1;2O | 1.168577 | 0.012521 |
| PC 28:1 | 0.481407 | 0.013432 |
| PC 34:1\|PC 16:0_18:1 | 0.037129 | 0.013512 |
| PE O-36:6\|PE O-18:4_18:2 | -0.31921 | 0.013624 |
| PS 36:5\|PS 18:2_18:3 | 0.191859 | 0.013696 |
| PC 34:0 | 0.419836 | 0.013748 |
| PI 36:5 | -0.4219 | 0.014763 |
| DG 28:2 | -0.50096 | 0.014764 |
| CL 70:6\|CL 16:0_18:2_18:2_18:2 | 0.678855 | 0.01517 |
| PE 36:2;2O\|PE 18:0_18:2;2O | -0.7518 | 0.015189 |
| CL 70:6\|CL 16:1_18:1_18:2_18:2 | 0.632378 | 0.015263 |
| PE O-32:1\|PE O-16:0_16:1 | 0.397859 | 0.015274 |
| TG 27:0\|TG 9:0_9:0_9:0 | -0.55725 | 0.015308 |
| PI 36:1\|PI 18:0_18:1 | 0.941542 | 0.015612 |
| PE O-36:3\|PE O-18:0_18:3 | -0.17617 | 0.015704 |
| DG 36:1\|DG 18:0_18:1 | 0.319342 | 0.015793 |
| PS 34:3 | 0.904349 | 0.01586 |
| PG 36:2\|PG 18:1_18:1 | -0.44546 | 0.015873 |
| PC 34:4 | -0.79692 | 0.016439 |
| DG 27:0 | 0.837988 | 0.016453 |
| PC O-32:1\|PC O-16:0_16:1 | 0.515609 | 0.016699 |
| PC 37:1 | 0.498519 | 0.016938 |
| PI 36:6 | -0.62253 | 0.017224 |
| PE O-36:0\|PE O-18:0_18:0 | -0.29423 | 0.017753 |
| TG 52:1;4O\|TG 14:0_16:0_22:1;4O | -1.13488 | 0.017878 |
| PC O-34:1\|PC O-18:0_16:1 | 0.45212 | 0.019268 |
| PE P-34:1\|PE P-18:0_16:1 | 0.270083 | 0.020119 |
| PE O-34:2\|PE O-18:1_16:1 | 0.230921 | 0.020217 |
| NAOrn 18:2;O | -0.52395 | 0.020229 |
| PI 36:2 | 0.399395 | 0.02085 |
| TG 36:0\|TG 12:0_12:0_12:0 | 0.473723 | 0.021009 |
| PE 32:2\|PE 16:1_16:1 | 0.414492 | 0.021063 |
| PC 35:2\|PC 17:0_18:2 | 0.214927 | 0.021156 |
| PG 34:2\|PG 16:0_18:2 | -0.27569 | 0.021236 |
| PC 32:0 | 0.403258 | 0.021901 |
| PC O-36:4 | -0.22107 | 0.022075 |
| PC 33:0 | 0.255177 | 0.02243 |
| PE O-36:3\|PE O-18:1_18:2 | -0.23298 | 0.023542 |
| TG O-42:0\|TG O-16:0_12:0_14:0 | -0.58886 | 0.023718 |
| PC 38:2 align_ID:4336 | -0.24451 | 0.02391 |
| FA 16:0 align_ID:301 | -0.40937 | 0.024131 |

| | | |
|---|---|---|
| PE 34:3\|PE 16:1_18:2 | 0.155012 | 0.024667 |
| Cer 48:11;4O\|Cer 13:1;2O/35:10;2O | -0.12621 | 0.024893 |
| PC O-40:8 | -0.64584 | 0.02595 |
| PE O-36:7\|PE O-18:4_18:3 | -0.38512 | 0.026414 |
| PC 34:4\|PC 18:1_16:3 | -0.1267 | 0.026533 |
| PC O-37:3 | -0.17746 | 0.026542 |
| PE 25:1;O\|PE 10:0_15:1;O | -0.33426 | 0.027318 |
| PS 32:2 | 0.940519 | 0.027782 |
| PS 34:4 | 0.962839 | 0.02798 |
| TG 41:2;1O\|TG 10:0_16:0_15:2;1O | -0.70972 | 0.028639 |
| NAE 9:0 | -0.3928 | 0.02879 |
| TG 49:2;2O\|TG 14:0_14:0_21:2;2O | 0.570477 | 0.029374 |
| PS 34:2 | 1.045597 | 0.029627 |
| PI 36:2\|PI 18:1_18:1 | 0.344755 | 0.029857 |
| FA 30:2 | -0.3863 | 0.029936 |
| PC 36:3 | 0.160932 | 0.030534 |
| LPC 18:1 | 0.461204 | 0.030676 |
| PE O-36:4\|PE O-18:1_18:3 | -0.37456 | 0.032956 |
| PE O-36:1\|PE O-18:0_18:1 | 0.06964 | 0.033535 |
| PE-Cer 32:2;2O\|PE-Cer 14:2;2O/18:0 | -0.3606 | 0.034496 |
| Cer 34:2;2O\|Cer 14:2;2O/20:0 | -0.22317 | 0.034541 |
| DG 35:1\|DG 17:0_18:1 | 0.531435 | 0.03567 |
| PI 36:6\|PI 18:3_18:3 | -0.62265 | 0.0364 |
| PE 36:5 | -0.78835 | 0.036475 |
| Cer 48:9;4O\|Cer 17:3;2O/31:6;2O | 0.104111 | 0.036633 |
| PE 36:3 | -0.15534 | 0.036869 |
| PE O-36:5 | -0.18574 | 0.036898 |
| PC 31:1 | -0.0828 | 0.037161 |
| PS 36:4 | 0.330631 | 0.037252 |
| PC 32:2 | 0.166408 | 0.038746 |
| PE 36:3;2O\|PE 18:1_18:2;2O | -0.55712 | 0.039098 |
| PG 32:1\|PG 16:0_16:1 | 0.373701 | 0.039716 |
| PC O-33:1 | -0.04933 | 0.040204 |
| PE-Cer 30:1;2O | -0.3202 | 0.042091 |
| PC O-36:2\|PC O-18:0_18:2 | -0.24591 | 0.042713 |
| PI 36:4\|PI 18:2_18:2 | -0.25068 | 0.043083 |
| PC 31:0\|PC 15:0_16:0 | 0.476998 | 0.043236 |
| TG 41:0\|TG 12:0_14:0_15:0 | 0.910582 | 0.043248 |
| DG 50:7 | -0.20171 | 0.043818 |
| PE O-34:1\|PE O-18:0_16:1 | 0.16329 | 0.044499 |
| PC O-37:4 | 0.25072 | 0.046159 |
| ST 28:4;O align_ID:1213 | -0.57775 | 0.047351 |
| TG 48:4\|TG 14:1_16:1_18:2 | -0.5211 | 0.04741 |
| PG 36:2\|PG 18:0_18:2 | -0.29754 | 0.047901 |
| Cer 34:1;2O\|Cer 14:1;2O/20:0 | -0.2059 | 0.047937 |
| PE O-38:3 | -0.11107 | 0.048601 |

| | | |
|---|---|---|
| PE 38:3\|PE 20:0_18:3 | -0.17222 | 0.048621 |
| PE O-37:3\|PE O-19:0_18:3 | -0.24873 | 0.049009 |
| DG 28:4 | -0.6406 | 0.049059 |
| PE O-38:1\|PE O-20:0_18:1 | 0.161993 | 0.050874 |
| SM 42:9;2O | -0.6493 | 0.050875 |
| ST 28:3;O align_ID:1116 | 0.606589 | 0.051332 |
| PC O-35:0 | -0.26575 | 0.052232 |
| SM 61:6;2O | -0.29115 | 0.055158 |
| PC 35:1\|PC 18:0_17:1 | 0.488378 | 0.056544 |
| PE 35:1\|PE 17:0_18:1 | 0.139341 | 0.057285 |
| PC O-37:8 | -0.52704 | 0.057506 |
| PG 35:3 | -0.37047 | 0.057945 |
| PE-Cer 38:2;2O | -0.31552 | 0.059568 |
| Cer 46:9;4O\|Cer 15:2;2O/31:7;2O | 0.194787 | 0.060019 |
| TG 43:1\|TG 13:0_14:0_16:1 | 0.571646 | 0.060313 |
| PE 32:3 | -0.21982 | 0.060483 |
| PE-Cer 36:0;3O | 0.260335 | 0.060599 |
| CAR 18:0 | -0.32391 | 0.061186 |
| PC 35:1 | 0.616396 | 0.063128 |
| Dioctyl phthalate | -0.36325 | 0.063932 |
| CL 76:3\|CL 16:0_24:0_18:1_18:2 | -0.36744 | 0.063967 |
| DG 21:0 | 0.632317 | 0.065216 |
| NAE 22:4 align_ID:1089 | 0.461382 | 0.067822 |
| TG 38:0\|TG 12:0_12:0_14:0 | 0.552767 | 0.068222 |
| DG 50:5 | 0.15635 | 0.068877 |
| FA 10:0 | -0.18114 | 0.069769 |
| TG 39:1;1O\|TG 10:0_16:0_13:1;1O | -0.60216 | 0.070009 |
| FA 34:1 | -0.49688 | 0.0721 |
| TG 46:3\|TG 14:1_16:1_16:1 | -0.25422 | 0.073219 |
| TG 43:0\|TG 14:0_14:0_15:0 | 0.886077 | 0.074149 |
| DG 36:5\|DG 18:2_18:3 | -0.35767 | 0.07531 |
| PE 30:1 | -0.2131 | 0.075483 |
| PE O-35:3\|PE O-17:0_18:3 | -0.17983 | 0.077624 |
| TG 41:1\|TG 13:0_14:0_14:1 | 0.403638 | 0.078789 |
| CL 72:8\|CL 18:2_18:2_18:2_18:2 | -0.21797 | 0.078915 |
| FA 11:0 | -0.27678 | 0.078945 |
| SM 39:4;3O | 0.151314 | 0.079426 |
| FA 16:2 | -0.58421 | 0.080522 |
| CL 80:5\|CL 18:0_26:0_18:2_18:3 | -0.52196 | 0.080738 |
| CL 72:4\|CL 39:1_33:3 | -0.18614 | 0.08086 |
| Cer 36:2;2O\|Cer 14:2;2O/22:0 | -0.17949 | 0.081014 |
| PC 32:1\|PC 16:0_16:1 | 0.061044 | 0.081969 |
| PC O-36:1\|PC O-18:0_18:1 | 0.370546 | 0.082304 |
| PE 36:1\|PE 18:0_18:1 | 0.069378 | 0.08247 |
| NAE 22:4 align_ID:1090 | 0.447364 | 0.08413 |
| PI 36:3\|PI 18:1_18:2 | 0.277145 | 0.08442 |

| | | |
|---|---|---|
| PE-Cer 34:1;2O\|PE-Cer 14:1;2O/20:0 | -0.23582 | 0.085002 |
| PE-Cer 33:1;2O | -0.25753 | 0.086136 |
| PE 35:4\|PE 17:2_18:2 | -0.35421 | 0.086568 |
| Cer 49:11;4O\|Cer 13:2;2O/36:9;2O | -0.41151 | 0.086748 |
| DG 28:2\|DG 14:1_14:1 | -0.25705 | 0.087324 |
| PE-Cer 38:0;2O\|PE-Cer 14:0;2O/24:0 | -0.29978 | 0.087678 |
| FA 20:3 | -0.48 | 0.088683 |
| PE O-40:1\|PE O-22:0_18:1 | 0.254805 | 0.088698 |
| DG O-42:4\|DG O-22:3_20:1 | -0.50901 | 0.09204 |
| PC O-36:6 | -0.43528 | 0.092965 |
| PE 34:4 | -0.1329 | 0.094291 |
| FA 28:1 | -0.27733 | 0.098649 |
| ST 29:1;O align_ID:1448 | 0.922859 | 0.100865 |
| FA 13:0 | -0.42837 | 0.1011 |
| DG 49:13 | -0.4252 | 0.102441 |
| PE-Cer 34:0;2O | -0.23138 | 0.103257 |
| DG 36:3\|DG 18:1_18:2 | 0.604441 | 0.106438 |
| PE 36:3;O\|PE 18:2_18:1;O | -0.46931 | 0.106735 |
| PC 35:5 | 0.32254 | 0.107046 |
| CL 81:4\|CL 17:0_28:0_18:2_18:2 | 0.678278 | 0.108078 |
| PI 34:2 | 0.207604 | 0.108467 |
| TG 37:1;1O\|TG 10:0_14:0_13:1;1O | -0.4878 | 0.108565 |
| PE O-36:3 | -0.10941 | 0.108913 |
| PE 36:2\|PE 18:0_18:2 | -0.03379 | 0.111076 |
| PC 37:4 | -0.2938 | 0.111793 |
| PE 40:2\|PE 22:0_18:2 | -0.23434 | 0.113031 |
| FA 15:0 | -0.35214 | 0.114652 |
| PA 36:4\|PA 18:2_18:2 | -0.43952 | 0.115075 |
| TG 39:1\|TG 12:0_13:0_14:1 | 0.251574 | 0.116004 |
| FA 14:1 | -0.52634 | 0.117274 |
| PE O-40:5\|PE O-22:3_18:2 | -0.36558 | 0.117494 |
| FA 15:1 | -0.45972 | 0.118973 |
| CL 70:7\|CL 16:1_18:2_18:2_18:2 | 0.234507 | 0.120184 |
| FA 22:1 | -0.32601 | 0.120637 |
| Cer 38:0;2O\|Cer 14:0;2O/24:0 | -0.20284 | 0.121537 |
| TG 44:1\|TG 14:0_14:0_16:1 | 0.310617 | 0.122242 |
| FA 17:0 | -0.31297 | 0.124383 |
| TG 47:1\|TG 14:0_15:0_18:1 | 0.519338 | 0.125874 |
| PE-Cer 37:3;3O | -0.18081 | 0.128336 |
| PI 36:3 | 0.182408 | 0.129426 |
| PC O-32:3 | -0.16195 | 0.13033 |
| TG 45:1\|TG 14:0_15:0_16:1 | 0.552904 | 0.131068 |
| TG 52:5\|TG 16:1_18:2_18:2 | -0.34682 | 0.131283 |
| DG 41:5 | 0.215077 | 0.131905 |
| TG 53:3;O2\|TG 14:0_16:1_8:0;O(FA 15:1) | 0.464925 | 0.134415 |
| TG 55:3;O2\|TG 16:0_16:1_8:0;O(FA 15:1) | 0.417771 | 0.135241 |

| | | |
|---|---|---|
| FA 17:1 | -0.39343 | 0.136086 |
| TG 43:2;2O\|TG 16:0_14:1_13:1;2O | -0.46058 | 0.136783 |
| TG 51:1\|TG 16:0_17:0_18:1 | 0.550601 | 0.13789 |
| PE 27:1;O\|PE 10:0_17:1;O align_ID:1108 | -0.22088 | 0.138047 |
| TG 52:1\|TG 16:0_18:0_18:1 | 0.349436 | 0.138441 |
| PC 38:1\|PC 20:0_18:1 | 0.373311 | 0.138894 |
| PC 35:3 | 0.145637 | 0.140181 |
| TG 42:1\|TG 12:0_14:0_16:1 | 0.303721 | 0.143116 |
| TG 40:0\|TG 12:0_14:0_14:0 | 0.597653 | 0.143856 |
| PC 30:1\|PC 14:0_16:1 | 0.162321 | 0.145961 |
| PC O-35:1 | 0.203178 | 0.147898 |
| TG 48:0\|TG 14:0_16:0_18:0 | 0.28536 | 0.14885 |
| SM 39:8;3O | -0.63914 | 0.149279 |
| PC 36:3\|PC 18:1_18:2 | 0.043837 | 0.14949 |
| SE 28:1/18:2 | 0.218912 | 0.150391 |
| Cer 46:1;4O | 0.260569 | 0.151604 |
| PE O-30:0\|PE O-16:0_14:0 | -0.13634 | 0.151723 |
| PI 32:2 | 0.298525 | 0.152312 |
| TG 52:4\|TG 16:1_18:1_18:2 | -0.25876 | 0.152919 |
| PC O-41:3 | -0.58786 | 0.153455 |
| TG 53:1\|TG 17:0_18:0_18:1 | 0.500298 | 0.153883 |
| TG 45:0\|TG 14:0_15:0_16:0 | 0.531328 | 0.155797 |
| FA 30:1 | -0.35446 | 0.159106 |
| DG 48:11 | -0.12853 | 0.161403 |
| FA 20:2 | -0.4064 | 0.162761 |
| PC O-32:1 | -0.09309 | 0.164092 |
| FA 19:2 | -0.40567 | 0.164349 |
| CAR 18:2 | -0.41727 | 0.165304 |
| TG 45:2\|TG 13:0_16:1_16:1 | 0.321257 | 0.166481 |
| FA 32:2 | -0.32487 | 0.167194 |
| PE-Cer 37:1;2O | -0.24374 | 0.167256 |
| PC O-35:3 align_ID:3729 | -0.15671 | 0.167804 |
| PC 32:2\|PC 16:1_16:1 | 0.127168 | 0.16963 |
| DG 34:2\|DG 16:0_18:2 | 0.191817 | 0.170399 |
| TG 49:1\|TG 15:0_16:0_18:1 | 0.556071 | 0.171303 |
| PE 35:2\|PE 17:0_18:2 | -0.09274 | 0.172323 |
| PC O-33:2 align_ID:3468 | -0.21113 | 0.172507 |
| PE-Cer 32:0;2O\|PE-Cer 14:0;2O/18:0 | -0.1475 | 0.174577 |
| PC 40:6 | -0.1937 | 0.174622 |
| PE 35:0 | 0.182057 | 0.174847 |
| FA 18:2 align_ID:346 | 0.294826 | 0.175432 |
| TG 42:0\|TG 14:0_14:0_14:0 | 0.639212 | 0.176503 |
| CL 78:6\|CL 24:0_18:2_18:2_18:2 | -0.2247 | 0.179109 |
| PI 34:3\|PI 16:1_18:2 | 0.191527 | 0.179962 |
| PC 34:2\|PC 16:1_18:1 | -0.02414 | 0.180012 |
| PE-Cer 34:1;3O\|PE-Cer 18:1;2O/16:0;O | 0.264221 | 0.180252 |

| | | |
|---|---|---|
| PC 38:6 | 0.336597 | 0.192968 |
| PE-Cer 34:2;2O\|PE-Cer 14:2;2O/20:0 | -0.19114 | 0.194775 |
| DG 30:2\|DG 14:1_16:1 | -0.27102 | 0.200272 |
| PC O-39:9 | -0.50044 | 0.200591 |
| ST 29:1;O align_ID:1223 | 0.489932 | 0.202352 |
| PC O-38:10 | 0.425498 | 0.204632 |
| DG 33:6 | 1.205956 | 0.207503 |
| PC 40:2 | -0.15988 | 0.207759 |
| DG 34:1\|DG 16:0_18:1 | 0.156157 | 0.20971 |
| PE O-38:2\|PE O-20:0_18:2 | -0.06883 | 0.210872 |
| LPE 18:2 | -0.20656 | 0.210936 |
| PS 36:6\|PS 18:3_18:3 | 0.237152 | 0.211827 |
| PE O-34:3\|PE O-16:1_18:2 | -0.17612 | 0.212005 |
| PC O-39:7 | -0.33846 | 0.213304 |
| DG 26:4 | -0.6354 | 0.213842 |
| DG O-39:2\|DG O-19:1_20:1 | 0.762657 | 0.21437 |
| Cer 44:1;3O\|Cer 18:1;2O/26:0;O | 0.244355 | 0.215055 |
| FA 34:3 | -0.36788 | 0.215591 |
| FA 19:1 | -0.3289 | 0.215628 |
| TG 39:2;2O\|TG 14:0_15:2_10:0;2O | -0.5293 | 0.216686 |
| PC 36:2 align_ID:4298 | -0.10157 | 0.218608 |
| FA 26:1 | -0.20077 | 0.218636 |
| TG 50:0\|TG 16:0_16:0_18:0 | 0.197992 | 0.22148 |
| PS 36:6 | 0.188782 | 0.221952 |
| DG O-42:3\|DG O-15:3_27:0 | 0.151395 | 0.223841 |
| PG 34:1\|PG 16:0_18:1 | 0.164719 | 0.225826 |
| PC 33:0\|PC 16:0_17:0 | 0.178218 | 0.225863 |
| PI 36:4 | -0.12332 | 0.227453 |
| LPC 18:1/0:0 | 0.164127 | 0.228107 |
| PE 30:1\|PE 14:0_16:1 | -0.14988 | 0.228486 |
| DG 33:1\|DG 15:0_18:1 | 0.304566 | 0.22911 |
| MG 26:6 | -0.89537 | 0.229653 |
| LPC 18:2/0:0 | -0.28325 | 0.230734 |
| PC O-37:1 | 0.258131 | 0.233174 |
| TG 44:3\|TG 14:1_14:1_16:1 | -0.16397 | 0.234261 |
| TG 46:1\|TG 14:0_16:0_16:1 | 0.278043 | 0.23639 |
| DG 24:0 | -0.17431 | 0.242171 |
| PE O-38:0\|PE O-18:0_20:0 | -0.12342 | 0.242236 |
| PE 35:0\|PE 17:0_18:0 | 0.148116 | 0.244384 |
| SM 36:2;3O align_ID:3629 | 0.100909 | 0.245676 |
| TG 40:1\|TG 12:0_12:0_16:1 | 0.244453 | 0.246231 |
| PE 38:0\|PE 18:0_20:0 | -0.16526 | 0.247322 |
| NAE 4:0 | -0.23274 | 0.249288 |
| PE P-36:5\|PE P-18:3_18:2 | 0.105543 | 0.25004 |
| FA 32:1 | -0.35042 | 0.251737 |
| Cer 42:0;4O\|Cer 18:0;3O/24:0;(2OH) | -0.22325 | 0.255166 |

| | | |
|---|---|---|
| CL 78:5\|CL 18:0_24:0_18:2_18:3 | -0.32383 | 0.255273 |
| SM 37:1;2O\|SM 21:0;2O/16:1 | -0.21272 | 0.25534 |
| CAR 24:0 | -0.12526 | 0.256751 |
| TG 52:6\|TG 16:1_18:2_18:3 | -0.47121 | 0.258226 |
| PE-Cer 36:2;2O\|PE-Cer 14:2;2O/22:0 | -0.15857 | 0.258635 |
| FA 22:4 | -0.13992 | 0.259616 |
| FA 16:1 | -0.30904 | 0.26053 |
| PC 26:0 align_ID:2772 | 0.148866 | 0.260711 |
| DG 31:1\|DG 15:0_16:1 | 0.286077 | 0.261234 |
| ST 27:1;O;S | -0.26234 | 0.261273 |
| PE-Cer 32:1;3O | 0.269451 | 0.262133 |
| DG 46:10 | 0.078906 | 0.262157 |
| PC O-35:2 | -0.07278 | 0.262737 |
| ST 27:1;O | -0.26954 | 0.263262 |
| TG 43:2\|TG 13:0_14:1_16:1 | 0.254591 | 0.26725 |
| SE 29:1/18:1 | 0.374421 | 0.267784 |
| LPC 18:3/0:0 | -0.24285 | 0.268318 |
| FA 20:1 | -0.24971 | 0.268578 |
| DG 26:0\|DG 12:0_14:0 | 0.170021 | 0.268875 |
| PC 28:0 | -0.11659 | 0.269275 |
| PC O-32:0 | -0.08725 | 0.269488 |
| NAE 20:1 | 0.333412 | 0.269568 |
| PE 38:1\|PE 20:0_18:1 | 0.095899 | 0.270024 |
| PE O-37:2 | 0.10717 | 0.271422 |
| DG 52:10 | -0.28 | 0.274747 |
| TG 34:0\|TG 10:0_12:0_12:0 | 0.414491 | 0.275731 |
| PI 34:2\|PI 16:1_18:1 | 0.169965 | 0.276287 |
| PE O-37:2\|PE O-19:0_18:2 | -0.12533 | 0.27672 |
| SE 28:1/18:1 | 0.285453 | 0.277356 |
| PC 38:7 | -0.34546 | 0.277889 |
| PE O-36:2\|PE O-18:0_18:2 align_ID:1391 | -0.0507 | 0.282406 |
| NAE 20:2 | 0.386669 | 0.282567 |
| Cer 32:1;2O\|Cer 14:1;2O/18:0 | -0.13683 | 0.282961 |
| SE 28:2/18:1 | 0.201239 | 0.283954 |
| DG 28:1\|DG 14:0_14:1 | -0.17924 | 0.284315 |
| PC O-38:5 | 0.158924 | 0.285707 |
| ST 28:4;O align_ID:1098 | -0.56401 | 0.291265 |
| TG 44:0\|TG 14:0_14:0_16:0 | 0.461071 | 0.291309 |
| PC O-36:5 | -0.18999 | 0.291471 |
| TG 38:1\|TG 12:0_12:0_14:1 | 0.228983 | 0.292023 |
| TG 49:0\|TG 15:0_16:0_18:0 | 0.25692 | 0.2921 |
| TG 49:2\|TG 15:0_16:1_18:1 | 0.333252 | 0.292117 |
| DG 32:2\|DG 16:1_16:1 | -0.1973 | 0.296159 |
| Cer 48:5;4O\|Cer 30:4;3O(FA 18:0) | 2.029747 | 0.296176 |
| LPE-N (FA)33:2\|LPE-N (FA 15:0)18:2 | 0.131492 | 0.297528 |
| DG 30:1\|DG 14:0_16:1 | -0.20708 | 0.301114 |

| | | |
|---|---|---|
| TG 47:0\|TG 15:0_16:0_16:0 | 0.290925 | 0.302491 |
| DG 24:0\|DG 12:0_12:0 | -0.19424 | 0.304043 |
| PS 35:1\|PS 10:0_25:1 | -0.15156 | 0.306344 |
| PC 32:4\|PC 16:1_16:3 | -0.08563 | 0.313163 |
| SE 29:1/16:2 | 0.226665 | 0.313439 |
| NAE 22:5 | 0.449985 | 0.314275 |
| FA 9:0 align_ID:131 | 0.281254 | 0.315513 |
| ST 28:4;O align_ID:1100 | -0.20344 | 0.317099 |
| PE O-34:3\|PE O-16:0_18:3 | -0.03894 | 0.317237 |
| SM 36:2;3O align_ID:3627 | -0.35345 | 0.317653 |
| FA 32:0 | -0.21397 | 0.31815 |
| PC 38:6\|PC 16:0_22:6 | -0.1723 | 0.318311 |
| PC 36:0 | -0.10926 | 0.318743 |
| FA 42:0 | -0.2194 | 0.321987 |
| DG 36:2\|DG 18:0_18:2 | 0.227859 | 0.324893 |
| DG 34:4\|DG 16:1_18:3 | 0.288342 | 0.324969 |
| Cer 42:0;2O | -0.17078 | 0.325286 |
| MGDG 35:2 | -0.33068 | 0.325828 |
| PC 34:3 | 0.061294 | 0.327068 |
| PE-Cer 36:5;2O | -0.12304 | 0.328744 |
| TG 48:1\|TG 14:0_16:0_18:1 | 0.287466 | 0.329395 |
| PG 36:3\|PG 18:1_18:2 | -0.12068 | 0.331016 |
| PE-Cer 34:0;2O\|PE-Cer 14:0;2O/20:0 | -0.11272 | 0.335376 |
| TG 48:3\|TG 16:1_16:1_16:1 | -0.17125 | 0.335709 |
| DG O-35:2\|DG O-21:1_14:1 | 0.463612 | 0.335908 |
| FA 16:1;O | -0.27834 | 0.33683 |
| TG 51:2\|TG 17:0_16:1_18:1 | 0.339311 | 0.336972 |
| PC O-36:2 | -0.07008 | 0.337859 |
| PC O-39:2 | 0.139547 | 0.339388 |
| Cer 35:1;2O\|Cer 14:1;2O/21:0 | 0.258056 | 0.34166 |
| CAR 16:0 | -0.20282 | 0.342857 |
| DG 34:0\|DG 16:0_18:0 | 0.112588 | 0.344417 |
| Cer 28:0;4O | -1.93072 | 0.34662 |
| Cer 54:11;4O\|Cer 12:2;2O/42:9;2O | 0.11243 | 0.348645 |
| FA 18:1;O | -0.20627 | 0.348733 |
| PE 40:3 | -0.07027 | 0.352078 |
| SM 42:8;2O | -0.2941 | 0.352928 |
| ST 28:3;O align_ID:1222 | 0.106376 | 0.358249 |
| PC O-35:4 | 0.05164 | 0.364527 |
| FA 34:4 | -0.31978 | 0.366653 |
| TG 44:2\|TG 12:0_16:1_16:1 | 0.15167 | 0.367401 |
| DG 50:6 | -0.08712 | 0.368109 |
| FA 18:0 align_ID:367 | -0.22248 | 0.369894 |
| PC 36:0\|PC 18:0_18:0 | -0.21912 | 0.37116 |
| DG 33:0\|DG 16:0_17:0 | -0.15345 | 0.372907 |
| TG 53:2\|TG 17:0_18:1_18:1 | 0.319692 | 0.375351 |

| | | |
|---|---|---|
| PC O-37:5 | -0.04866 | 0.376216 |
| TG 50:1\|TG 16:0_16:0_18:1 | 0.235045 | 0.3763 |
| PE O-36:5\|PE O-18:3_18:2 | -0.1007 | 0.377862 |
| PE 36:1;2O\|PE 18:1_18:0;2O | 0.192388 | 0.379239 |
| DG O-33:1\|DG O-17:0_16:1 | 0.664973 | 0.385359 |
| Cer 42:1;3O\|Cer 18:1;2O/24:0;(2OH) | 0.161502 | 0.387781 |
| TG 54:5\|TG 18:1_18:2_18:2 | -0.20731 | 0.388517 |
| NAE 5:0 | -0.20502 | 0.388777 |
| PC O-33:2 align_ID:3465 | 0.25875 | 0.38929 |
| Cer 47:5;4O\|Cer 29:2;3O(FA 18:2) | Inf | 0.391002 |
| FA 24:1 | -0.23148 | 0.391239 |
| PE-Cer 38:5;2O | -0.15386 | 0.393247 |
| PE 27:1;O\|PE 10:0_17:1;O align_ID:1107 | -0.13001 | 0.393564 |
| PE-Cer 33:1;3O\|PE-Cer 14:0;2O/19:1;O | 0.32383 | 0.394658 |
| PE 34:2;3O\|PE 16:1_18:1;3O | -0.13936 | 0.394698 |
| NAE 14:0 | 5.978626 | 0.397616 |
| PE-Cer 36:0;2O\|PE-Cer 14:0;2O/22:0 | -0.0854 | 0.402916 |
| MG 37:1 | -0.61145 | 0.406898 |
| TG 46:0\|TG 14:0_16:0_16:0 | 0.262537 | 0.408151 |
| TG 54:0\|TG 16:0_18:0_20:0 | 0.520112 | 0.409939 |
| TG 54:6\|TG 18:2_18:2_18:2 | -0.32932 | 0.413317 |
| SM 42:7;2O | -0.23032 | 0.413518 |
| FA 39:0 | -0.16381 | 0.415902 |
| TG 62:1\|TG 14:0_16:0_32:1 | 0.219711 | 0.415982 |
| PC 35:4;O\|PC 18:2_17:2;O | -0.66804 | 0.419338 |
| PC 37:5 | 0.153826 | 0.419661 |
| PE-Cer 32:0;3O | -0.12403 | 0.419729 |
| PG 34:4 | -0.11781 | 0.419959 |
| PG 32:2\|PG 16:1_16:1 | -0.12847 | 0.420185 |
| CE 18:1 | -0.12509 | 0.4215 |
| DG 26:1\|DG 12:0_14:1 | -0.11232 | 0.421994 |
| LPE O-20:0 | -0.13088 | 0.422989 |
| LPC 18:2 | -0.16963 | 0.426126 |
| LPC 16:1 | 0.209031 | 0.427393 |
| DG 49:10 | -0.2823 | 0.428576 |
| NAE 22:6 | 0.923528 | 0.429265 |
| PE 32:1\|PE 16:0_16:1 | 0.063712 | 0.430936 |
| CL 71:5 | -0.04116 | 0.432109 |
| FA 30:0 | -0.22503 | 0.432202 |
| PC O-29:3 | -0.36479 | 0.437416 |
| SM 40:7;2O | -0.27115 | 0.440127 |
| DG 36:4\|DG 18:2_18:2 | 0.249036 | 0.442103 |
| FA 27:0 align_ID:639 | -0.20353 | 0.443222 |
| PG 36:1\|PG 18:0_18:1 | 0.143165 | 0.449804 |
| MG 36:3 | -0.13109 | 0.451767 |
| CL 80:3\|CL 18:0_26:0_18:1_18:2 | -0.27842 | 0.457787 |

| | | |
|---|---|---|
| PC O-39:1 | 0.464839 | 0.459595 |
| PE 34:4\|PE 16:1_18:3 | -0.07529 | 0.461331 |
| TG O-37:0\|TG O-13:0_12:0_12:0 | 0.148632 | 0.469758 |
| TG 58:3\|TG 16:1_18:1_24:1 | -0.26717 | 0.471616 |
| TG 50:1;2O\|TG 17:0_17:0_16:1;2O | 0.083194 | 0.472556 |
| PE 38:1 | 0.065356 | 0.47978 |
| LPE 16:1 | 0.191151 | 0.481973 |
| Cer 41:1;4O | 0.151643 | 0.485311 |
| TG 50:5\|TG 16:1_16:1_18:3 | 0.541454 | 0.485541 |
| PC 38:2\|PC 20:0_18:2 | -0.17286 | 0.487126 |
| CL 78:3\|CL 18:0_26:0_16:1_18:2 | -0.20257 | 0.48818 |
| FA 14:0 | -0.17842 | 0.489105 |
| PE 38:2\|PE 20:0_18:2 | -0.04669 | 0.489549 |
| DG 52:6 | 0.04531 | 0.490254 |
| DG 41:6 | -0.71057 | 0.493588 |
| LPC 16:0/0:0 | -0.06791 | 0.496975 |
| Cer 46:1;3O\|Cer 20:1;2O/26:0;O | 0.102055 | 0.497196 |
| SM 44:1;3O | 0.085131 | 0.497826 |
| DG 39:5 | 0.100872 | 0.498944 |
| TG 51:3\|TG 15:0_18:1_18:2 | 0.162807 | 0.502185 |
| PE 36:2\|PE 18:1_18:1 | -0.09289 | 0.50257 |
| PE 34:1\|PE 16:0_18:1 | 0.019324 | 0.502599 |
| DG 28:0\|DG 14:0_14:0 | -0.13373 | 0.503367 |
| PE-Cer 34:1;3O | -0.10663 | 0.50604 |
| Cer 42:1;2O\|Cer 18:1;2O/24:0 | 0.131656 | 0.508086 |
| FA 25:0 | -0.16474 | 0.514014 |
| ST 28:1;O | -0.99518 | 0.51418 |
| DG O-35:1\|DG O-19:0_16:1 | 0.322142 | 0.5148 |
| FA 36:0 | -0.1663 | 0.519365 |
| PE 27:0 | -0.08789 | 0.519461 |
| Cer 42:1;3O\|Cer 18:1;2O/24:0;O | 0.108821 | 0.52142 |
| FA 42:5 | 0.251418 | 0.521435 |
| PC O-36:9 | 0.300122 | 0.524822 |
| DG 40:5 | -0.16977 | 0.528672 |
| PC O-33:2 align_ID:3472 | -0.05182 | 0.53191 |
| ST 29:2;O | 0.316509 | 0.533272 |
| PE-Cer 36:1;2O\|PE-Cer 14:1;2O/22:0 | -0.08391 | 0.535573 |
| Cer 64:13;4O\|Cer 46:9;3O(FA 18:3) | -0.06341 | 0.539272 |
| DG 41:8 | -0.11118 | 0.542382 |
| PE O-36:4 | 0.074187 | 0.54448 |
| FA 19:0 | -0.13882 | 0.545685 |
| CL 74:7 | -0.09064 | 0.54619 |
| MG 18:0 | -0.10343 | 0.551077 |
| FA 27:0 align_ID:640 | -0.20646 | 0.553174 |
| PE-Cer 38:1;2O\|PE-Cer 14:1;2O/24:0 | -0.06886 | 0.561713 |
| PE 34:2 | -0.0561 | 0.561967 |

| | | |
|---|---|---|
| FA 9:0 align_ID:135 | -0.16306 | 0.562666 |
| DG 44:4 | 0.226609 | 0.563857 |
| TG 56:4\|TG 20:0_18:2_18:2 | -0.18355 | 0.565099 |
| PC 35:3\|PC 17:1_18:2 | 0.062447 | 0.566731 |
| TG 54:1\|TG 18:0_18:0_18:1 | 0.113648 | 0.57022 |
| TG 47:5;1O\|TG 14:1_18:2_15:2;1O | -0.03169 | 0.571038 |
| PC 33:3 | -0.05134 | 0.576906 |
| PE 40:1\|PE 22:0_18:1 | 0.0813 | 0.579469 |
| PC O-35:3 align_ID:3728 | -0.61433 | 0.579864 |
| SM 43:8;2O | -0.62761 | 0.579901 |
| FA 18:1 align_ID:353 | 0.294797 | 0.580498 |
| Cer 48:0;3O | 0.155924 | 0.581671 |
| PC 37:2 | 0.088906 | 0.582345 |
| PC 34:2 | 0.03647 | 0.583394 |
| Cer 40:1;4O | -0.18527 | 0.584123 |
| SM 40:6;3O | -0.60576 | 0.584243 |
| CL 72:7\|CL 18:1_18:2_18:2_18:2 | 0.081889 | 0.586736 |
| Cer 44:0;2O | -0.11406 | 0.58783 |
| PC 39:3 | -0.22997 | 0.588396 |
| PE 30:2;O\|PE 16:1_14:1;O | -0.07837 | 0.593151 |
| PC 33:1\|PC 15:0_18:1 | -0.01265 | 0.593538 |
| DG 31:0\|DG 15:0_16:0 | -0.11418 | 0.593751 |
| FA 18:0 align_ID:361 | 0.055257 | 0.605375 |
| Cer 43:1;4O | 0.087384 | 0.605439 |
| ST 27:2;O | -0.08616 | 0.607399 |
| Cer 34:1;4O | 0.069451 | 0.608288 |
| PE 40:2 | -0.06961 | 0.609863 |
| FA 34:0 | 0.089468 | 0.610236 |
| FA 24:0 align_ID:559 | -0.18291 | 0.614642 |
| PE O-34:2\|PE O-16:0_18:2 | -0.03089 | 0.615007 |
| SM 39:2;3O | 0.068505 | 0.619046 |
| PC 31:2;O\|PC 16:1_15:1;O | -0.07595 | 0.624108 |
| PC 30:3 | -0.09154 | 0.624952 |
| DGGA 32:0\|DGGA 16:0_16:0 | -0.0584 | 0.625591 |
| NAE 18:4 | -0.09106 | 0.638862 |
| FA 18:1 align_ID:352 | 0.095137 | 0.638963 |
| FA 28:2 | -0.06164 | 0.640232 |
| TG 50:3\|TG 16:1_16:1_18:1 | -0.08647 | 0.640519 |
| TG 58:2\|TG 16:0_18:1_24:1 | -0.16715 | 0.645516 |
| PE O-35:2 | -0.03143 | 0.647214 |
| TG 28:0\|TG 8:0_10:0_10:0 | 0.524133 | 0.64888 |
| Cer 36:1;2O\|Cer 14:1;2O/22:0 | -0.03613 | 0.650551 |
| PI 34:4\|PI 16:1_18:3 | 0.107847 | 0.652358 |
| FA 24:0 align_ID:557 | -0.06049 | 0.654984 |
| PC O-38:8 | 0.331736 | 0.656274 |
| DG 48:12 | -0.39155 | 0.659794 |

| | | |
|---|---|---|
| PC 30:2 | -0.05665 | 0.660463 |
| SM 43:1;3O | 0.052278 | 0.666597 |
| DG 29:0\|DG 14:0_15:0 | -0.07475 | 0.666667 |
| Cer 34:0;3O\|Cer 14:0;3O/20:0 | -0.04485 | 0.667193 |
| PE-Cer 34:0;3O | 0.068939 | 0.667551 |
| TG 54:3\|TG 18:0_18:1_18:2 | 0.12614 | 0.669922 |
| CL 82:9\|CL 28:0_18:3_18:3_18:3 | -0.14582 | 0.670505 |
| DG 34:3\|DG 16:1_18:2 | 0.054303 | 0.671705 |
| TG 57:8;1O\|TG 22:3_21:5_14:0;1O | 0.616065 | 0.672275 |
| PE 34:1 | 0.025455 | 0.676449 |
| TG 60:1\|TG 14:0_14:0_32:1 | 0.07354 | 0.679495 |
| PC 33:1\|PC 16:0_17:1 | 0.169382 | 0.683988 |
| LPC 18:0/0:0 | -0.05324 | 0.687223 |
| Cer 49:10;4O\|Cer 14:2;2O/35:8;2O | -0.03477 | 0.690976 |
| Cer 44:1;4O\|Cer 18:1;3O/26:0;(2OH) | 0.069352 | 0.694099 |
| PI 34:4 | -0.05774 | 0.700064 |
| TG 54:2\|TG 18:0_18:1_18:1 | 0.091299 | 0.705515 |
| MG 18:2 | -0.05689 | 0.708676 |
| Cer 42:1;4O | 0.079292 | 0.71318 |
| SM 39:6;3O | -0.34864 | 0.71966 |
| PC 38:1 align_ID:4354 | -0.25468 | 0.721933 |
| FA 16:0 align_ID:298 | -0.0648 | 0.724731 |
| DG 38:1\|DG 20:0_18:1 | -0.09637 | 0.727252 |
| TG 56:1\|TG 18:0_20:0_18:1 | 0.071239 | 0.727604 |
| TG 52:2\|TG 16:0_18:1_18:1 | 0.083856 | 0.731393 |
| DG 43:10 | -0.30422 | 0.732079 |
| FA 18:3 align_ID:341 | 0.107004 | 0.732087 |
| FA 21:0 align_ID:459 | -0.11605 | 0.737389 |
| Cer 46:0;3O\|Cer 24:0;2O/22:0;O | 0.064392 | 0.73944 |
| DG 30:3 | -0.19696 | 0.741874 |
| Diisodecyl phthalate (also known as the production of plastic) | 0.070906 | 0.742924 |
| DG 26:1 | -0.25453 | 0.743882 |
| PE O-37:5\|PE O-19:2_18:3 | -0.03871 | 0.745792 |
| PE O-36:2\|PE O-18:0_18:2 align_ID:1392 | -0.03767 | 0.746805 |
| TG 62:2\|TG 14:0_16:1_32:1 | -0.11692 | 0.748438 |
| FA 18:3 align_ID:342 | 0.068126 | 0.750298 |
| PS 35:2;O\|PS 18:1_17:1;O | -0.04574 | 0.751146 |
| CAR 18:1 | 0.083562 | 0.751612 |
| DG 20:0 | -0.20811 | 0.752266 |
| Cer 46:0;3O\|Cer 22:0;3O/24:0 | 0.086607 | 0.754762 |
| DG 42:9 | -0.31259 | 0.756915 |
| PE O-40:2\|PE O-22:0_18:2 | -0.0366 | 0.757021 |
| Cer 42:1;3O\|Cer 19:0;2O/23:1;O | 0.060206 | 0.760652 |
| PC O-38:9 | 0.172536 | 0.762495 |
| TG 54:1\|TG 12:0_14:0_28:1 | -0.05823 | 0.764275 |

| | | |
|---|---|---|
| TG 40:2\|TG 12:0_14:1_14:1 | 0.056759 | 0.765144 |
| PC 37:3 | 0.043097 | 0.769843 |
| Cer 36:0;3O\|Cer 14:0;3O/22:0 | -0.05374 | 0.771824 |
| FA 22:0 | 0.041894 | 0.771885 |
| TG 46:2\|TG 14:0_16:1_16:1 | 0.047055 | 0.772755 |
| Cer 34:0;2O\|Cer 14:0;2O/20:0 | 0.040763 | 0.777676 |
| DG 30:0 | 0.183209 | 0.778719 |
| DG 31:3 | -0.21887 | 0.781171 |
| PC O-39:4 | -0.10199 | 0.783675 |
| PA 36:3\|PA 18:1_18:2 | 0.072479 | 0.783792 |
| TG 56:2\|TG 14:0_18:1_24:1 | -0.0974 | 0.784319 |
| MG 21:1 | -0.04384 | 0.784453 |
| PE-Cer 37:2;3O\|PE-Cer 14:1;2O/23:1;O | -0.01941 | 0.785067 |
| DG 32:1\|DG 16:0_16:1 | -0.0537 | 0.785837 |
| Cer 34:1;3O\|Cer 18:1;2O/16:0;(2OH) | -0.05781 | 0.789606 |
| PI 34:3 | -0.03608 | 0.791199 |
| TG O-40:0\|TG O-14:0_12:0_14:0 | -0.12634 | 0.792435 |
| TG 26:0\|TG 8:0_8:0_10:0 | 0.288143 | 0.793922 |
| FAHFA 26:0;O\|FAHFA 16:0/10:0;O | 0.074142 | 0.803993 |
| NAE 20:0 | 0.200922 | 0.804234 |
| Cer 36:0;2O\|Cer 14:0;2O/22:0 | -0.03148 | 0.806059 |
| DG 36:5 | -0.20176 | 0.806907 |
| PE 36:2 | -0.02384 | 0.815203 |
| DG 35:5 | -0.15585 | 0.817017 |
| TG 24:0\|TG 8:0_8:0_8:0 | 0.230758 | 0.817939 |
| TG O-50:2\|TG O-18:0_16:1_16:1 | 0.106768 | 0.820052 |
| PC 35:2 | 0.014585 | 0.820281 |
| PE O-40:7\|PE O-22:5_18:2 | 0.037565 | 0.822548 |
| Cer 42:1;3O\|Cer 21:1;2O/21:0;O | -0.04787 | 0.826217 |
| Cer 44:1;3O\|Cer 26:0;2O/18:1;O | -0.04897 | 0.826674 |
| TG 54:7\|TG 18:2_18:2_18:3 | -0.13473 | 0.829215 |
| PC 40:7 | -0.03456 | 0.830151 |
| PE 35:3\|PE 17:1_18:2 | -0.02667 | 0.830357 |
| PC 38:5 | 0.027645 | 0.831897 |
| PC 34:4\|PC 16:1_18:3 | -0.02525 | 0.83402 |
| Norethisterone acetate | 0.051579 | 0.837858 |
| NAE 19:0 | -0.0825 | 0.83803 |
| PE O-34:2 | 0.024017 | 0.838062 |
| TG 42:2\|TG 12:0_14:1_16:1 | -0.03308 | 0.848386 |
| PC 36:2 align_ID:4091 | 0.056061 | 0.850544 |
| DG 32:0\|DG 16:0_16:0 | 0.034582 | 0.855289 |
| TG 54:4\|TG 18:1_18:1_18:2 | -0.04266 | 0.85828 |
| TG 64:2\|TG 16:0_16:1_32:1 | 0.083721 | 0.859237 |
| SM 63:7;2O | 0.027207 | 0.860244 |
| TG 45:2;1O\|TG 16:0_16:1_13:1;1O | -0.3099 | 0.860926 |
| TG 52:3\|TG 16:1_18:1_18:1 | 0.033093 | 0.861977 |

| | | |
|---|---|---|
| Cer 38:1;2O\|Cer 14:1;2O/24:0 | -0.01883 | 0.86249 |
| PC 34:2;O\|PC 18:1_16:1;O | 0.039027 | 0.870873 |
| PE O-35:2\|PE O-17:0_18:2 | -0.00906 | 0.873481 |
| Cer 44:0;3O\|Cer 20:0;3O/24:0 | 0.032961 | 0.874433 |
| MG 18:1 | 0.077917 | 0.875321 |
| PC 37:6 | -0.01456 | 0.878728 |
| PC O-34:8 | -0.10988 | 0.880701 |
| 4-Imidazoleacrylic acid | -0.07034 | 0.881023 |
| PC O-41:7 | -0.05771 | 0.882077 |
| PE O-37:0 | -0.01407 | 0.884643 |
| Cer 43:0;3O | -0.03634 | 0.884675 |
| TG 47:3\|TG 13:0_16:1_18:2 | -0.03272 | 0.886125 |
| CL 59:0 | -0.12376 | 0.887114 |
| PC O-36:3 | 0.010941 | 0.890799 |
| DG O-36:2\|DG O-18:0_18:2 | 0.022067 | 0.891633 |
| MG 38:1 | -0.02516 | 0.893032 |
| Cer 44:1;3O\|Cer 19:0;2O/25:1;O | 0.02712 | 0.893349 |
| PE-Cer 38:3;2O | -0.02516 | 0.894547 |
| TG 56:3\|TG 20:0_18:1_18:2 | 0.050444 | 0.897608 |
| PC O-34:2\|PC O-16:0_18:2 | -0.01905 | 0.897905 |
| TG 54:8\|TG 18:2_18:3_18:3 | -0.08904 | 0.904683 |
| FA 18:2 align_ID:347 | -0.02194 | 0.907206 |
| Cer 40:1;2O\|Cer 18:1;2O/22:0 | -0.01801 | 0.912321 |
| DG 18:0 | -0.07258 | 0.91248 |
| Cer 40:1;2O\|Cer 16:1;2O/24:0 | 0.015347 | 0.913325 |
| FA 20:0 | 0.023312 | 0.917993 |
| TG O-38:0\|TG O-14:0_12:0_12:0 | 0.021057 | 0.918571 |
| DG 37:5 | -0.01888 | 0.918906 |
| DG 42:10 | -0.07738 | 0.920618 |
| SE 29:1/18:2 | 0.027686 | 0.922842 |
| TG 50:2\|TG 16:0_16:1_18:1 | -0.02164 | 0.923198 |
| PC 34:3\|PC 16:1_18:2 | 0.004745 | 0.923357 |
| FA 16:0 align_ID:295 | -0.01318 | 0.924594 |
| TG 48:2\|TG 14:0_16:1_18:1 | -0.01695 | 0.927908 |
| LPC 16:0 | 0.01827 | 0.928583 |
| FA 21:0 align_ID:460 | 0.025741 | 0.93279 |
| PE-Cer 36:1;2O\|PE-Cer 18:0;2O/18:1 | 0.008798 | 0.935225 |
| PE P-36:6\|PE P-18:3_18:3 | -0.01296 | 0.935362 |
| TG 52:0\|TG 16:0_18:0_18:0 | 0.01504 | 0.936461 |
| PC O-37:2 | 0.009505 | 0.938982 |
| PE 37:3\|PE 19:1_18:2 | -0.01024 | 0.939146 |
| MG 16:0 | -0.01414 | 0.949955 |
| SHexCer 33:2;3O | 0.006696 | 0.955274 |
| LPE 18:1 align_ID:793 | 0.010248 | 0.957371 |
| PE O-34:1\|PE O-17:0_17:1 | 0.005469 | 0.961201 |
| FA 28:0 | -0.01149 | 0.965947 |

| | | |
|---|---|---|
| PC O-38:4 | 0.041173 | 0.965995 |
| DG 22:1 | -0.03358 | 0.966431 |
| PC 38:2 align_ID:4335 | 0.036041 | 0.966575 |
| FA 18:1 align_ID:355 | -0.00828 | 0.967056 |
| PC O-32:2 | -0.01252 | 0.967522 |
| FAHFA 30:0;O\|FAHFA 16:0/14:0;O | -0.01553 | 0.968233 |
| PC O-38:2 | -0.00349 | 0.970561 |
| LPE 18:1 align_ID:1741 | -0.00612 | 0.971283 |
| PG 34:3\|PG 16:1_18:2 | 0.002841 | 0.972565 |
| PG 32:0\|PG 16:0_16:0 | -0.00447 | 0.975765 |
| FA 29:0 | -0.00718 | 0.976934 |
| PC 36:1 | 0.010533 | 0.979 |
| DG 25:1 | -0.01102 | 0.979912 |
| FA 18:0 align_ID:365 | 0.005639 | 0.980541 |
| FA 12:0 | -0.00365 | 0.988109 |
| Dodecylbenzenesulfonic acid | 0.002278 | 0.993918 |
| FA 26:0 | -0.00179 | 0.994382 |
| ST 28:4;O align_ID:1101 | -0.00081 | 0.994452 |
| FA 23:0 | -0.00188 | 0.994913 |
| Cer 42:0;3O\|Cer 18:0;3O/24:0 | 0.000869 | 0.996869 |
| **Differentially Regulated Lipids between Male and Female brains** | | |
| **Lipid identity** | **log2FC (male/female)** | **p-value** |
| DG 24:0\|DG 12:0_12:0 | -1.18859 | 4.30E-09 |
| PC O-36:4 | -0.55223 | 6.57E-07 |
| DG 26:1\|DG 12:0_14:1 | -0.93187 | 6.81E-07 |
| TG 53:3;O2\|TG 14:0_16:1_8:0;O(FA 15:1) | 1.922688 | 1.57E-06 |
| PC 35:1\|PC 18:0_17:1 | -0.40994 | 4.19E-06 |
| TG 55:3;O2\|TG 16:0_16:1_8:0;O(FA 15:1) | 3.112587 | 4.27E-06 |
| PC 37:1 | -1.08442 | 1.44E-05 |
| TG 49:2;2O\|TG 14:0_14:0_21:2;2O | 2.770394 | 1.98E-05 |
| PC O-36:5 | -0.9363 | 1.98E-05 |
| DG 24:0 | -0.94535 | 3.51E-05 |
| CAR 18:2 | -0.76943 | 3.79E-05 |
| TG 58:2\|TG 16:0_18:1_24:1 | 1.388462 | 6.02E-05 |
| PE O-30:0\|PE O-16:0_14:0 | 0.228063 | 6.73E-05 |
| TG 62:2\|TG 14:0_16:1_32:1 | 1.314253 | 8.80E-05 |
| PE 34:2\|PE 16:0_18:2 | 0.092223 | 0.000102 |
| TG 56:2\|TG 14:0_18:1_24:1 | 1.202214 | 0.000111 |
| TG 36:0\|TG 12:0_12:0_12:0 | -1.17475 | 0.000144 |
| DG 26:0\|DG 12:0_14:0 | -0.57773 | 0.000217 |
| CAR 18:1 | -0.58529 | 0.000248 |
| TG 58:3\|TG 16:1_18:1_24:1 | 1.093297 | 0.000288 |
| LPE 16:1 | 0.791035 | 0.00036 |
| PC 38:1\|PC 20:0_18:1 | -0.39073 | 0.000394 |

| | | |
|---|---|---|
| PC 37:2 | -0.55336 | 0.000422 |
| DG 39:5 | -0.17947 | 0.000453 |
| DG 49:10 | -1.95317 | 0.000497 |
| PC O-32:0 | -0.27181 | 0.000683 |
| PE O-34:2\|PE O-18:1_16:1 | 0.346978 | 0.000842 |
| DG 21:0 | -0.93912 | 0.000936 |
| PE O-34:3\|PE O-16:1_18:2 | 0.499019 | 0.001044 |
| DG 40:5 | -2.05807 | 0.001073 |
| PC 33:0\|PC 16:0_17:0 | -0.37324 | 0.001095 |
| PE 32:1\|PE 16:0_16:1 | 0.264621 | 0.001103 |
| PE 35:0 | -0.33865 | 0.00118 |
| TG 64:2\|TG 16:0_16:1_32:1 | 0.852538 | 0.001243 |
| TG 56:3\|TG 20:0_18:1_18:2 | 0.686801 | 0.001311 |
| PC 35:3 | -0.33766 | 0.00141 |
| Cer 34:2;2O\|Cer 14:2;2O/20:0 | -0.26556 | 0.001461 |
| LPC 16:1 | 0.623378 | 0.001548 |
| PC 31:0\|PC 15:0_16:0 | -0.39182 | 0.001659 |
| CAR 24:1 | 2.01551 | 0.001699 |
| TG 37:0\|TG 12:0_12:0_13:0 | -0.79193 | 0.001828 |
| PE-Cer 32:2;2O\|PE-Cer 14:2;2O/18:0 | 0.348119 | 0.001977 |
| PE-Cer 30:1;2O | 0.454438 | 0.002155 |
| PC O-45:11 | 0.960753 | 0.002159 |
| TG 38:0\|TG 12:0_12:0_14:0 | -0.76091 | 0.002258 |
| PC 33:1 | -0.22851 | 0.002362 |
| PE 34:1\|PE 16:0_18:1 | 0.079313 | 0.00242 |
| CAR 22:0 | -0.18681 | 0.002442 |
| PC 35:2\|PC 17:0_18:2 | -0.30537 | 0.002648 |
| TG 48:4\|TG 14:1_16:1_18:2 | 0.395718 | 0.002796 |
| TG 39:2;2O\|TG 14:0_15:2_10:0;2O | -1.10134 | 0.003023 |
| PC 34:3 | -0.13978 | 0.003453 |
| PC O-38:5 | -0.25042 | 0.003529 |
| PC 33:0 | -0.35336 | 0.003587 |
| DG 41:8 | -0.71052 | 0.003877 |
| PC 31:1 | 0.069539 | 0.004259 |
| TG 50:4\|TG 16:1_16:1_18:2 | 0.311155 | 0.004344 |
| PE-Cer 32:1;2O\|PE-Cer 14:1;2O/18:0 | 0.318782 | 0.004585 |
| DG 28:2\|DG 14:1_14:1 | -0.58895 | 0.005288 |
| DG 27:0 | -0.8999 | 0.005356 |
| PC 35:1\|PC 17:0_18:1 | -0.42042 | 0.005359 |
| PC 38:1 align_ID:4355 | -0.36808 | 0.005424 |
| PE O-36:3\|PE O-18:1_18:2 | 0.137998 | 0.005701 |
| PC 34:0 | -0.48687 | 0.005783 |
| Cer 38:0;2O\|Cer 14:0;2O/24:0 | 0.242338 | 0.006138 |
| PC 32:4\|PC 16:1_16:3 | 0.298607 | 0.006832 |
| DG 28:0\|DG 14:0_14:0 | -0.40787 | 0.006985 |
| PE 38:1\|PE 20:0_18:1 | -0.23859 | 0.007521 |

| | | |
|---|---|---|
| DG 32:5 | 0.631581 | 0.007546 |
| PC O-36:1 | -0.34846 | 0.007584 |
| TG 43:2;2O|TG 16:0_14:1_13:1;2O | -0.88539 | 0.007754 |
| TG 52:2|TG 16:0_18:1_18:1 | 0.331766 | 0.007808 |
| FA 42:0 | 0.420889 | 0.007999 |
| PE 38:0|PE 18:0_20:0 | -0.32669 | 0.008443 |
| PE O-37:0 | -0.28875 | 0.008716 |
| Cer 49:10;4O|Cer 14:2;2O/35:8;2O | -0.22123 | 0.00907 |
| FA 16:1;O | -1.02302 | 0.009195 |
| Cer 36:2;2O|Cer 14:2;2O/22:0 | -0.23763 | 0.009425 |
| TG 52:6|TG 16:1_18:2_18:3 | 0.706488 | 0.009943 |
| PC O-36:6 | -0.5417 | 0.010419 |
| LPE 18:1 align_ID:1741 | 0.446969 | 0.011034 |
| PE 35:0|PE 17:0_18:0 | -0.23832 | 0.011089 |
| TG 50:3|TG 16:1_16:1_18:1 | 0.24473 | 0.011545 |
| PE O-36:5 | -0.25272 | 0.011602 |
| PE 30:1|PE 14:0_16:1 | 0.326195 | 0.011638 |
| CL 71:5 | 0.114008 | 0.012068 |
| TG 50:2|TG 16:0_16:1_18:1 | 0.365298 | 0.012808 |
| FA 28:2 | -0.70235 | 0.012886 |
| TG 46:3|TG 14:1_16:1_16:1 | 0.311829 | 0.013175 |
| PE 40:3 | -0.3676 | 0.013226 |
| CAR 16:0 | -0.39755 | 0.01369 |
| LPC 18:2/0:0 | 0.506782 | 0.013705 |
| LPC 18:2 | 0.483198 | 0.014412 |
| PE P-34:1|PE P-18:0_16:1 | 0.241008 | 0.014526 |
| Cer 48:10;4O|Cer 12:2;2O/36:8;2O | 0.0431 | 0.01501 |
| LPE 18:2 | 0.429711 | 0.015283 |
| DG 28:1|DG 14:0_14:1 | -0.40058 | 0.015313 |
| PC O-36:1|PC O-18:0_18:1 | -0.27746 | 0.015938 |
| TG 38:1|TG 12:0_12:0_14:1 | -0.62957 | 0.016344 |
| PC 36:1|PC 18:0_18:1 | -0.12127 | 0.016955 |
| PE 37:1 | -0.24362 | 0.016976 |
| PC O-34:1|PC O-18:0_16:1 | -0.2596 | 0.018618 |
| TG 40:0|TG 12:0_14:0_14:0 | -0.50302 | 0.018866 |
| PE O-36:4|PE O-18:1_18:3 | 0.154847 | 0.019432 |
| FA 39:0 | 0.311666 | 0.019516 |
| PG 32:2|PG 16:1_16:1 | 0.562104 | 0.01952 |
| DG 44:4 | -0.85942 | 0.019753 |
| DG 28:2 | -1.13554 | 0.020226 |
| TG 52:0|TG 16:0_18:0_18:0 | 0.444531 | 0.020339 |
| TG 48:3|TG 16:1_16:1_16:1 | 0.312761 | 0.020845 |
| PE O-35:2 | 0.1449 | 0.021324 |
| FA 18:1 align_ID:353 | 1.11361 | 0.021415 |
| DG 46:10 | -0.14109 | 0.022514 |
| PE O-34:2 | 0.178373 | 0.022635 |

| | | |
|---|---|---|
| LPE 18:1 align_ID:793 | 0.426885 | 0.024136 |
| DG 38:1\|DG 20:0_18:1 | -0.65263 | 0.024585 |
| SM 31:6;2O | -0.52322 | 0.024803 |
| PC O-41:3 | 0.559198 | 0.026519 |
| PE O-32:0\|PE O-18:0_14:0 | 0.156917 | 0.027683 |
| PC 35:2 | -0.16407 | 0.030562 |
| MG 18:0 | -0.26916 | 0.030577 |
| DG 41:5 | -0.12058 | 0.032119 |
| PC 38:2 align_ID:4336 | -0.17713 | 0.032511 |
| Norethisterone acetate | -0.72043 | 0.0326 |
| PE O-37:5\|PE O-19:2_18:3 | 0.294454 | 0.03304 |
| PE-Cer 34:1;2O\|PE-Cer 14:1;2O/20:0 | 0.131836 | 0.033959 |
| FA 18:1;O | -0.69323 | 0.034526 |
| PE O-32:1 | 0.161078 | 0.035311 |
| PG 32:0\|PG 16:0_16:0 | 0.273442 | 0.035737 |
| TG 39:0\|TG 12:0_13:0_14:0 | -0.57903 | 0.035766 |
| PE 35:1;2O\|PE 18:0_17:1;2O | -0.92194 | 0.036263 |
| TG 54:2\|TG 18:0_18:1_18:1 | 0.482352 | 0.036289 |
| PE 37:3\|PE 19:1_18:2 | -0.24502 | 0.036349 |
| DG 32:2\|DG 16:1_16:1 | -0.56031 | 0.036791 |
| TG 54:1\|TG 12:0_14:0_28:1 | 0.30715 | 0.039181 |
| PE 36:3 | 0.161495 | 0.040122 |
| PC 26:0 align_ID:2772 | -0.38704 | 0.040666 |
| PC 28:0 | 0.17353 | 0.041189 |
| TG 46:4\|TG 12:0_16:1_18:3 | 0.308598 | 0.041264 |
| CAR 20:0 | -0.22805 | 0.041448 |
| PE 32:3\|PE 14:0_18:3 | 0.269309 | 0.041639 |
| PC 35:3\|PC 17:1_18:2 | -0.23496 | 0.04191 |
| TG 56:4\|TG 20:0_18:2_18:2 | 0.392096 | 0.041971 |
| TG 48:2\|TG 14:0_16:1_18:1 | 0.276562 | 0.042447 |
| TG 50:1;2O\|TG 17:0_17:0_16:1;2O | 0.211157 | 0.04246 |
| Cer 36:0;2O\|Cer 14:0;2O/22:0 | 0.189154 | 0.042737 |
| PG 34:2\|PG 16:0_18:2 | 0.204235 | 0.042909 |
| PC O-33:1 | 0.04991 | 0.043003 |
| PS 36:2 align_ID:4113 | -0.67428 | 0.043494 |
| PE 34:1 | 0.125524 | 0.043755 |
| Cer 46:9;4O\|Cer 15:2;2O/31:7;2O | 0.105956 | 0.044089 |
| PE 32:3 | 0.251833 | 0.044845 |
| PS 36:1\|PS 18:0_18:1 | -0.83091 | 0.045051 |
| PA 36:4\|PA 18:2_18:2 | 0.419445 | 0.045068 |
| TG 52:3\|TG 16:1_18:1_18:1 | 0.231269 | 0.045939 |
| PC 37:5 | -0.39802 | 0.046057 |
| CAR 24:0 | -0.31421 | 0.046303 |
| DG 52:6 | -0.0921 | 0.046966 |
| PC 26:0 align_ID:2773 | -0.69558 | 0.046983 |
| TG O-37:0\|TG O-13:0_12:0_12:0 | -0.53397 | 0.047502 |

| | | |
|---|---|---|
| CL 72:4\|CL 39:1_33:3 | 0.312387 | 0.048711 |
| TG 56:1\|TG 18:0_20:0_18:1 | 0.277113 | 0.049215 |
| FA 16:0 align_ID:298 | -0.24488 | 0.049639 |
| PC 35:1 | -0.29725 | 0.050359 |
| LPC 18:1 | 0.275292 | 0.051729 |
| SM 44:1;3O | -0.17825 | 0.051816 |
| PG 32:1\|PG 16:0_16:1 | 0.21891 | 0.052107 |
| MG 18:1 | 0.74724 | 0.052757 |
| PC O-33:2 align_ID:3472 | 0.131738 | 0.056247 |
| PS 34:1\|PS 16:0_18:1 | -0.71882 | 0.056557 |
| PC 32:0 | -0.36227 | 0.057495 |
| DG 29:0\|DG 14:0_15:0 | -0.51898 | 0.058105 |
| PC O-34:1 | -0.24718 | 0.058657 |
| DG 20:0 | -1.11355 | 0.058699 |
| PC 36:0 | -0.20653 | 0.059231 |
| FAHFA 26:0;O\|FAHFA 16:0/10:0;O | -0.62853 | 0.059539 |
| DG 32:0\|DG 16:0_16:0 | -0.3931 | 0.05955 |
| PE 36:3;O\|PE 18:2_18:1;O | 0.4214 | 0.059582 |
| DG 18:0 | -1.071 | 0.05989 |
| PE 34:2;2O\|PE 16:0_18:2;2O | 0.461625 | 0.06002 |
| DG 31:0\|DG 15:0_16:0 | -0.54312 | 0.060959 |
| TG 60:1\|TG 14:0_14:0_32:1 | 0.402787 | 0.061162 |
| Cer 47:9;4O\|Cer 15:2;2O/32:7;2O | -0.20031 | 0.061915 |
| FA 14:1 | -1.39501 | 0.062188 |
| FA 16:2 | -1.44011 | 0.062453 |
| NAE 20:1 | -1.03261 | 0.062539 |
| DG O-39:2\|DG O-19:1_20:1 | -3.12281 | 0.062743 |
| PC 37:6 | -0.28866 | 0.06353 |
| TG 42:0\|TG 14:0_14:0_14:0 | -0.47562 | 0.064164 |
| PC O-34:2\|PC O-16:0_18:2 | -0.17403 | 0.064541 |
| FA 20:0 | -0.59335 | 0.066891 |
| PC 40:2 | -0.21852 | 0.067441 |
| FA 16:0 align_ID:295 | -0.2193 | 0.067711 |
| NAE 22:4 align_ID:1089 | -0.884 | 0.068119 |
| PE 27:1;O\|PE 10:0_17:1;O align_ID:1107 | -0.31135 | 0.068404 |
| Cer 48:9;4O\|Cer 17:3;2O/31:6;2O | 0.067918 | 0.068958 |
| DG 44:7 | -0.15945 | 0.069154 |
| PE O-36:2\|PE O-18:0_18:2 align_ID:1391 | 0.056722 | 0.069436 |
| FA 18:0 align_ID:365 | -0.34784 | 0.069697 |
| PS 36:2 align_ID:4110 | -0.5248 | 0.070207 |
| PC O-35:3 align_ID:3728 | 2.980159 | 0.070564 |
| PS 36:2\|PS 18:1_18:1 | -0.69899 | 0.07124 |
| TG 52:5\|TG 16:1_18:2_18:2 | 0.393068 | 0.071488 |
| PE 34:3\|PE 16:1_18:2 | 0.122262 | 0.071743 |
| Cer 48:5;4O\|Cer 30:4;3O(FA 18:0) | -6.18129 | 0.072273 |
| CL 70:6\|CL 16:1_18:1_18:2_18:2 | -0.20369 | 0.072368 |

| | | |
|---|---|---|
| PE 36:1;O\|PE 18:0_18:1;O | 0.95934 | 0.072687 |
| Cer 40:1;4O | -2.8411 | 0.072942 |
| DG 33:0\|DG 16:0_17:0 | -0.47388 | 0.073399 |
| FA 15:1 | -1.17829 | 0.07492 |
| CE 18:1 | -0.38362 | 0.075292 |
| PG 34:1\|PG 16:0_18:1 | 0.243892 | 0.075419 |
| SM 40:7;2O | 0.452311 | 0.07594 |
| FA 30:2 | -0.28709 | 0.076304 |
| PG 36:2\|PG 18:1_18:1 | 0.258073 | 0.076389 |
| TG 54:1\|TG 18:0_18:0_18:1 | 0.402586 | 0.077415 |
| CL 71:7 | 0.125616 | 0.078486 |
| SE 28:2/18:1 | 0.240221 | 0.079293 |
| TG 34:0\|TG 10:0_12:0_12:0 | -0.90802 | 0.079366 |
| FA 36:0 | 0.320995 | 0.080663 |
| PI 32:1\|PI 16:0_16:1 | -0.26504 | 0.080682 |
| NAE 14:0 | Inf | 0.081596 |
| TG 54:3\|TG 18:0_18:1_18:2 | 0.363384 | 0.082032 |
| FA 30:0 | 0.333122 | 0.082264 |
| PI 36:1\|PI 18:0_18:1 | -0.38954 | 0.083129 |
| PC O-36:2\|PC O-18:0_18:2 | -0.11922 | 0.084414 |
| FA 24:0 align_ID:559 | 0.395624 | 0.084451 |
| SM 42:9;2O | 0.454832 | 0.085281 |
| MG 16:0 | -0.42271 | 0.085392 |
| PE 38:1 | -0.10647 | 0.085529 |
| NAE 20:0 | 1.832514 | 0.085759 |
| DG O-35:2\|DG O-21:1_14:1 | -3.10192 | 0.086107 |
| FA 28:1 | -0.36782 | 0.086397 |
| PE 36:4 | -0.11821 | 0.086728 |
| TG 41:0\|TG 12:0_14:0_15:0 | -0.48549 | 0.08772 |
| FA 16:1 | -0.82566 | 0.090258 |
| PS 36:4 | -0.22905 | 0.090725 |
| PE 38:2\|PE 20:0_18:2 | -0.10543 | 0.092027 |
| PC 33:3 | -0.1348 | 0.092373 |
| PE P-36:5\|PE P-18:3_18:2 | -0.13393 | 0.092782 |
| PE 36:4\|PE 18:2_18:2 | 0.059482 | 0.093903 |
| Cer 36:0;3O\|Cer 14:0;3O/22:0 | 0.153296 | 0.095719 |
| TG 40:1\|TG 12:0_12:0_16:1 | -0.36789 | 0.095853 |
| Cer 42:0;4O\|Cer 18:0;3O/24:0;(2OH) | 0.353706 | 0.097261 |
| LPE 16:0 | 0.18987 | 0.097651 |
| NAE 5:0 | -0.14972 | 0.097694 |
| PS 36:5 | -0.21614 | 0.098525 |
| FA 13:0 | -0.93446 | 0.099381 |
| PS 36:6\|PS 18:3_18:3 | -0.25681 | 0.100968 |
| PE-Cer 34:0;3O | -0.1334 | 0.101227 |
| PI 36:4 | 0.111177 | 0.101547 |
| PC 38:2 align_ID:4335 | 1.472318 | 0.101786 |

| | | |
|---|---|---|
| TG 54:4\|TG 18:1_18:1_18:2 | 0.293265 | 0.102065 |
| TG 52:4\|TG 16:1_18:1_18:2 | 0.281673 | 0.102667 |
| FA 27:0 align_ID:640 | 0.344194 | 0.103656 |
| DG 32:2 | -1.25766 | 0.104161 |
| PC 31:2;O\|PC 16:1_15:1;O | -0.2274 | 0.104595 |
| PC 30:1\|PC 14:0_16:1 | 0.164407 | 0.106545 |
| TG 51:1\|TG 16:0_17:0_18:1 | 0.282117 | 0.106707 |
| PC 39:3 | 0.699102 | 0.10747 |
| PE-Cer 34:0;2O | 0.090469 | 0.108178 |
| PS 36:3 | -0.38925 | 0.108261 |
| FA 14:0 | -0.68061 | 0.108745 |
| FA 34:1 | 0.445732 | 0.110168 |
| PS 34:2 | -0.44843 | 0.112231 |
| Diisodecyl phthalate | -0.46536 | 0.1123 |
| PE 34:0\|PE 16:0_18:0 | -0.07781 | 0.112851 |
| PE 33:1\|PE 16:0_17:1 | 0.155346 | 0.114198 |
| LPC 18:3/0:0 | 0.25664 | 0.114291 |
| FA 12:0 | -0.58254 | 0.116303 |
| LPC 16:0 | 0.201121 | 0.116604 |
| FA 15:0 | -0.58545 | 0.116695 |
| PE-Cer 37:2;3O\|PE-Cer 14:1;2O/23:1;O | 0.102207 | 0.117943 |
| DG O-42:3\|DG O-15:3_27:0 | 0.260768 | 0.119334 |
| SM 42:8;2O | 0.472346 | 0.119567 |
| TG 51:2\|TG 17:0_16:1_18:1 | 0.217592 | 0.121307 |
| CL 59:0 | 2.188376 | 0.121658 |
| Dodecylbenzenesulfonic acid | -0.42308 | 0.121894 |
| PE O-34:2\|PE O-16:0_18:2 | 0.082241 | 0.122657 |
| SE 29:1/18:1 | -0.24153 | 0.123496 |
| TG 54:5\|TG 18:1_18:2_18:2 | 0.288372 | 0.123805 |
| PE 34:4 | 0.272742 | 0.12419 |
| Cer 47:5;4O\|Cer 29:2;3O(FA 18:2) | #NAME? | 0.124744 |
| DG 41:6 | 2.96017 | 0.124993 |
| PE 36:5\|PE 18:2_18:3 | 0.084148 | 0.12513 |
| PC 38:5 | -0.26026 | 0.125385 |
| FA 18:3 align_ID:342 | -0.30893 | 0.12542 |
| PS 36:3\|PS 10:0_26:3 | -0.30678 | 0.125556 |
| FA 20:3 | -0.77956 | 0.126252 |
| DG O-42:4\|DG O-22:3_20:1 | 0.541276 | 0.127061 |
| DG 36:5 | 1.696843 | 0.127755 |
| PE 36:3;2O\|PE 18:1_18:2;2O | 0.362696 | 0.127889 |
| LPE-N (FA)33:2\|LPE-N (FA 15:0)18:2 | 0.121067 | 0.128503 |
| DG 43:10 | 1.919615 | 0.12885 |
| TG 42:1\|TG 12:0_14:0_16:1 | -0.21527 | 0.130547 |
| PC 34:2 | -0.08386 | 0.13068 |
| DG 31:3 | 1.594281 | 0.131696 |
| DG 48:12 | 1.851849 | 0.133372 |

| | | |
|---|---|---|
| FA 22:4 | -0.38799 | 0.134002 |
| PE O-40:1\|PE O-22:0_18:1 | -0.15418 | 0.135146 |
| PE-Cer 36:0;2O\|PE-Cer 14:0;2O/22:0 | 0.101712 | 0.135209 |
| TG 54:6\|TG 18:2_18:2_18:2 | 0.288456 | 0.136437 |
| Cer 49:11;4O\|Cer 13:2;2O/36:9;2O | -0.21098 | 0.138053 |
| PE 36:4;2O\|PE 18:2_18:2;2O | 0.339799 | 0.138388 |
| FA 21:0 align_ID:459 | 0.330267 | 0.138984 |
| DG 26:1 | 1.522673 | 0.139387 |
| PC 30:2 | 0.134248 | 0.139432 |
| FA 42:5 | -0.51076 | 0.140173 |
| MG 21:1 | -0.34635 | 0.140334 |
| TG 47:3\|TG 13:0_16:1_18:2 | 0.196922 | 0.140684 |
| TG 44:3\|TG 14:1_14:1_16:1 | 0.239813 | 0.140694 |
| DG 30:1\|DG 14:0_16:1 | -0.25241 | 0.141803 |
| SM 39:6;3O | 2.0876 | 0.142337 |
| TG 51:3\|TG 15:0_18:1_18:2 | 0.138768 | 0.143198 |
| TG 54:0\|TG 16:0_18:0_20:0 | 0.187642 | 0.143575 |
| DG 42:9 | 2.281153 | 0.144293 |
| PS 34:2\|PS 16:0_18:2 | -0.35589 | 0.145573 |
| TG 53:2\|TG 17:0_18:1_18:1 | 0.219731 | 0.14588 |
| PC 34:1\|PC 16:0_18:1 | -0.01386 | 0.146565 |
| PE 40:1\|PE 22:0_18:1 | -0.18251 | 0.146885 |
| PE O-36:1\|PE O-18:0_18:1 | 0.06173 | 0.147231 |
| CL 81:4\|CL 17:0_28:0_18:2_18:2 | -0.30337 | 0.147398 |
| FA 17:0 | -0.43068 | 0.148137 |
| PC O-37:8 | 0.36639 | 0.150529 |
| PC 38:1 align_ID:4354 | 1.274983 | 0.150618 |
| PC O-39:4 | 0.576404 | 0.150702 |
| PI 34:1 | -0.25277 | 0.151342 |
| CL 78:3\|CL 18:0_26:0_16:1_18:2 | 0.345458 | 0.155291 |
| TG 49:2\|TG 15:0_16:1_18:1 | 0.239631 | 0.155482 |
| PC 36:2 align_ID:4091 | -0.2636 | 0.156337 |
| NAE 22:4 align_ID:1090 | -0.57689 | 0.157083 |
| HexCer 34:2;2O | -0.13689 | 0.159123 |
| SM 40:6;3O | 2.60155 | 0.15928 |
| DG 50:5 | -0.11012 | 0.160772 |
| Cer 34:0;2O\|Cer 14:0;2O/20:0 | 0.100774 | 0.160821 |
| SM 43:8;2O | 2.523383 | 0.161063 |
| PI 32:1 | -0.18071 | 0.163516 |
| PC 30:0\|PC 14:0_16:0 | 0.056079 | 0.164412 |
| ST 29:1;O align_ID:1223 | -0.2375 | 0.164766 |
| PE 38:3\|PE 20:0_18:3 | -0.11414 | 0.165538 |
| FA 17:1 | -0.65718 | 0.165684 |
| DG 30:2\|DG 14:1_16:1 | -0.34129 | 0.165852 |
| PS 36:6 | -0.17879 | 0.166712 |
| TG 44:1\|TG 14:0_14:0_16:1 | -0.1615 | 0.168347 |

| | | |
|---|---|---|
| PC O-36:2 | -0.09558 | 0.169489 |
| PG 36:2\|PG 18:0_18:2 | 0.149592 | 0.171086 |
| PC 40:6 | -0.19193 | 0.171454 |
| DG 34:3\|DG 16:1_18:2 | -0.13772 | 0.171619 |
| PE O-34:1\|PE O-18:0_16:1 | 0.058565 | 0.172015 |
| DG 31:1\|DG 15:0_16:1 | -0.34357 | 0.172577 |
| PE O-32:1\|PE O-16:0_16:1 | 0.109991 | 0.174895 |
| PC 36:6\|PC 18:3_18:3 | -0.18623 | 0.174901 |
| PE-Cer 34:0;2O\|PE-Cer 14:0;2O/20:0 | 0.12634 | 0.175579 |
| DG 34:4\|DG 16:1_18:3 | -0.17327 | 0.176626 |
| SE 28:1/18:1 | -0.23561 | 0.177208 |
| FA 9:0 align_ID:135 | 0.415636 | 0.17929 |
| PE-Cer 32:0;2O\|PE-Cer 14:0;2O/18:0 | 0.160645 | 0.180353 |
| TG 41:2;1O\|TG 10:0_16:0_15:2;1O | 0.306972 | 0.184076 |
| PC O-37:3 | 0.092099 | 0.184523 |
| NAE 20:2 | -0.6151 | 0.185833 |
| PS 36:3\|PS 18:1_18:2 | -0.28138 | 0.189153 |
| PC O-29:3 | 0.559813 | 0.189501 |
| TG 46:2\|TG 14:0_16:1_16:1 | 0.148342 | 0.19059 |
| DG 42:10 | 1.186349 | 0.190849 |
| PE 36:2\|PE 18:0_18:2 | 0.025877 | 0.192589 |
| PE-Cer 36:0;3O | -0.11291 | 0.197866 |
| PE 36:2;2O\|PE 18:0_18:2;2O | 0.303135 | 0.198431 |
| Cer 43:1;4O | 0.211922 | 0.199517 |
| TG 50:0\|TG 16:0_16:0_18:0 | 0.219859 | 0.201593 |
| PS 36:4\|PS 18:2_18:2 | -0.16848 | 0.202382 |
| PG 34:3\|PG 16:1_18:2 | 0.162395 | 0.203457 |
| PC 38:2\|PC 20:0_18:2 | -0.14798 | 0.20545 |
| TG O-40:0\|TG O-14:0_12:0_14:0 | 0.648643 | 0.20643 |
| DG 35:5 | 0.972671 | 0.21026 |
| PE-Cer 38:2;2O | -0.16772 | 0.211967 |
| FA 32:0 | -0.32085 | 0.212697 |
| DG 34:2\|DG 16:0_18:2 | -0.15825 | 0.213362 |
| PC O-39:7 | 0.308393 | 0.21338 |
| PC 36:4 | -0.05441 | 0.215074 |
| PE 40:2 | -0.13977 | 0.218677 |
| PE P-36:4\|PE P-18:2_18:2 | -0.0769 | 0.221861 |
| PS 35:1\|PS 10:0_25:1 | -0.172 | 0.223436 |
| CL 66:5\|CL 16:1_16:1_16:1_18:2 | -0.17455 | 0.225181 |
| PE 35:1\|PE 17:0_18:1 | -0.06653 | 0.225957 |
| PS 36:5\|PS 18:2_18:3 | -0.14429 | 0.226398 |
| TG O-38:0\|TG O-14:0_12:0_12:0 | 0.256364 | 0.22738 |
| PE O-36:2\|PE O-18:0_18:2 align_ID:1392 | 0.109843 | 0.22864 |
| PE 33:2;2O\|PE 16:1_17:1;2O | -0.4051 | 0.229642 |
| PE 36:2;O\|PE 18:2_18:0;O | 0.478973 | 0.230094 |
| Cer 42:0;2O | 0.22773 | 0.230174 |

| | | |
|---|---|---|
| Cer 44:0;2O | 0.279328 | 0.230437 |
| PC O-37:1 | -0.12083 | 0.231343 |
| PA 36:3\|PA 18:1_18:2 | 0.233108 | 0.231678 |
| PE-Cer 36:1;2O\|PE-Cer 18:0;2O/18:1 | -0.06686 | 0.2317 |
| PI 34:0\|PI 16:0_18:0 | -0.33522 | 0.232586 |
| PC 35:5 | -0.2001 | 0.233172 |
| FA 11:0 | -0.20573 | 0.23323 |
| PE 27:1;O\|PE 10:0_17:1;O align_ID:1108 | -0.15053 | 0.234533 |
| CL 70:7\|CL 16:1_18:2_18:2_18:2 | -0.08665 | 0.235521 |
| PI 34:1\|PI 16:0_18:1 | -0.18791 | 0.239285 |
| PC 38:7 | 0.187499 | 0.240701 |
| MG 37:1 | 0.691385 | 0.242065 |
| PE 36:1;2O\|PE 18:1_18:0;2O | -0.11076 | 0.242917 |
| PE 27:0 | -0.16681 | 0.244153 |
| DG 30:3 | 0.735388 | 0.244464 |
| SM 42:7;2O | 0.280594 | 0.246863 |
| PC 36:5 | 0.113929 | 0.247221 |
| Cer 44:1;4O\|Cer 18:1;3O/26:0;(2OH) | 0.214228 | 0.247379 |
| TG 43:0\|TG 14:0_14:0_15:0 | -0.31354 | 0.247851 |
| FA 34:0 | -0.59415 | 0.248547 |
| TG 53:1\|TG 17:0_18:0_18:1 | 0.130023 | 0.248692 |
| PC 32:3 | 0.124698 | 0.250901 |
| LPE O-18:0 | 0.098583 | 0.253329 |
| PG 36:5\|PG 18:2_18:3 | -0.10591 | 0.254444 |
| PC 34:2;O\|PC 18:1_16:1;O | -0.20956 | 0.255809 |
| PC 34:4 | 0.299831 | 0.256201 |
| NAE 22:5 | -0.47294 | 0.257286 |
| PC 33:2 | -0.09235 | 0.259348 |
| PC 28:1 | -0.17084 | 0.259934 |
| TG 47:1\|TG 14:0_15:0_18:1 | 0.212986 | 0.260955 |
| PI 34:3 | 0.18355 | 0.261187 |
| PC O-38:2 | -0.09608 | 0.262391 |
| ST 28:3;O align_ID:1222 | -0.0607 | 0.262771 |
| PE-Cer 33:1;2O | 0.124847 | 0.2636 |
| CL 78:5\|CL 18:0_24:0_18:2_18:3 | 0.263579 | 0.268449 |
| FAHFA 30:0;O\|FAHFA 16:0/14:0;O | -0.44414 | 0.268536 |
| PC O-40:9 | 0.271072 | 0.270001 |
| PC 36:2\|PC 18:1_18:1 | -0.01115 | 0.271527 |
| FA 18:1 align_ID:355 | -0.33699 | 0.271844 |
| PE-Cer 36:1;2O\|PE-Cer 14:1;2O/22:0 | 0.079677 | 0.272875 |
| PE-Cer 38:0;2O\|PE-Cer 14:0;2O/24:0 | 0.146242 | 0.273477 |
| DG O-33:1\|DG O-17:0_16:1 | -0.90441 | 0.274726 |
| PS 34:3 | -0.20297 | 0.274917 |
| PC O-36:3 | -0.09123 | 0.277852 |
| TG 47:5;1O\|TG 14:1_18:2_15:2;1O | 0.099174 | 0.280821 |
| PC 34:0\|PC 16:0_18:0 | -0.06495 | 0.282398 |

| | | |
|---|---|---|
| DG 36:1\|DG 18:0_18:1 | -0.1361 | 0.287377 |
| TG 26:0\|TG 8:0_8:0_10:0 | -1.03742 | 0.288271 |
| PE 30:2;O\|PE 16:1_14:1;O | -0.15871 | 0.290113 |
| PE O-35:2\|PE O-17:0_18:2 | 0.069164 | 0.291389 |
| PG 34:4 | -0.15096 | 0.294813 |
| TG 24:0\|TG 8:0_8:0_8:0 | -1.01009 | 0.294842 |
| PE-Cer 36:2;2O\|PE-Cer 14:2;2O/22:0 | -0.10189 | 0.297339 |
| TG 28:0\|TG 8:0_10:0_10:0 | -1.25253 | 0.300077 |
| PC 32:0\|PC 16:0_16:0 | -0.05155 | 0.301714 |
| PE 34:2 | 0.078933 | 0.306821 |
| TG 49:1\|TG 15:0_16:0_18:1 | 0.208163 | 0.309291 |
| TG 45:0\|TG 14:0_15:0_16:0 | -0.18559 | 0.310598 |
| PC 36:0\|PC 18:0_18:0 | -0.10385 | 0.310711 |
| PI 36:5\|PI 18:2_18:3 | 0.107592 | 0.311591 |
| ST 28:4;O align_ID:1098 | 0.495163 | 0.311858 |
| DG 34:0\|DG 16:0_18:0 | -0.13367 | 0.315365 |
| DG O-35:1\|DG O-19:0_16:1 | -0.56597 | 0.316767 |
| PC 32:2\|PC 16:1_16:1 | 0.055096 | 0.31769 |
| DG 50:6 | 0.067331 | 0.317775 |
| PC 36:5\|PC 18:2_18:3 | -0.04886 | 0.318529 |
| FA 18:2 align_ID:347 | -0.22098 | 0.321229 |
| PE 32:2\|PE 16:1_16:1 | 0.158653 | 0.321301 |
| PC O-40:8 | 0.219513 | 0.321565 |
| PI 36:2 | -0.14035 | 0.321597 |
| LPE O-20:0 | 0.09013 | 0.324936 |
| LPC 18:1/0:0 | 0.12374 | 0.331229 |
| SE 28:1/18:2 | -0.35946 | 0.331266 |
| TG 52:1\|TG 16:0_18:0_18:1 | 0.174479 | 0.333395 |
| CL 70:6\|CL 16:0_18:2_18:2_18:2 | -0.09453 | 0.334129 |
| PC 38:6\|PC 16:0_22:6 | -0.1735 | 0.334675 |
| TG O-50:2\|TG O-18:0_16:1_16:1 | 0.229021 | 0.335881 |
| PS 34:4 | -0.19038 | 0.336775 |
| PC O-33:2 align_ID:3468 | -0.1122 | 0.336875 |
| TG 44:0\|TG 14:0_14:0_16:0 | -0.24309 | 0.338124 |
| MG 18:2 | 0.142081 | 0.339061 |
| Cer 41:1;4O | 0.162815 | 0.339147 |
| PC O-35:2 | 0.041657 | 0.339199 |
| CL 72:8\|CL 18:2_18:2_18:2_18:2 | -0.14687 | 0.339785 |
| PC O-34:8 | 0.731622 | 0.34198 |
| ST 29:1;O align_ID:1448 | -0.35678 | 0.342657 |
| CL 77:7\|CL 39:3_38:4 | 0.177327 | 0.347813 |
| CL 82:9\|CL 28:0_18:3_18:3_18:3 | 0.236794 | 0.349847 |
| SM 39:8;3O | 0.102525 | 0.35048 |
| PC 36:1 | 0.33717 | 0.351413 |
| SM 61:6;2O | 0.121215 | 0.352117 |
| Cer 28:0;4O | 1.877883 | 0.3525 |

| | | |
|---|---|---|
| TG 62:1\|TG 14:0_16:0_32:1 | 0.149248 | 0.357833 |
| TG 50:1\|TG 16:0_16:0_18:1 | 0.144968 | 0.358443 |
| NAE 22:6 | 0.589812 | 0.359146 |
| FA 34:4 | -0.33571 | 0.361281 |
| CL 68:6\|CL 16:1_16:1_18:2_18:2 | -0.08734 | 0.364415 |
| PS 34:3\|PS 16:1_18:2 | -0.12926 | 0.370838 |
| DG 32:1\|DG 16:0_16:1 | -0.1119 | 0.371203 |
| Cer 46:1;4O | 0.153899 | 0.378399 |
| TG 40:2\|TG 12:0_14:1_14:1 | -0.19661 | 0.381979 |
| DG 25:1 | 0.372071 | 0.383883 |
| Cer 64:13;4O\|Cer 46:9;3O(FA 18:3) | -0.09121 | 0.386662 |
| TG 46:0\|TG 14:0_16:0_16:0 | -0.16379 | 0.389221 |
| PE-Cer 37:1;2O | 0.099584 | 0.392271 |
| PE O-34:3 | -0.04023 | 0.393595 |
| Cer 48:0;3O | 0.28116 | 0.395299 |
| FA 24:0 align_ID:557 | -0.10187 | 0.396949 |
| TG 54:7\|TG 18:2_18:2_18:3 | 0.200057 | 0.399093 |
| FA 20:1 | -0.28837 | 0.400043 |
| PE 36:2\|PE 18:1_18:1 | 0.073123 | 0.401146 |
| ST 27:1;O;S | 0.245453 | 0.403745 |
| PE O-37:2 | 0.04376 | 0.404013 |
| PC 35:4 | -0.12699 | 0.404564 |
| SHexCer 33:2;3O | -0.11475 | 0.406962 |
| PE 36:3\|PE 18:0_18:3 | 0.024416 | 0.407396 |
| PC O-38:9 | 0.445956 | 0.408666 |
| PE O-36:6\|PE O-18:4_18:2 | -0.09294 | 0.411358 |
| PE 40:2\|PE 22:0_18:2 | -0.09037 | 0.411898 |
| NAOrn 18:2;O | -0.12079 | 0.412615 |
| FA 18:0 align_ID:361 | -0.08267 | 0.413559 |
| FA 28:0 | -0.18103 | 0.413779 |
| PC O-38:4 | -0.45844 | 0.413836 |
| PI 36:2\|PI 18:1_18:1 | -0.13478 | 0.419706 |
| FA 32:1 | 0.235209 | 0.426114 |
| PE 36:5 | 0.103751 | 0.426808 |
| PE 35:2;O\|PE 18:1_17:1;O | 0.093676 | 0.429855 |
| FA 18:1 align_ID:352 | -0.09817 | 0.429912 |
| DG 34:1\|DG 16:0_18:1 | -0.08448 | 0.431671 |
| FA 22:1 | -0.24278 | 0.431947 |
| PC O-38:10 | -0.43997 | 0.432056 |
| PE O-36:5\|PE O-18:3_18:2 | -0.07516 | 0.432988 |
| PI 36:6\|PI 18:3_18:3 | 0.176109 | 0.434521 |
| PE-Cer 36:5;2O | 0.063246 | 0.437534 |
| PS 35:2;O\|PS 18:1_17:1;O | -0.10507 | 0.438696 |
| MG 36:3 | -0.1228 | 0.438861 |
| FA 22:0 | -0.11833 | 0.439473 |
| FA 19:0 | -0.19728 | 0.439503 |

| | | |
|---|---|---|
| ST 28:4;O align_ID:1100 | 0.172844 | 0.439866 |
| PE O-38:0\|PE O-18:0_20:0 | -0.08847 | 0.442671 |
| PE-Cer 32:1;3O | -0.12297 | 0.44352 |
| PE 34:4\|PE 16:1_18:3 | 0.114507 | 0.444333 |
| PI 36:5 | 0.08281 | 0.444532 |
| SE 29:1/16:2 | 0.112521 | 0.445187 |
| FA 26:0 | -0.13975 | 0.446876 |
| Cer 36:1;2O\|Cer 14:1;2O/22:0 | -0.03576 | 0.454029 |
| PE O-38:2\|PE O-20:0_18:2 | 0.030648 | 0.457934 |
| DG 36:4\|DG 18:2_18:2 | 0.075391 | 0.462409 |
| ST 27:1;O | 0.200848 | 0.464433 |
| PC 38:6 | -0.23999 | 0.468569 |
| PC O-41:7 | 0.284139 | 0.46988 |
| DG 33:1\|DG 15:0_18:1 | -0.15481 | 0.474307 |
| PE O-36:3\|PE O-18:0_18:3 | 0.041086 | 0.478412 |
| Cer 46:0;3O\|Cer 22:0;3O/24:0 | 0.180527 | 0.479519 |
| DG 49:13 | 0.267958 | 0.480642 |
| Cer 34:1;3O\|Cer 18:1;2O/16:0;(2OH) | 0.142072 | 0.48153 |
| PE O-40:5\|PE O-22:3_18:2 | 0.133816 | 0.481607 |
| DG 26:4 | 0.321524 | 0.482853 |
| TG 39:1;1O\|TG 10:0_16:0_13:1;1O | 0.173599 | 0.484212 |
| FA 27:0 align_ID:639 | 0.175431 | 0.486155 |
| PC 32:1\|PC 16:0_16:1 | 0.011789 | 0.48728 |
| PC 34:4\|PC 16:1_18:3 | -0.06188 | 0.488049 |
| Cer 42:1;3O\|Cer 18:1;2O/24:0;O | 0.122937 | 0.489626 |
| TG 48:0\|TG 14:0_16:0_18:0 | -0.10133 | 0.490957 |
| Cer 42:1;4O | 0.13469 | 0.492127 |
| MG 38:1 | 0.125119 | 0.494479 |
| ST 27:2;O | 0.067778 | 0.496432 |
| PC O-37:5 | 0.030978 | 0.497726 |
| Cer 44:0;3O\|Cer 20:0;3O/24:0 | 0.158233 | 0.498697 |
| SM 63:7;2O | -0.07973 | 0.501324 |
| Cer 44:1;3O\|Cer 19:0;2O/25:1;O | 0.124362 | 0.502582 |
| FA 23:0 | -0.15307 | 0.505333 |
| PC 36:4\|PC 18:2_18:2 | 0.012111 | 0.506056 |
| PE-Cer 38:1;2O\|PE-Cer 14:1;2O/24:0 | 0.054074 | 0.509299 |
| PC 37:8 | 0.06314 | 0.515844 |
| PI 34:2 | -0.06618 | 0.516515 |
| SM 36:2;3O align_ID:3627 | -0.17005 | 0.517577 |
| PI 34:3\|PI 16:1_18:2 | -0.06571 | 0.519391 |
| PE 34:3\|PE 16:0_18:3 | 0.034502 | 0.519518 |
| PE-Cer 32:0;3O | 0.069755 | 0.519951 |
| PE 32:2 | -0.07487 | 0.522326 |
| TG 50:5\|TG 16:1_16:1_18:3 | 0.179852 | 0.522962 |
| PE P-36:6\|PE P-18:3_18:3 | -0.07335 | 0.524611 |
| PE O-36:4 | 0.045955 | 0.530044 |

| | | |
|---|---|---|
| Cer 42:0;3O\|Cer 18:0;3O/24:0 | 0.137967 | 0.530733 |
| PG 36:4\|PG 18:2_18:2 | 0.059663 | 0.531127 |
| DG 48:11 | -0.04598 | 0.53363 |
| PI 36:4\|PI 18:2_18:2 | 0.046932 | 0.544975 |
| SM 43:1;3O | -0.03981 | 0.547301 |
| FA 20:2 | -0.26123 | 0.547727 |
| PC 37:4 | -0.07976 | 0.548196 |
| PE 36:0\|PE 18:0_18:0 | -0.04766 | 0.549107 |
| PS 34:4\|PS 16:1_18:3 | -0.09161 | 0.549438 |
| Cer 46:1;3O\|Cer 20:1;2O/26:0;O | 0.115048 | 0.551654 |
| TG 45:2\|TG 13:0_16:1_16:1 | 0.09402 | 0.551891 |
| PC O-39:9 | -0.15779 | 0.553447 |
| PE-Cer 34:1;3O | 0.06646 | 0.554043 |
| PC O-37:2 | 0.056844 | 0.557379 |
| PE O-34:1\|PE O-17:0_17:1 | 0.03327 | 0.557543 |
| PE 36:1\|PE 18:0_18:1 | -0.0259 | 0.560079 |
| PC O-32:1 | 0.025313 | 0.561863 |
| PC O-33:0 | 0.046444 | 0.56782 |
| Cer 43:0;3O | 0.11472 | 0.569104 |
| Cer 44:1;3O\|Cer 18:1;2O/26:0;O | 0.107557 | 0.569239 |
| FA 16:0 align_ID:301 | 0.130864 | 0.569898 |
| FA 19:1 | -0.22171 | 0.574265 |
| DG 22:1 | -0.31793 | 0.576415 |
| SM 36:2;3O align_ID:3629 | -0.03718 | 0.576985 |
| DGGA 32:0\|DGGA 16:0_16:0 | 0.051149 | 0.576987 |
| PI 34:2\|PI 16:1_18:1 | -0.06737 | 0.578804 |
| FA 32:2 | -0.14437 | 0.579963 |
| SM 44:7;2O\|SM 14:1;2O/30:6 | -0.04034 | 0.5833 |
| Cer 42:1;3O\|Cer 21:1;2O/21:0;O | 0.114614 | 0.583478 |
| CL 80:3\|CL 18:0_26:0_18:1_18:2 | -0.14368 | 0.587722 |
| PG 35:3 | -0.08948 | 0.588548 |
| Cer 35:1;2O\|Cer 14:1;2O/21:0 | -0.18286 | 0.589912 |
| DG 36:5\|DG 18:2_18:3 | -0.048 | 0.593002 |
| SM 41:0;3O | -0.0404 | 0.593697 |
| TG 39:1\|TG 12:0_13:0_14:1 | -0.09977 | 0.597734 |
| PC 30:3 | 0.051521 | 0.598483 |
| PE O-40:3\|PE O-22:0_18:3 | 0.047389 | 0.599426 |
| FA 18:0 align_ID:367 | 0.128307 | 0.601825 |
| ST 28:4;O align_ID:1213 | 0.122701 | 0.602028 |
| PC 37:3 | 0.071395 | 0.602468 |
| PC 33:1\|PC 16:0_17:1 | 0.23601 | 0.604156 |
| Cer 44:1;3O\|Cer 26:0;2O/18:1;O | 0.116037 | 0.605412 |
| PE O-34:0 | -0.03318 | 0.606415 |
| PI 36:3 | 0.03705 | 0.612891 |
| PE O-34:0\|PE O-18:0_16:0 | -0.03149 | 0.61341 |
| PG 36:3\|PG 18:1_18:2 | 0.055519 | 0.628078 |

| | | |
|---|---|---|
| PC 34:3\|PC 16:1_18:2 | -0.02094 | 0.630858 |
| DG 30:0 | 0.352271 | 0.63558 |
| LPC 18:0/0:0 | -0.05049 | 0.636668 |
| Cer 46:0;3O\|Cer 24:0;2O/22:0;O | 0.107029 | 0.636872 |
| PE O-37:3\|PE O-19:0_18:3 | -0.04416 | 0.639648 |
| PC O-33:2 align_ID:3465 | -0.1109 | 0.640516 |
| TG 43:1\|TG 13:0_14:0_16:1 | -0.08824 | 0.642499 |
| PE 35:4\|PE 17:2_18:2 | 0.071945 | 0.65044 |
| DG 44:9 | -0.04531 | 0.657847 |
| MG 26:6 | 0.267641 | 0.662036 |
| TG 27:0\|TG 9:0_9:0_9:0 | 0.159221 | 0.66247 |
| PC O-39:1 | 0.127705 | 0.663906 |
| FA 10:0 | 0.044223 | 0.666105 |
| PC 33:2\|PC 15:0_18:2 | 0.012831 | 0.668628 |
| TG 47:0\|TG 15:0_16:0_16:0 | -0.0729 | 0.669801 |
| TG 44:2\|TG 12:0_16:1_16:1 | 0.04805 | 0.672177 |
| ST 28:1;O | -0.41418 | 0.674925 |
| PE-Cer 34:1;3O\|PE-Cer 18:1;2O/16:0;O | 0.032668 | 0.678552 |
| FA 24:1 | 0.132232 | 0.683769 |
| CL 80:5\|CL 18:0_26:0_18:2_18:3 | -0.11611 | 0.685008 |
| CL 68:6\|CL 16:1_18:2_16:1_18:2 | -0.04623 | 0.68588 |
| FA 29:0 | 0.077909 | 0.687766 |
| MGDG 35:2 | -0.09505 | 0.694039 |
| PE 36:2 | 0.027994 | 0.69573 |
| PC 34:2\|PC 16:1_18:1 | -0.00391 | 0.69643 |
| NAE 9:0 | -0.07043 | 0.69746 |
| FA 30:1 | -0.09335 | 0.702686 |
| PG 34:3\|PG 16:0_18:3 | 0.040078 | 0.711847 |
| PI 36:6 | 0.074747 | 0.714444 |
| PE 36:6\|PE 18:3_18:3 | 0.051929 | 0.716145 |
| FA 21:0 align_ID:460 | -0.0737 | 0.717258 |
| PE-Cer 37:3;3O | -0.04548 | 0.717443 |
| PI 36:3\|PI 18:1_18:2 | -0.03288 | 0.718913 |
| DG 33:6 | -0.41889 | 0.720679 |
| FA 25:0 | -0.09837 | 0.725052 |
| DG O-36:2\|DG O-18:0_18:2 | 0.044145 | 0.727814 |
| PC O-39:2 | -0.02495 | 0.728731 |
| LPC 16:0/0:0 | 0.032529 | 0.732817 |
| PC 36:3\|PC 18:1_18:2 | 0.006615 | 0.735467 |
| PE 34:2;3O\|PE 16:1_18:1;3O | -0.05247 | 0.746216 |
| TG 39:2;1O\|TG 10:0_16:1_13:1;1O | 0.075195 | 0.746507 |
| Cer 48:11;4O\|Cer 13:1;2O/35:10;2O | -0.01557 | 0.747671 |
| PE 32:0\|PE 16:0_16:0 | 0.019335 | 0.754781 |
| CL 73:6 | 0.017136 | 0.756252 |
| Cer 42:1;3O\|Cer 19:0;2O/23:1;O | 0.056369 | 0.757368 |
| SE 29:1/18:2 | 0.030279 | 0.760168 |

| | | |
|---|---|---|
| TG 57:8;1O\|TG 22:3_21:5_14:0;1O | -0.28887 | 0.761783 |
| NAE 18:4 | -0.06338 | 0.762279 |
| PI 34:4\|PI 16:1_18:3 | -0.05922 | 0.762281 |
| PC 35:4;O\|PC 18:2_17:2;O | 0.125023 | 0.765125 |
| TG 45:1\|TG 14:0_15:0_16:1 | 0.052054 | 0.768061 |
| PC O-35:0 | -0.0314 | 0.769417 |
| PC 40:7 | -0.0412 | 0.769608 |
| DG 28:4 | 0.123184 | 0.775687 |
| PC O-32:2 | -0.09284 | 0.777474 |
| PC O-38:8 | -0.29521 | 0.778635 |
| DG 52:10 | -0.05274 | 0.778976 |
| PE O-37:2\|PE O-19:0_18:2 | 0.028414 | 0.784115 |
| TG 54:8\|TG 18:2_18:3_18:3 | 0.085391 | 0.784894 |
| Cer 42:1;3O\|Cer 18:1;2O/24:0;(2OH) | 0.049979 | 0.79356 |
| Cer 34:0;3O\|Cer 14:0;3O/20:0 | 0.022445 | 0.794371 |
| PE O-36:3 | 0.018186 | 0.795809 |
| Cer 38:1;2O\|Cer 14:1;2O/24:0 | 0.019502 | 0.797145 |
| FA 18:2 align_ID:346 | 0.037034 | 0.797854 |
| TG 41:1\|TG 13:0_14:0_14:1 | -0.0516 | 0.808947 |
| 4-Imidazoleacrylic acid | 0.142434 | 0.809544 |
| PE-Cer 33:1;3O\|PE-Cer 14:0;2O/19:1;O | -0.04183 | 0.80992 |
| PC O-36:9 | -0.08413 | 0.811949 |
| PC O-35:1 | 0.019428 | 0.813517 |
| PE 30:1 | 0.030372 | 0.813791 |
| CL 78:6\|CL 24:0_18:2_18:2_18:2 | -0.02298 | 0.818374 |
| PE O-35:3\|PE O-17:0_18:3 | 0.026152 | 0.819691 |
| Cer 34:1;4O | -0.05055 | 0.826383 |
| TG 48:1\|TG 14:0_16:0_18:1 | 0.040994 | 0.827834 |
| PI 34:4 | 0.027543 | 0.828531 |
| TG 49:0\|TG 15:0_16:0_18:0 | 0.030945 | 0.829233 |
| PC O-35:3 align_ID:3729 | -0.01671 | 0.829346 |
| PE O-40:7\|PE O-22:5_18:2 | 0.027132 | 0.832161 |
| FA 26:1 | -0.03768 | 0.832372 |
| CAR 18:0 | -0.0217 | 0.835977 |
| PC 36:2 align_ID:4298 | -0.01473 | 0.84478 |
| DG 36:3\|DG 18:1_18:2 | 0.027058 | 0.848395 |
| PE O-38:3 | 0.011184 | 0.850203 |
| PC O-32:3 | -0.02003 | 0.851824 |
| PC 33:1\|PC 15:0_18:1 | 0.003884 | 0.860694 |
| FA 19:2 | -0.06593 | 0.862391 |
| ST 28:3;O align_ID:1116 | -0.02488 | 0.865299 |
| PE O-36:0\|PE O-18:0_18:0 | 0.017718 | 0.867228 |
| FA 18:3 align_ID:341 | 0.028461 | 0.868105 |
| PE 33:2 | -0.01767 | 0.869746 |
| Dioctyl phthalate | -0.04245 | 0.874145 |
| PG 36:1\|PG 18:0_18:1 | 0.018513 | 0.875531 |

| | | |
|---|---|---|
| CL 76:3\|CL 16:0_24:0_18:1_18:2 | -0.02368 | 0.877483 |
| NAE 4:0 | -0.04161 | 0.881176 |
| DG 50:7 | -0.01498 | 0.883934 |
| PC 40:8 | -0.02224 | 0.885632 |
| NAE 19:0 | -0.03572 | 0.89169 |
| SM 39:4;3O | -0.0105 | 0.894572 |
| PE 25:1;O\|PE 10:0_15:1;O | -0.02352 | 0.897096 |
| TG 46:1\|TG 14:0_16:0_16:1 | -0.017 | 0.899574 |
| FA 34:3 | 0.038557 | 0.90032 |
| SM 39:2;3O | -0.01168 | 0.904147 |
| CL 72:7\|CL 18:1_18:2_18:2_18:2 | 0.015247 | 0.906176 |
| CL 76:5\|CL 24:0_16:1_18:1_18:3 | 0.013094 | 0.908406 |
| TG 45:2;1O\|TG 16:0_16:1_13:1;1O | 0.15964 | 0.914135 |
| PE O-40:2\|PE O-22:0_18:2 | -0.00964 | 0.914685 |
| PS 32:2 | -0.02941 | 0.916541 |
| PC O-37:4 | -0.00914 | 0.916618 |
| Cer 40:1;2O\|Cer 18:1;2O/22:0 | 0.016884 | 0.91689 |
| TG 52:1;4O\|TG 14:0_16:0_22:1;4O | 0.026854 | 0.921108 |
| Cer 42:1;2O\|Cer 18:1;2O/24:0 | 0.020476 | 0.92485 |
| PE O-36:7\|PE O-18:4_18:3 | -0.01218 | 0.925724 |
| PE-Cer 38:3;2O | 0.01487 | 0.930337 |
| PE O-38:1\|PE O-20:0_18:1 | 0.005514 | 0.932069 |
| TG 43:2\|TG 13:0_14:1_16:1 | 0.015446 | 0.933768 |
| DG 36:2\|DG 18:0_18:2 | 0.006255 | 0.937913 |
| PE 35:2\|PE 17:0_18:2 | -0.00432 | 0.938718 |
| CL 74:7 | 0.01085 | 0.938918 |
| TG 37:1;1O\|TG 10:0_14:0_13:1;1O | 0.018843 | 0.939558 |
| PC 32:2 | -0.00532 | 0.940085 |
| SM 37:1;2O\|SM 21:0;2O/16:1 | 0.009248 | 0.952602 |
| PC 36:3 | -0.0037 | 0.95993 |
| TG O-42:0\|TG O-16:0_12:0_14:0 | -0.0112 | 0.960775 |
| Cer 32:1;2O\|Cer 14:1;2O/18:0 | -0.00535 | 0.962303 |
| ST 28:4;O align_ID:1101 | 0.004671 | 0.96836 |
| PE 35:3\|PE 17:1_18:2 | -0.00344 | 0.977119 |
| Cer 54:11;4O\|Cer 12:2;2O/42:9;2O | 0.001909 | 0.977877 |
| DG 35:1\|DG 17:0_18:1 | 0.004754 | 0.978781 |
| Cer 40:1;2O\|Cer 16:1;2O/24:0 | 0.003886 | 0.980095 |
| Cer 34:1;2O\|Cer 14:1;2O/20:0 | 0.001612 | 0.981432 |
| DG 37:5 | -0.00208 | 0.982491 |
| TG 42:2\|TG 12:0_14:1_16:1 | -0.00403 | 0.983023 |
| LPE 18:0 | 0.001967 | 0.983618 |
| PE-Cer 34:2;2O\|PE-Cer 14:2;2O/20:0 | 0.001608 | 0.98471 |
| PC O-32:1\|PC O-16:0_16:1 | -0.00188 | 0.985972 |
| PE O-38:3\|PE O-20:0_18:3 | -0.00102 | 0.986956 |
| PI 32:2 | -0.00141 | 0.989863 |
| FA 9:0 align_ID:131 | -0.00272 | 0.990704 |

| | | |
|---|---|---|
| PE-Cer 38:5;2O | -0.00177 | 0.991337 |
| PC 34:4\|PC 18:1_16:3 | -0.00047 | 0.993358 |
| PC O-35:4 | 0.00031 | 0.993954 |
| PE O-34:3\|PE O-16:0_18:3 | -0.00019 | 0.996585 |
| ST 29:2;O | -0.00029 | 0.999073 |
| PE 34:3 | 0 | 1 |
