## Supplemental Table 2 for "Neuronal lipid droplets play a conserved and sex-biased role in maintaining whole-body energy homeostasis"

| Differentially regulated genes in Female brains with neuronal loss of *dATGL* | | | | |
|---|---|---|---|---|
| | RNAi / GAL4 control | | RNAi / UAS control | |
| Genes | log2FC | p adjusted | log2FC | p adjusted |
| AstA | 0.209425 | 0.011259 | 0.251449 | 0.002984 |
| CG10924 | 3.005566 | 1.41E-05 | 2.131678 | 0.004519 |
| CG10960 | 0.236739 | 0.024442 | 0.251505 | 0.040467 |
| CG11893 | -3.44709 | 0.005232 | -3.43574 | 0.017519 |
| CG12512 | 0.665365 | 0.001383 | 0.618936 | 0.013166 |
| CG12861 | 15.29163 | 0.004718 | 15.42658 | 0.01414 |
| CG14715 | -1.07968 | 6.31E-09 | -0.9477 | 4.31E-06 |
| CG2127 | -13.7017 | 0.00035 | -13.7352 | 0.001214 |
| CG3160 | -0.27639 | 0.01131 | -0.33506 | 0.002334 |
| CG3842 | -1.90183 | 0.000135 | -2.08315 | 4.76E-05 |
| CG4078 | 0.399897 | 5.66E-08 | 0.477159 | 1.29E-10 |
| CG6654 | 0.453096 | 0.000182 | -0.53934 | 5.54E-06 |
| CG6834 | -1.19472 | 5.97E-09 | -0.89813 | 0.000278 |
| CG6912 | 0.602418 | 0.0044 | -0.59533 | 0.015754 |
| CG6928 | -0.89832 | 1.43E-08 | -0.60931 | 0.003347 |
| CG7054 | -0.69262 | 0.020916 | -0.91952 | 0.001042 |
| CHKov1 | -0.8512 | 0.000182 | -0.8198 | 0.001403 |
| CHKov2 | -0.67891 | 0.002386 | -0.74363 | 0.001858 |
| CR42547 | 0.506121 | 0.019315 | 0.609182 | 0.00579 |
| CR45214 | 0.932749 | 0.003442 | 0.827601 | 0.044164 |
| CR45244 | 0.960755 | 1.64E-10 | 0.92426 | 3.17E-09 |
| CR45600 | 0.826563 | 2.38E-06 | -0.56743 | 0.017519 |
| GM130 | -0.4684 | 0.000601 | -0.60661 | 4.26E-06 |
| Hsp70Bb | 2.788584 | 9.51E-05 | 2.345705 | 0.006917 |
| Mp20 | -4.68722 | 0.000309 | -4.09821 | 0.010377 |
| mTerf3 | -1.59864 | 9.17E-11 | -1.87289 | 2.02E-14 |
| Myo28B1 | 0.523335 | 0.001231 | 0.798936 | 7.92E-08 |
| NimC1 | 1.442862 | 0.000117 | -1.03904 | 0.019862 |
| Spindly | -2.19827 | 0.000601 | -2.26214 | 0.001271 |
| Tk | 0.397737 | 1.17E-06 | 0.37918 | 1.98E-05 |
| w | 1.930631 | 4.27E-17 | 1.181114 | 1.11E-05 |
| **Differentially regulated genes in Male brains with neuronal loss of *dATGL*** | | | | |
| | RNAi / GAL4 control | | RNAi / UAS control | |
| Genes | log2FC | p adjusted | log2FC | p adjusted |
| CG10562 | -0.85009 | 5.13E-06 | -0.81255 | 5.48E-05 |
| CG11893 | -4.2678 | 0.017644 | -4.76124 | 0.009816 |
| CG13502 | -1.3115 | 0.007421 | -1.41748 | 0.005885 |

| | | | | |
|---|---|---|---|---|
| CG14715 | -1.06443 | 1.87E-07 | -0.78802 | 0.002728 |
| CG6293 | -0.64092 | 0.026929 | 0.792104 | 0.009985 |
| CG6654 | 0.341749 | 0.025698 | -0.54542 | 4.33E-06 |
| Cp15 | -16.6405 | 1.23E-05 | -16.3003 | 6.23E-05 |
| CR45244 | 0.86234 | 1.27E-07 | 0.725938 | 6.23E-05 |
| GM130 | -0.39144 | 0.017288 | -0.43722 | 0.009816 |
| Hsp70Bb | 2.764664 | 0.000182 | 3.144014 | 1.80E-05 |
| mre11 | 0.536346 | 6.44E-08 | -0.32801 | 0.010466 |
| mTerf3 | -0.9834 | 0.00108 | -1.26842 | 4.01E-06 |
| Obp99b | -1.61699 | 0.025698 | -1.79279 | 0.016866 |
| Spindly | -1.86869 | 0.028808 | -2.13523 | 0.012013 |
| Su(P) | -0.34491 | 0.012295 | -0.35225 | 0.020744 |
| w | 1.777906 | 2.80E-14 | 1.817485 | 3.82E-15 |

**Differentially regulated genes between Female and Male brains**

| | GAL4 control | | UAS control | | RNAi | |
|---|---|---|---|---|---|---|
| **Genes** | **log2FC** | **p adjusted** | **log2FC** | **p adjusted** | **log2FC** | **p adjusted** |
| Ac78C | -0.27099 | 0.007318 | 0.272124 | 0.011058 | -0.17907 | 0.132404 |
| Acer | 0.27125 | 0.029647 | 0.281298 | 0.021599 | 0.397897 | 0.000184 |
| ade5 | -0.74134 | 8.92E-06 | -0.6593 | 0.000147 | -0.45475 | 0.022897 |
| AIF | -0.23402 | 0.013956 | -0.20638 | 0.047982 | -0.1757 | 0.100375 |
| AMPdeam | 0.234903 | 4.83E-07 | 0.220914 | 3.56E-06 | 0.197881 | 5.50E-05 |
| Ant2 | 1.058096 | 0.007939 | 0.934543 | 0.024674 | 1.276684 | 0.000396 |
| AnxB10 | 0.461694 | 5.11E-05 | 0.346536 | 0.007612 | 0.489817 | 1.13E-05 |
| AnxB11 | 0.41925 | 6.59E-05 | 0.436859 | 3.54E-05 | 0.402028 | 0.000184 |
| asl | 0.519668 | 0.026184 | 0.533124 | 0.024504 | 0.278033 | 0.303337 |
| Axs | -0.41401 | 0.022834 | -0.66969 | 1.58E-05 | -0.59786 | 0.000191 |
| Bap60 | 0.307116 | 0.000163 | 0.314659 | 0.000142 | 0.323706 | 5.72E-05 |
| brv3 | -5.97097 | 8.85E-16 | -5.16803 | 2.68E-09 | -5.97214 | 9.46E-13 |
| bys | -0.33958 | 0.026678 | -0.50006 | 0.00014 | -0.42831 | 0.001928 |

| | | | | | | |
|---|---|---|---|---|---|---|
| c12.1 | 0.385692 | 0.008619 | 0.384405 | 0.009454 | 0.398501 | 0.005347 |
| CalpC | -0.9632 | 9.17E-05 | -0.68512 | 0.025495 | -1.30261 | 7.01E-08 |
| CG10352 | -0.71291 | 5.20E-05 | -0.80535 | 2.62E-06 | -0.56645 | 0.002872 |
| CG11158 | -0.45104 | 0.002613 | -0.54912 | 0.000101 | -0.33391 | 0.052205 |
| CG11160 | 0.631456 | 0.034183 | 0.776157 | 0.003882 | 0.768668 | 0.005945 |
| CG11164 | -2.02398 | 0.000185 | -1.54027 | 0.003041 | -0.95126 | 0.115499 |
| CG11368 | -1.39144 | 0.017872 | -1.25368 | 0.044568 | -1.074 | 0.094824 |
| CG11403 | -0.75683 | 0.017037 | -1.07475 | 0.000246 | -0.93668 | 0.001042 |
| CG12112 | 0.622269 | 0.00657 | 0.650067 | 0.003676 | 0.787744 | 0.000124 |
| CG12116 | 1.673563 | 0.002988 | 1.386071 | 0.0238 | 2.491572 | 5.45E-07 |
| CG12344 | -0.66854 | 1.44E-06 | -0.39855 | 0.020884 | -0.46382 | 0.002661 |
| CG12861 | -22.8687 | 9.06E-08 | -23.0645 | 7.17E-08 | -6.88626 | 0.284177 |
| CG12994 | 0.425431 | 3.44E-06 | 0.392798 | 4.08E-05 | 0.392977 | 2.46E-05 |
| CG13404 | 0.498814 | 2.37E-11 | 0.235665 | 0.015108 | 0.390675 | 7.19E-07 |
| CG1354 | -0.27946 | 0.024783 | -0.32806 | 0.00486 | -0.26956 | 0.0306 |
| CG1468 | 0.78412 | 2.25E-13 | 0.652154 | 2.91E-09 | 0.842727 | 2.32E-15 |
| CG14683 | -0.17351 | 0.029636 | -0.3052 | 2.04E-06 | -0.14518 | 0.08826 |
| CG14814 | -0.97142 | 1.79E-10 | -0.8051 | 1.21E-06 | -0.93009 | 4.34E-09 |
| CG1503 | -1.37776 | 0.005176 | -1.49536 | 0.001422 | -1.54573 | 0.000556 |
| CG1529 | -0.99051 | 6.54E-14 | -1.19917 | 2.99E-19 | -1.30342 | 7.69E-23 |
| CG15343 | 0.881167 | 0.049811 | 1.505544 | 0.00015 | 0.724057 | 0.163773 |
| CG15365 | -1.47064 | 1.78E-05 | -1.30018 | 0.000158 | -1.5754 | 1.45E-06 |
| CG15445 | 0.455116 | 5.20E-05 | 0.541218 | 2.61E-07 | 0.505655 | 3.23E-06 |
| CG15449 | -1.14289 | 0.000826 | -0.84442 | 0.04435 | -1.58505 | 4.69E-07 |
| CG15478 | 0.256643 | 0.019332 | 0.233992 | 0.047306 | 0.319818 | 0.001088 |
| CG15784 | 1.056993 | 0.0004 | 0.842162 | 0.009149 | 0.839443 | 0.007933 |
| CG15914 | -0.97734 | 0.026678 | -1.27166 | 0.001513 | -0.88484 | 0.049406 |
| CG15916 | 0.626457 | 8.48E-06 | 0.444552 | 0.006508 | 0.63989 | 4.14E-06 |
| CG1637 | -0.24843 | 1.87E-09 | -0.26803 | 6.04E-11 | -0.20828 | 1.45E-06 |
| CG17376 | -6.40442 | 0.011206 | -5.31769 | 0.028323 | -5.55618 | 0.051322 |
| CG17544 | -0.34743 | 1.15E-05 | -0.23375 | 0.012432 | -0.40018 | 1.52E-07 |
| CG17646 | 0.213222 | 0.005572 | 0.299217 | 1.05E-05 | 0.234414 | 0.001333 |
| CG17896 | 0.23085 | 0.000224 | 0.220479 | 0.000595 | 0.232571 | 0.000175 |
| CG1812 | 0.496455 | 8.83E-13 | 0.624129 | 1.80E-20 | 0.637864 | 1.96E-21 |

| | | | | | | |
|---|---|---|---|---|---|---|
| CG18467 | -0.65353 | 2.85E-08 | -0.7176 | 6.83E-10 | -0.58732 | 1.16E-06 |
| CG18508 | 0.624151 | 1.96E-10 | 0.421074 | 0.000129 | 0.635083 | 1.01E-10 |
| CG2003 | 1.967339 | 1.16E-05 | 1.422166 | 0.003563 | 1.742656 | 0.000923 |
| CG2016 | 0.375255 | 4.40E-12 | 0.210723 | 0.001127 | 0.299181 | 1.42E-07 |
| CG2124 | -0.9981 | 9.21E-23 | -0.9265 | 9.54E-19 | -0.81842 | 1.20E-14 |
| CG2135 | -0.44185 | 0.000739 | -0.47117 | 0.000245 | -0.27842 | 0.076236 |
| CG2267 | -6.65014 | 0.047255 | -7.91496 | 0.010161 | -5.83073 | 0.092241 |
| CG2680 | -0.38415 | 9.19E-06 | -0.37602 | 1.52E-05 | -0.3444 | 0.000112 |
| CG2681 | -1.47519 | 3.11E-05 | -1.3793 | 0.000154 | -1.50241 | 2.05E-05 |
| CG2691 | -0.41097 | 1.35E-06 | -0.43709 | 2.04E-07 | -0.49366 | 1.45E-09 |
| CG2889 | -0.52858 | 4.67E-09 | -0.61166 | 5.85E-12 | -0.54813 | 1.50E-09 |
| CG2918 | -0.10341 | 0.022547 | -0.12597 | 0.002489 | -0.06915 | 0.189396 |
| CG3009 | 0.260662 | 0.004449 | 0.254966 | 0.005925 | 0.26921 | 0.002446 |
| CG3011 | 0.613353 | 0.00033 | 0.498596 | 0.008348 | 0.606585 | 0.000417 |
| CG30424 | -0.57713 | 1.07E-05 | -0.57235 | 3.54E-05 | -0.55516 | 5.38E-05 |
| CG31226 | -4.67398 | 0.032586 | -5.77014 | 0.010897 | -4.60221 | 0.056271 |
| CG31709 | -6.64947 | 0.024715 | -7.58721 | 0.006471 | -5.79503 | 0.06062 |
| CG31948 | -7.05421 | 0.022547 | -7.80916 | 0.008241 | -6.31758 | 0.048424 |
| CG31988 | -5.21871 | 0.012978 | -4.78507 | 0.024908 | -3.93363 | 0.082807 |
| CG3213 | -8.46916 | 0.008842 | -7.8938 | 0.019497 | -6.85405 | 0.050056 |
| CG32436 | -2.45065 | 0.034001 | -2.89652 | 0.008329 | -1.81124 | 0.161972 |
| CG32486 | 0.208961 | 3.34E-05 | 0.216634 | 1.55E-05 | 0.150538 | 0.007933 |
| CG32512 | -1.25691 | 6.22E-10 | -0.79292 | 0.000588 | -0.68219 | 0.002577 |
| CG3270 | -0.35812 | 0.001744 | -0.40412 | 0.000223 | -0.39904 | 0.000266 |
| CG32706 | -0.8214 | 0.001307 | -1.24998 | 3.11E-08 | -0.62512 | 0.033197 |
| CG32732 | 0.588224 | 3.72E-12 | 0.856297 | 1.29E-25 | 0.597437 | 9.46E-13 |
| CG32795 | 0.626229 | 4.78E-31 | 0.608771 | 4.99E-29 | 0.697693 | 4.57E-38 |
| CG33340 | -5.83936 | 0.027539 | -6.13349 | 0.018598 | -6.61532 | 0.024835 |
| CG3376 | 0.212021 | 0.014293 | 0.282789 | 0.000305 | 0.214571 | 0.011412 |
| CG3704 | -0.38533 | 0.001526 | -0.34772 | 0.008288 | -0.24493 | 0.100185 |
| CG3706 | -5.52683 | 8.75E-07 | -4.76306 | 7.66E-05 | -7.56559 | 3.60E-07 |
| CG3822 | 0.423165 | 0.000576 | 0.382391 | 0.003037 | 0.366169 | 0.004484 |
| CG4020 | 2.61744 | 1.49E-16 | 3.177022 | 7.21E-22 | 2.810022 | 4.65E-20 |
| CG40470 | -0.23007 | 0.000125 | -0.31875 | 5.73E-09 | -0.31266 | 1.33E-08 |
| CG4050 | 0.535507 | 1.50E-12 | 0.624338 | 6.28E-17 | 0.524774 | 4.24E-12 |
| CG4061 | -0.37289 | 2.30E-05 | -0.44219 | 1.20E-06 | -0.34991 | 0.00015 |

| | | | | | | |
|---|---|---|---|---|---|---|
| CG4078 | -0.37593 | 5.61E-07 | 0.79554 | 1.58E-23 | -0.02076 | 0.878915 |
| CG4096 | 1.275386 | 0.000146 | 1.411645 | 1.62E-05 | 1.009645 | 0.005945 |
| CG4199 | -0.34666 | 0.000819 | -0.34425 | 0.000996 | -0.23388 | 0.055156 |
| CG4250 | 0.781117 | 0.027514 | 1.375341 | 3.88E-07 | 1.286711 | 3.79E-06 |
| CG42709 | -0.35972 | 1.63E-05 | -0.2994 | 0.000934 | -0.30081 | 0.000666 |
| CG4293 | -0.78064 | 5.93E-20 | -0.88297 | 1.28E-24 | -0.74288 | 6.51E-18 |
| CG43673 | 4.956124 | 0.008614 | 5.749011 | 0.004949 | 3.296308 | 0.201172 |
| CG4546 | -3.65715 | 0.024783 | -3.77733 | 0.018197 | -2.02797 | 0.289264 |
| CG4586 | -4.9494 | 1.19E-32 | -4.59938 | 1.59E-31 | -4.72141 | 3.57E-36 |
| CG4615 | 0.249635 | 0.015577 | 0.251702 | 0.016675 | 0.191859 | 0.092351 |
| CG4836 | -7.25729 | 0.000826 | -7.07311 | 0.001327 | -6.57163 | 0.006054 |
| CG4872 | -1.24405 | 1.46E-05 | -1.00668 | 0.000834 | -0.90982 | 0.003718 |
| CG5261 | -0.12368 | 0.046319 | -0.13044 | 0.034737 | -0.2288 | 2.00E-06 |
| CG5404 | -0.31433 | 0.042652 | -0.41507 | 0.002384 | -0.29895 | 0.052844 |
| CG6356 | -0.37877 | 0.031229 | -0.39274 | 0.023735 | -0.14985 | 0.480986 |
| CG6999 | -0.56071 | 0.010112 | -0.68174 | 0.001002 | -0.86231 | 3.30E-06 |
| CG7024 | -1.58371 | 2.75E-12 | -1.41899 | 2.32E-08 | -1.44472 | 6.87E-09 |
| CG7135 | -0.77864 | 1.67E-09 | -0.71779 | 4.80E-08 | -0.92767 | 1.32E-13 |
| CG7149 | -0.18419 | 0.014561 | -0.16928 | 0.04435 | -0.14221 | 0.087316 |
| CG7280 | -0.44405 | 0.007033 | -0.37642 | 0.036034 | -0.44277 | 0.005773 |
| CG7556 | 0.306131 | 0.000653 | 0.29244 | 0.001614 | 0.309025 | 0.000564 |
| CG7607 | 0.624286 | 1.16E-05 | 0.546997 | 0.000255 | 0.663523 | 1.91E-06 |
| CG7692 | -0.29614 | 6.59E-05 | -0.20644 | 0.01664 | -0.16299 | 0.079745 |
| CG8128 | 0.390208 | 4.70E-05 | 0.503456 | 2.45E-08 | 0.493573 | 4.53E-08 |
| CG8289 | -0.3104 | 3.54E-06 | -0.32446 | 7.45E-07 | -0.2368 | 0.001066 |
| CG8300 | 0.326114 | 0.008161 | 0.341318 | 0.004806 | 0.376443 | 0.00106 |
| CG8565 | -5.22748 | 0.016298 | -3.65435 | 0.031913 | -1.95134 | 0.29159 |
| CG8701 | -4.00755 | 0.030918 | -6.02165 | 0.000834 | -2.39856 | 0.26735 |
| CG8939 | -0.77217 | 2.17E-09 | -0.49618 | 0.000757 | -0.53925 | 0.000147 |
| CG9123 | 0.476397 | 0.000562 | 0.371436 | 0.021163 | 0.34624 | 0.0329 |
| CG9164 | -0.18214 | 0.013956 | -0.17892 | 0.017947 | -0.22124 | 0.001027 |
| CG9203 | -0.63463 | 0.000616 | -0.79275 | 3.34E-05 | -0.65977 | 0.00029 |
| CG9314 | -3.98671 | 0.049333 | -4.22433 | 0.038607 | -1.69549 | 0.469973 |
| CG9507 | -0.21657 | 0.014749 | -0.25744 | 0.001861 | -0.26058 | 0.001268 |
| CG9609 | 0.352109 | 0.00033 | 0.255867 | 0.027557 | 0.31454 | 0.002225 |
| CG9629 | -0.28888 | 0.000657 | -0.33264 | 3.67E-05 | -0.1937 | 0.048582 |
| CG9657 | -0.4814 | 3.11E-06 | -0.50998 | 5.36E-07 | -0.34892 | 0.002304 |

| | | | | | | |
|---|---|---|---|---|---|---|
| CG9689 | 0.832069 | 0.000145 | 1.279375 | 5.59E-10 | 1.116232 | 1.23E-07 |
| CG9743 | 0.600587 | 0.000991 | 0.622059 | 0.000604 | 0.44908 | 0.0306 |
| CG9784 | -0.33309 | 1.16E-05 | -0.30926 | 7.66E-05 | -0.38655 | 1.06E-07 |
| CG9975 | -4.57346 | 0.001922 | -3.31651 | 0.023782 | -2.55715 | 0.126362 |
| Cht2 | 5.009455 | 5.26E-39 | 4.854719 | 2.89E-34 | 4.544107 | 1.93E-30 |
| Cp7Fb | 1.342367 | 4.46E-06 | 1.025014 | 0.001905 | 0.784816 | 0.029423 |
| CR17567 | -7.13905 | 0.012936 | -6.57184 | 0.029105 | -4.25673 | 0.194812 |
| CR34335 | 0.658866 | 1.58E-13 | 0.615131 | 1.33E-11 | 0.577229 | 3.10E-10 |
| CR42491 | 0.737159 | 2.30E-11 | 0.783047 | 2.45E-12 | 0.939606 | 3.74E-18 |
| CR43199 | -0.42484 | 0.000712 | -0.42978 | 0.000667 | -0.64709 | 4.34E-09 |
| CR43837 | 1.550885 | 1.85E-06 | 2.359579 | 1.16E-11 | 1.631468 | 1.16E-06 |
| CR43864 | -5.91346 | 0.004174 | -5.56678 | 0.009518 | -2.50242 | NA |
| CR44417 | 0.36381 | 0.038295 | 0.454205 | 0.005957 | 0.399779 | 0.017826 |
| CR44662 | -0.73858 | 0.032709 | -0.82766 | 0.009454 | -0.17048 | 0.705214 |
| CR44841 | -1.7929 | 6.15E-10 | -1.73676 | 3.17E-09 | -1.55224 | 3.29E-07 |
| CR44961 | -3.40399 | 6.27E-15 | -4.46745 | 8.41E-15 | -3.23916 | 1.95E-14 |
| CR45009 | -0.50871 | 0.000197 | -0.58862 | 1.10E-05 | -0.65629 | 3.77E-07 |
| CR45010 | -2.11466 | 8.30E-07 | -1.09044 | 0.01732 | -1.48322 | 0.000374 |
| CR45474 | -4.05374 | 0.017199 | -4.02861 | 0.003156 | -2.60139 | 0.049112 |
| CR45479 | -2.62548 | 1.13E-28 | -2.57724 | 1.48E-27 | -2.38528 | 7.43E-24 |
| CR45528 | 2.201843 | 0.000339 | 1.566723 | 0.024085 | 1.529491 | 0.022323 |
| CR45601 | 2.733779 | 0.000207 | 3.09514 | 9.55E-05 | 2.376358 | 0.001095 |
| CR45629 | 9.42148 | 1.91E-12 | 8.637695 | 2.66E-10 | 9.431543 | 1.83E-12 |
| CR45668 | 2.651887 | 2.93E-14 | 2.253273 | 6.88E-11 | 2.475931 | 1.95E-14 |
| Crg-1 | -2.64454 | 0.003402 | -3.53973 | 0.000897 | -2.73406 | 0.00106 |
| Cyp4ae1 | -0.75664 | 0.000102 | -1.05612 | 2.15E-09 | -0.72885 | 0.000247 |
| Cyp4d2 | -2.3883 | 3.16E-53 | -2.36291 | 5.29E-51 | -2.56325 | 2.37E-61 |
| daw | 0.900268 | 0.001714 | 0.705544 | 0.035892 | 0.599788 | 0.082807 |
| Dh44 | 0.216445 | 0.038617 | 0.245898 | 0.012788 | 0.308777 | 0.000472 |
| dl | 0.952005 | 5.83E-06 | 0.953279 | 2.59E-06 | 0.761301 | 0.000532 |
| Dok | -1.2546 | 7.28E-11 | -1.13502 | 2.26E-09 | -1.17539 | 1.52E-09 |
| Drep-1 | 0.231556 | 0.003741 | 0.216507 | 0.009362 | 0.139613 | 0.148514 |
| drm | -0.71486 | 0.035524 | -0.83278 | 0.00936 | -0.49256 | 0.20711 |

| | | | | | | |
|---|---|---|---|---|---|---|
| dsx | -1.07573 | 3.58E-06 | -0.88368 | 0.000353 | -0.75375 | 0.003969 |
| EfTuM | -0.20022 | 0.018489 | -0.24845 | 0.001455 | -0.19492 | 0.022064 |
| eIF3-S8 | -0.19223 | 5.37E-05 | -0.16029 | 0.001837 | -0.15393 | 0.002855 |
| Es2 | 0.482862 | 4.49E-06 | 0.378837 | 0.000975 | 0.283091 | 0.029048 |
| ewg | 0.192171 | 0.047688 | 0.334966 | 1.25E-05 | 0.283809 | 0.000412 |
| fit | 2.851146 | 0.000737 | 3.35968 | 9.91E-05 | 3.846685 | 7.95E-06 |
| Fmo-2 | 1.189099 | 0.004113 | 1.271031 | 0.005026 | 1.152952 | 0.006653 |
| fs(1)Yb | -4.47982 | 0.000136 | -2.66857 | 0.000398 | -2.20338 | 0.002868 |
| Fuca | -0.32384 | 0.026639 | -0.32525 | 0.033456 | -0.40354 | 0.001978 |
| Gel | 0.158043 | 0.018029 | 0.168465 | 0.009136 | 0.215287 | 0.00019 |
| Gga | -0.2633 | 0.010827 | -0.31006 | 0.001704 | -0.29507 | 0.002817 |
| GlcAT-I | 0.734002 | 7.29E-18 | 0.421581 | 1.25E-05 | 0.584078 | 4.24E-11 |
| Glt | 0.949105 | 0.03456 | 1.100579 | 0.009136 | 1.464681 | 9.04E-05 |
| Gs2 | -0.22542 | 0.006708 | -0.33334 | 3.30E-06 | -0.17338 | 0.057156 |
| GstS1 | -0.2867 | 0.008969 | -0.26628 | 0.020119 | -0.22615 | 0.058235 |
| hec | -0.76821 | 1.46E-13 | -0.39842 | 0.001813 | -0.65157 | 3.77E-09 |
| Hexo2 | 0.234603 | 0.000453 | 0.377127 | 2.43E-10 | 0.300799 | 1.16E-06 |
| His3.3B | 0.356032 | 4.75E-06 | 0.227188 | 0.015332 | 0.293678 | 0.000374 |
| Hsp60C | -4.70963 | 0.009795 | -4.94268 | 0.00294 | -3.15117 | 0.116665 |
| Hug | 0.789142 | 2.10E-06 | 0.500289 | 0.011829 | 0.618821 | 0.000541 |
| Ilp3 | 1.82441 | 1.03E-32 | 1.304435 | 3.54E-16 | 1.278484 | 2.32E-15 |
| Ilp6 | -0.5689 | 0.002245 | -0.46119 | 0.027915 | -0.37184 | 0.094158 |
| Inx7 | 1.81171 | 1.91E-11 | 2.029181 | 1.14E-12 | 1.564227 | 1.65E-07 |
| jigr1 | 0.181196 | 0.02486 | 0.16707 | 0.049012 | 0.209845 | 0.004603 |
| Karl | 1.237324 | 0.020499 | 1.996738 | 2.62E-06 | 1.024953 | 0.052885 |
| l(1)1Bi | -0.51412 | 0.00162 | -0.41858 | 0.020242 | -0.50579 | 0.002082 |
| l(1)G0020 | -0.77047 | 0.000273 | -0.79008 | 0.000288 | -0.61667 | 0.008477 |
| l(1)G0045 | -1.56217 | 0.000207 | -1.94471 | 9.42E-07 | -1.48708 | 0.000481 |
| l(1)G0230 | 0.331939 | 7.70E-07 | 0.232179 | 0.00248 | 0.362827 | 3.53E-08 |
| l(1)G0320 | 0.243671 | 2.11E-05 | 0.243021 | 2.58E-05 | 0.292514 | 7.50E-08 |
| l(2)05714 | 0.588109 | 0.000129 | 0.539185 | 0.000612 | 0.506518 | 0.001634 |

| | | | | | | |
|---|---|---|---|---|---|---|
| lawc | 0.546837 | 3.62E-05 | 0.485332 | 0.000519 | 0.598509 | 3.60E-06 |
| Lint-1 | 0.320246 | 0.001714 | 0.416385 | 1.25E-05 | 0.348468 | 0.000452 |
| Lkr | 0.40996 | 0.027528 | 0.482333 | 0.003563 | 0.403747 | 0.024876 |
| LM408 | -0.40157 | 4.33E-12 | -0.35415 | 2.68E-09 | -0.28785 | 4.20E-06 |
| Lon | -0.13414 | 0.033755 | -0.13047 | 0.046778 | -0.17415 | 0.001816 |
| loopin-1 | -5.0858 | 0.007404 | -4.45632 | 0.024825 | -2.92497 | 0.199882 |
| mab-21 | 0.34431 | 0.005825 | 0.43207 | 0.000214 | 0.393881 | 0.00091 |
| MAPk-Ak2 | 0.258629 | 7.01E-07 | 0.221185 | 6.32E-05 | 0.172289 | 0.003956 |
| Mapmodulin | -0.22843 | 2.98E-05 | -0.27723 | 9.84E-08 | -0.29114 | 1.32E-08 |
| Mcm3 | -0.6787 | 0.027855 | -1.61675 | 2.43E-10 | -1.48543 | 1.31E-09 |
| Mdr65 | 0.319084 | 9.75E-06 | 0.333118 | 2.85E-06 | 0.326942 | 4.15E-06 |
| mew | -0.76191 | 1.91E-23 | -0.31823 | 0.00036 | -0.58768 | 2.62E-14 |
| mfas | -0.3237 | 1.99E-06 | -0.30587 | 1.11E-05 | -0.24386 | 0.001042 |
| mod(r) | 0.348259 | 0.005742 | 0.333743 | 0.009533 | 0.362379 | 0.002997 |
| mRpL14 | -0.38306 | 0.007146 | -0.32044 | 0.041901 | -0.34443 | 0.01902 |
| mRpL16 | -0.30547 | 0.01817 | -0.39164 | 0.000834 | -0.35544 | 0.0031 |
| mRpS25 | -0.37056 | 0.015227 | -0.51255 | 0.000154 | -0.32093 | 0.047253 |
| msl-2 | -0.50973 | 3.74E-14 | -0.30449 | 6.22E-05 | -0.43575 | 2.88E-10 |
| Mst87F | -4.66011 | 0.010454 | -3.92623 | 0.027896 | -2.52475 | 0.218815 |
| mthl1 | -0.45295 | 0.003776 | -0.44534 | 0.004969 | -0.57761 | 4.38E-05 |
| Myo28B1 | -0.62472 | 2.11E-05 | -0.81991 | 5.37E-09 | -0.38574 | 0.022866 |
| nAChRbeta3 | 1.215622 | 0.021522 | 1.435341 | 0.001551 | 0.833155 | 0.134292 |
| Ndc80 | -2.19295 | 0.007383 | -1.94677 | 0.023667 | -0.91539 | 0.352979 |
| ninaB | -0.28622 | 0.008111 | -0.31959 | 0.00199 | -0.14539 | 0.280082 |
| NnaD | -0.61728 | 9.35E-08 | -0.43556 | 0.000754 | -0.55882 | 2.10E-06 |
| nod | -3.19342 | 2.87E-29 | -3.47462 | 5.46E-28 | -3.06461 | 7.29E-27 |
| Nup205 | -0.49708 | 0.006889 | -0.42966 | 0.030877 | -0.58224 | 0.000654 |
| Obp18a | 1.318056 | 1.24E-19 | 1.071054 | 2.76E-12 | 1.080089 | 5.27E-13 |
| Obp44a | -0.37253 | 0.000174 | -0.56373 | 2.43E-10 | -0.31722 | 0.002473 |
| Obp99a | 2.291537 | 0.005378 | 2.430783 | 0.003676 | 3.230713 | 2.68E-05 |
| Obp99b | -4.49689 | 5.30E-13 | -3.11039 | 1.01E-08 | -5.2259 | 3.34E-06 |
| obst-A | -1.60326 | 1.21E-08 | -2.3374 | 2.37E-14 | -2.01292 | 1.32E-12 |
| Orct | -0.2675 | 0.022834 | -0.29655 | 0.008338 | -0.29408 | 0.008136 |
| P5CDh1 | -0.14442 | 0.008514 | -0.18541 | 0.000183 | -0.11182 | 0.063804 |
| Pi3K21B | -0.36832 | 0.04908 | -0.38329 | 0.038261 | -0.32451 | 0.08924 |
| Plod | 0.237347 | 0.035211 | 0.362106 | 0.00014 | 0.315317 | 0.001321 |
| Pmp70 | -0.28736 | 0.010393 | -0.2809 | 0.015085 | -0.36719 | 0.000282 |

| | | | | | | |
|---|---|---|---|---|---|---|
| png | -3.43235 | 0.00089 | -2.89246 | 0.002066 | -2.19287 | 0.025474 |
| PPP4R2r | -0.15712 | 0.025683 | -0.18739 | 0.004253 | -0.11127 | 0.163228 |
| Proc | 1.021715 | 1.19E-32 | 0.535833 | 1.37E-08 | 0.654825 | 5.10E-13 |
| pyd3 | 0.609194 | 0.002613 | 0.590221 | 0.003826 | 0.712075 | 0.000157 |
| Rcd-1 | 0.360322 | 1.11E-07 | 0.317798 | 8.59E-06 | 0.276015 | 0.000184 |
| regucalcin | 0.366973 | 0.028683 | 0.555735 | 7.73E-05 | 0.518655 | 0.000281 |
| RhoGAP102A | -0.90392 | 0.006431 | -0.77445 | 0.031128 | -1.13056 | 0.000167 |
| RhoGAP1A | 0.3295 | 0.001636 | 0.488749 | 1.15E-07 | 0.433245 | 4.23E-06 |
| roX1 | -8.72114 | 0 | -8.18519 | 0 | -8.42937 | 0 |
| roX2 | -8.37253 | 3.21E-177 | -7.71497 | 2.46E-198 | -8.38122 | 6.05E-158 |
| S-Lap2 | -8.13746 | 0.000363 | -7.83974 | 0.000754 | -4.90564 | 0.037117 |
| S-Lap4 | -6.6938 | 0.010426 | -6.54421 | 0.009533 | -3.5262 | 0.229746 |
| S-Lap8 | -7.33945 | 0.009963 | -7.84767 | 0.00476 | -7.352 | 0.008916 |
| Seipin | -0.83567 | 0.003026 | -0.81147 | 0.00429 | -0.86674 | 0.001717 |
| SIP2 | -5.80833 | 0.00231 | -5.84604 | 0.001271 | -4.86851 | 0.012843 |
| slim | 0.215663 | 0.009047 | 0.226421 | 0.005043 | 0.203497 | 0.014396 |
| sn | -0.64741 | 2.15E-05 | -0.45072 | 0.010897 | -0.77484 | 9.61E-08 |
| sofe | -1.14056 | 0.015577 | -1.2163 | 0.013933 | -1.00236 | 0.033851 |
| spi | 0.168965 | 0.002772 | 0.1538 | 0.009287 | 0.139481 | 0.02121 |
| Ssu72 | 0.314491 | 0.022834 | 0.39764 | 0.001704 | 0.151142 | 0.356799 |
| ssx | 0.713849 | 8.83E-13 | -0.30838 | 0.018843 | 0.586335 | 1.70E-08 |
| ST6Gal | 0.28929 | 1.07E-05 | 0.380929 | 9.66E-10 | 0.351713 | 1.80E-08 |
| stas | 0.266587 | 2.43E-10 | 0.179487 | 0.000176 | 0.262838 | 1.00E-09 |
| temp | 0.415765 | 5.72E-12 | 0.417265 | 1.33E-11 | 0.398796 | 7.26E-11 |
| Tig | 1.29522 | 0.005478 | 1.46287 | 0.000426 | 0.827151 | 0.121037 |
| Treh | 0.301099 | 0.007757 | 0.357897 | 0.000698 | 0.358057 | 0.000574 |
| TrxT | -4.81433 | 0.004803 | -6.3213 | 0.000281 | -4.22483 | 0.017819 |
| tyn | -0.94419 | 3.24E-29 | -0.62479 | 1.05E-12 | -1.00927 | 6.33E-33 |
| xmas-2 | -0.51045 | 2.11E-05 | -0.39261 | 0.003037 | -0.43494 | 0.000583 |
| XRCC1 | -0.49851 | 0.022454 | -0.48021 | 0.032131 | -0.38281 | 0.103922 |
| y | 1.831981 | 1.18E-05 | 1.673541 | 0.000488 | 2.150552 | 4.23E-06 |
| yellow-h | -0.86498 | 0.046478 | -0.76753 | 0.046273 | -0.49132 | 0.288193 |

| | | | | | | |
|---|---|---|---|---|---|---|
| Yp1 | 5.766279 | 4.75E-06 | 6.430996 | 1.66E-07 | 7.017535 | 4.87E-09 |
| Yp2 | 5.473473 | 8.39E-11 | 5.700106 | 1.81E-11 | 6.739718 | 3.65E-16 |
| Yp3 | 6.042375 | 1.17E-12 | 6.196801 | 5.38E-13 | 6.67014 | 2.78E-15 |
